# The role of the gut microbiota in patients with Kleefstra syndrome

**DOI:** 10.1101/2022.02.04.478662

**Authors:** Mirjam Bloemendaal, Priscilla Vlaming, Anneke de Boer, Karlijn Vermeulen-Kalk, Arianne Bouman, Tjitske Kleefstra, Alejandro Arias Vasquez

**Author notes:** These authors contributed equally (Mirjam Bloemendaal, Priscilla Vlaming).

## Abstract

Kleefstra Syndrome (KS) is a rare monogenetic syndrome, caused by haploinsufficiency of the EHMT1 gene, an important regulator of neurodevelopment. The clinical features of KS include intellectual disability, autistic behavior and gastrointestinal problems. The gut microbiota may constitute a, yet unexplored, mechanism underlying clinical variation, as they are an important modifier of the gut-brain-axis. To test whether variation in the gut microbiota is part of KS, we investigated the gut microbiota composition of 23 individuals with KS (patients) and 40 of their family members. Both alpha and beta diversity of patients were different from their family members. Genus *Coprococcus* 3 was lower in abundance in patients compared to family members. Moreover, abundance of genus *Merdibacter* was lower in patients versus family members, but only in the participants reporting intestinal complaints. Within the patient group, behavioral problems explained 7% variance in the beta diversity. Also, within this group, we detected higher levels of *Coprococcus* 3 and *Atopobiaceae – uncultured* associated with higher symptoms severity. Our results show significant differences in the gut microbiota composition of patients with KS compared to their family members, suggesting that these differences are part of the KS phenotype.

## Introduction

Kleefstra Syndrome (KS) is a rare genetic neurodevelopmental syndrome, caused by haploinsufficiency (microdeletion or an intragenic mutation) of the euchromatic histone methyl transferase 1 (*EHMT1*) gene (Kleefstra et al., 2006). Since the identification of the syndrome in the early 1990s, over 100 individuals with KS, from here on referred to as patients, have been described (Willemsen et al., 2012). Clinically, KS is characterized by intellectual disability and developmental delay, typical facial appearance and hypotonia in childhood (Kleefstra et al., 2006, 2009; Stewart & Kleefstra, 2007; Willemsen et al., 2012). Co-morbid autism spectrum disorders (ASDs) are highly prevalent in this syndrome (Vermeulen, de Boer, et al., 2017). Over the lifespan of these children, a major concern is the occurrence of regression during adolescence and young adulthood. This is often thought to be due to a (manic) psychotic disorder, causing loss of functioning, disorganization and sleep problems. These episodes of regression typically start during puberty or young adulthood (Vermeulen, Staal, et al., 2017). Besides the effect on neurodevelopment, many patients or their guardians (such as family members) report gastrointestinal problems co-morbid with KS, such as severe constipation which, in some cases, requires surgery. Personal communication from Kleefstra et al. (unpublished data) indicates digestive problems in 59.6% (56/94) of patients, of which 21 report severe and 18 mild constipation. Repeated respiratory infections at pediatric ages are also prevalent (Kleefstra et al., 2009). Hence, both mental (intellectual disability, ASD) as well as somatic disorders (gastrointestinal and proneness to infections) are part of the KS phenotype. Treatment options are limited to medication aimed at reducing problematic behavior and relieving somatic complaints. The genotype-phenotype correlation in KS is low as there is still high (unexplained) variability in functioning between patients with a similar size or variant of the EHMT1 haploinsufficiency.

An unexplored mechanism underlying the mental and somatic symptoms in KS and explaining part of the variability in phenotype may be impairments in the bi-directional communication that exists between the gastrointestinal tract and the central nervous system (CNS), called the gut-brain-axis (GBA) (Cryan & Dinan, 2012). The GBA encompasses several routes, including endocrine signaling through hormones and neuro-active metabolites as well as signaling through the immune system and vagal nerve. The trillion of bacteria commensally living in our gastrointestinal system (also known as the gut microbiota) are an essential hub of the GBA, that modulate the bidirectional communication routes with the CNS (Cryan et al., 2019). Directly after birth and during early life, the gut microbiota play a crucial role in building a healthy gut, educating immune processes and shaping neurodevelopment (Borre et al., 2014; Sordillo et al., 2019).

GBA alterations and differences in gut microbial composition (compared to family members or healthy control subjects) are observed in many neurodevelopmental and intellectual disability disorders (NDDs and IDDs) with similar symptoms to those also observed in KS (Dam et al., 2019; Q. Li, Han, Dy, & Hagerman, 2017; Van Ameringen et al., 2019). As there are, to our knowledge, hardly any studies reporting on the gut microbiota in homogeneous monogenetic (or Mendelian) syndromes such as KS, these disorders are the only relevant reference we currently have. Of these, cohorts of patients with ASDs are particularly relevant for KS and have been quite extensively studied, at least in comparison to other NDDs. Similar to KS, gastrointestinal problems occur more frequently in patients with ASDs compared to family members and the general population (Holingue, Newill, Lee, Pasricha, & Daniele Fallin, 2018) and bowel problem severity correlates with ASD-symptoms severity in a study by Adams, Johansen, Powell, Quig, & Rubin (2011). Most studies investigating the gut microbiota in ASD show less diverse microbiota in the ASD group compared to controls, as reviewed by Bundgaard-Nielsen et al. (2020). The most often observed differences include an increased abundance of bacterial genera *Bacteriodes* and *Clostridium* as well as a decreased abundance of *Prevotella, Bifidobacterium and Streptococcus* (Bundgaard-Nielsen et al., 2020).

A few (smaller) studies exist investigating the role the gut microbiota play in monogenetic IDDs, such as Down Syndrome (trisomy 21) and Rett syndrome, which are also characterized by similar symptoms as seen in KS such as ASD behaviors and gastrointestinal problems (Bermudez et al., 2019; Smeets, Pelc, & Dan, 2012). In Down Syndrome, increased relative abundance of (amongst others) genus *Sutterella* was observed, positively correlating with the Aberrant Behavior Checklist score (Biagi et al., 2014). In Rett syndrome both alpha and beta diversity were altered and several genera showed increased abundance (Borghi et al., 2017; Strati et al., 2016).

These above-mentioned findings in disorders similar and/or co-morbid with KS suggest that KS may similarly be characterized by alterations in gut microbial composition. These alterations may contribute to gastrointestinal symptoms but also immune functioning and behavioral impairments. Hence, we hypothesize that the gut microbiota is an integral component of the clinical development and presentation of patients with KS.

In this exploratory study we aimed to identify gut microbial variation associated with the clinical presentation of KS. We were able to collect data from 23 patients diagnosed with KS and 40 of their family member (n=14 fathers, n=16 mothers and n=10 siblings). To this aim, we assessed differences in global gut microbial community (alpha and beta diversity) as well as microbial composition at the taxonomic genus level, and compare them between patients with KS and their family members. Moreover, we tested the association between the abundance of bacterial genera with intestinal complaints (in both patients and family members), the EHMT1 genotype (of the patients) and two instruments measuring behavioral problems in the patients; the Autism Diagnostic Observation Schedule (ADOS) and Child Behavior CheckList (CBCL).

## Methods

### Participants

In total 63 subjects participated in this study: 23 patients with KS and 40 of their family members consisting of 16 mothers, 14 fathers and 10 siblings. (See **Table 1** for their descriptive statistics). As shown in table 1, our sample includes both adults and children (both patients and family members). This age difference limits our ability to compare Body Mass Index values (Cole, Bellizzi, Flegal, & Dietz, 2000). Therefore, we first grouped our participants by age and then grouped each subjects’ BMI into their age and gender appropriate category (healthy, overweight or obese) (Kist-van Holthe et al., 2012). Analyses were done using these categories rather than crude BMI values. For seven samples (one patient, one father and five siblings) BMI was imputed using the median of their family group used in the analyses (patient or family members), see **Table S1** for samples with missing values. The variable “dietary setting” was used to indicate shared or separate dietary intake between patients and family members. For example, when patients live at home they likely have a largely similar dietary intake across that whole family. In case of a patient living in a community or a sibling or parent living in a different household (e.g. in case of divorced parents) we assumed no shared dietary intake. No information about dietary setting (of family members) was available for two siblings so we grouped them as living at home. We used the “dietary setting” variable as a covariate in our models (coded as “living at home” and “not living at home”). The variable “familyID” was used to correct for family relatedness in all analyses when including all subjects. As there is a large imbalance in age between the patient and family members, we corrected for age as well as non-linear effects of age (age^2^) in our analyses, see the section Statistical Analysis.

**Table 1.** Descriptive statistics of patients with Kleefstra syndrome and their family members.

| Variable (options) | Patients | Fathers | Mothers | Siblings |
| --- | --- | --- | --- | --- |
| Sample size (n) | 23 | 14 | 16 | 10 |
| Gender (% male) | 35 | 100 | 0 | 30 |
| Age (mean & range) | 18.8 (4.6 – 39.8) | 46.8 (26.6 – 64.0) | 42.9 (26.8 – 66.2) | 12.8 (3.7 – 23.3) |
| **BMI** |  |  |  |  |
| Healthy | 8 | 7 | 7 | 9 |
| Overweight | 12 | 6 | 7 | 1 |
| Obese | 3 | 1 | 2 | 0 |
| **Dietary Setting** |  |  |  |  |
| At home | 16 | 13 | 16 | 10 |
| Not at home | 7 | 1* | 0 | 0 |
| **ADOS comparative score** mean (range) | 6.7 (5-10) | NA | NA | NA |
| **CBCL total problems score** mean (range) | 68.4 (50-85) | NA | NA | NA |
| **Genetic variants** |  |  |  |  |
| Deletion <1MB | 6 | NA | NA | NA |
| Deletion >1MB | 4 | NA | NA | NA |
| Pathogenic mutation | 12 | NA | NA | NA |

\* divorced, co-parenting.

To further characterize the patient sample and the relation among their phenotypic features, being behavioural and somatic symptoms and their genetic variants, correlations were performed between these 4 measures.

### Experimental procedure

All known families of patients with KS at our expert center (EC) at time of initiation of the study were approached for participation (via the Radboudumc EC for rare genetic neurodevelopmental syndromes). Patients were included if they had a molecular proven diagnoses of KS with a genetic description and if they had a biological age of at least three years old. The three year old cut-off was used as the behaviour problems are not yet classified. Moreover, from that age children eat food similar to their parents, making their gut microbiota more comparable. The involved medical specialist informed them about the study and the families received an information letter. Written informed consent was obtained by the (adult) family members and/or their legal guardian. After the written informed consent was signed, the families received questionnaires to complete as well as the material for fecal collection. Data were collected between July and November 2017.

### Instruments

#### Autism diagnostic observation schedule (ADOS)

The Autism diagnostic observation schedule (ADOS) (C Lord et al., 2012) is a standardised diagnostic instrument, which is frequently used worldwide (Catherine Lord, Elsabbagh, Baird, & Veenstra-Vanderweele, 2018) to assess autism features. It provides a semi-structured observation, which is performed by a certified psychologist or psychiatrist and consists of four modules based on the language capacity of the participant. It runs from module 1, pre-verbal to minimal verbal capacity and includes activities, to module 4, fluently verbal and includes adult topics to discuss. A comparison score is available to compare the different modules. The score for clinical suspicion of an autism spectrum disorder ranges between 5 - 10 (moderate to severe suspicion) (Vermeulen, de Boer, et al., 2017).

Our previous studies in KS patients showed a high prevalence of ASD, in which the ADOS-2 instrument best discriminated the ASD-symptoms from the developmental delay (Vermeulen, Egger, et al., 2017). All patients in this study scored 5 or higher, see **Table 1**.

#### Child Behavior CheckList (CBCL)

The Child Behavior CheckList (CBCL) is a widely used diagnostic instrument to quantify problem behaviour and skills of children and adolescents, completed by the parents or guardian (T.M. Achenbach & Rescorla, 2001; Thomas M Achenbach, Dumenci, & Rescorla, 2001). The version for children from 1,5 years to 5 years old was used, corresponding to the developmental age of the patients. Parents or guardians answer 99 questions about emotional and behavioural problems they observe in the child, answers ranging from 0 “not true” to 2 “very true”. A total t-score (for a broad range of problematic behaviour, internalising as well as externalising) was calculated and used in this study. In patients with NDDs, like KS, the total t-score is the best reflection of behavioural problems, because there is often a mix of ID, comorbid NDDs (like ASD) and they are prone to develop major psychiatric disorders as well, like psychosis or mood disorders (Vermeulen, de Boer, et al., 2017). A total score of 59 or below indicates non-clinical symptoms, a score between 60-64 indicates the child is at risk for problem behaviours and a score of 65 or higher indicates clinical symptoms (T. Achenbach & Rescorla, 2001). Three patients scored below the non-clinical symptom cut-off score, 5 were in the at risk category and all others (n=15) scored within the clinical symptom range.

#### Lifestyle questionnaire – Intestinal complaints

A semiquantitative lifestyle questionnaire was used to assess dietary habits, medication, smoking, allergies, dietary setting, incontinence, physical and mental health. For dietary habits, habitual intake frequency of 9 core food and beverage groups was assessed (processed meat, fish, fruit, legumes, vegetable, milk, sweetened beverages, chocolate and alcohol). For processed meat, fish, fruit, legumes, vegetable and sweetened beverages the 4 response options range are “never”, “monthly”, “weekly” or “everyday”. The consumption of alcohol, chocolate and milk was quantified in terms of daily intake; alcohol at weekdays and weekends was indicated in terms of 0, 1-2, 3-4, 5-6 or >6 glasses per day. Consumption of chocolate was listed as 0-1, 1-5, 5-10 or >10 bars of 200 grams per month and milk as no, 1-2, 2-4, >4 glasses of 200 ml milk per day.

Besides diet, a range of physical and mental health questions was included in this lifestyle questionnaire, for an overview see **Table S2**. Of specific clinical interest are “intestinal complaints” which were indicated for patients and family members on a 7 point scale ranging from “never”,” < 1 times per month”, >1 times per month”, “2-3 times per month”, “1 day per week”, “>1 day per week” or “every day”. As these 7 points results in very small sample sizes per category we coded these into two groups: “No”, containing the “never” group and “Yes”, containing the other groups. Detailed reported medication prescriptions by patients and family members are listed separately in **Table S3**.

#### Genetic variants

Loss of function in *EHMT1* gene accounts for most features in Kleefstra syndrome, which can be caused by several genetic variants. We performed two analyses on the effect of genetic variants on gut microbial community measures (see **Table 2** for an overview on the analyses): first, using two groups of genetic variants: pathogenic mutations + deletions < 1MB and deletions > 1MB and second, using three dissociable groups: pathogenic mutation, small deletion < 1Mb) and large deletion (> 1Mb). The patients with pathogenic mutations or small deletions have a similar clinical severity spectrum (Kleefstra & de Leeuw, 2019). However, patients with larger deletions usually have more severe intellectual disability and more severe medical/somatic problems such as congenital anomalies, feeding problems and respiratory problems (Kleefstra & de Leeuw, 2019).

**Table 2.**
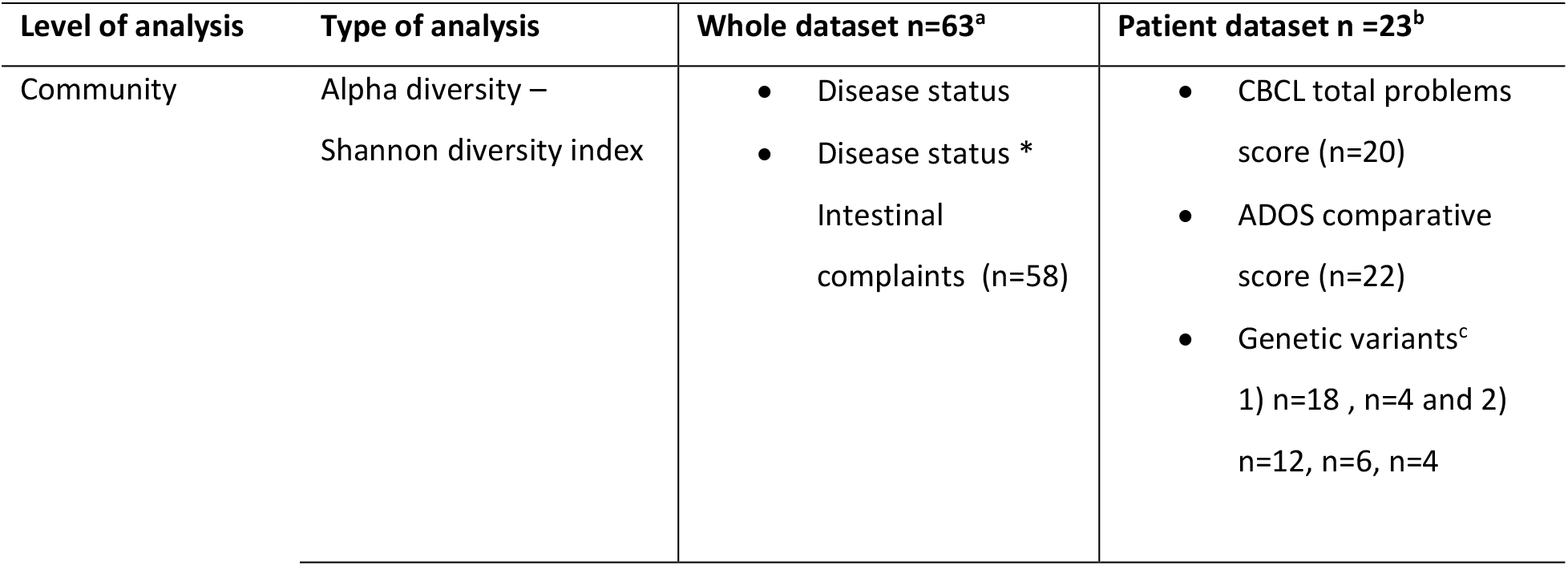
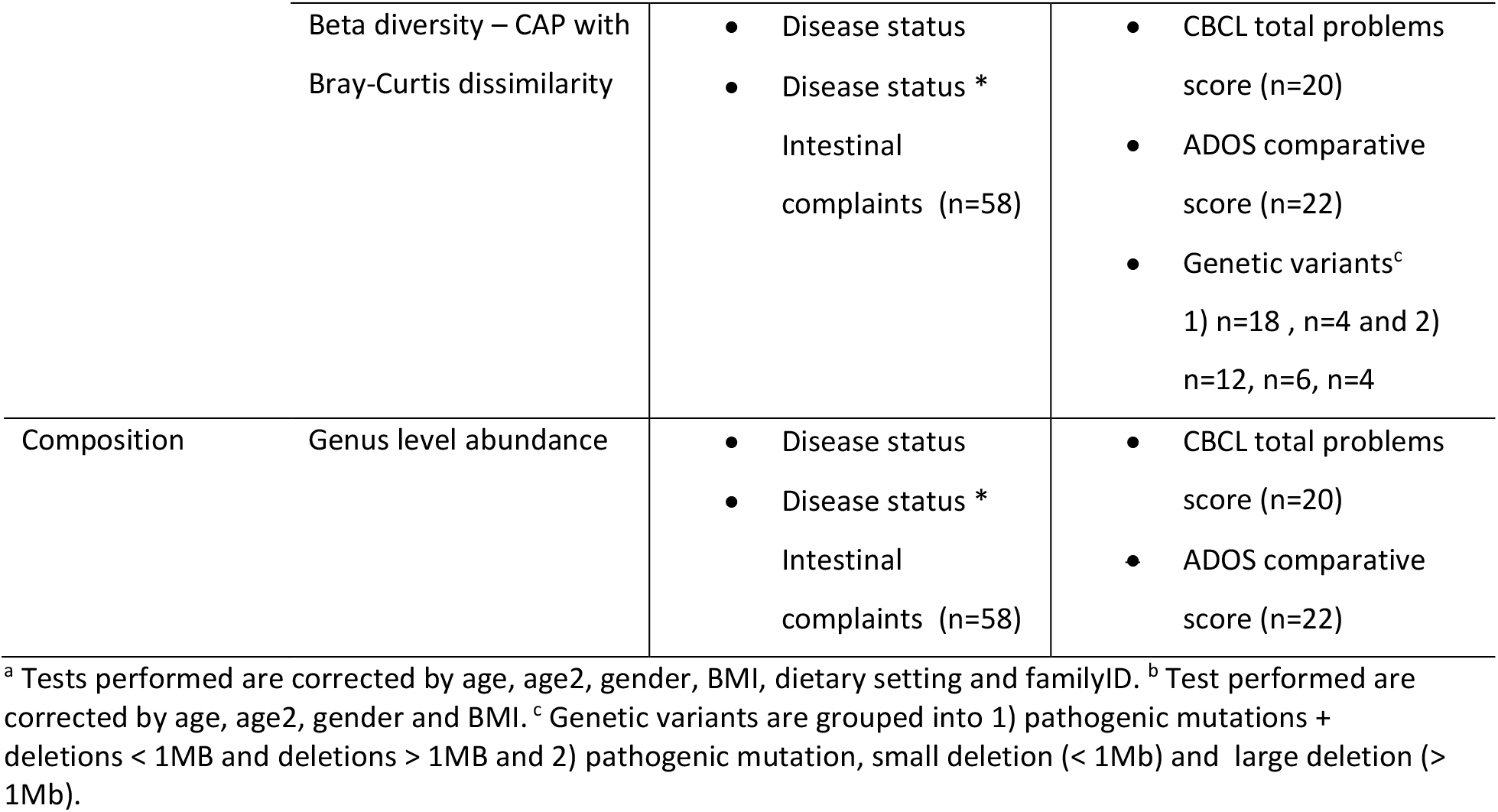
Schematic of the analyses performed in the study.

| Level of analysis | Type of analysis | Whole dataset n=63<sup>a</sup> | Patient dataset n =23<sup>b</sup> |
| --- | --- | --- | --- |
| Community | Alpha diversity –<br>Shannon diversity index | Disease status<br>Disease status *<br>Intestinal complaints (n=58) | CBCL total problems score (n=20)<br>ADOS comparative score (n=22)<br>Genetic variants<sup>c</sup><br>1) n=18 , n=4 and 2)<br>n=12, n=6, n=4 |
|  | Beta diversity – CAP with<br>Bray-Curtis dissimilarity | • Disease status<br>• Disease status *<br>Intestinal<br>complaints (n=58) | • CBCL total problems<br>score (n=20)<br>• ADOS comparative<br>score (n=22)<br>• Genetic variants<sup>c</sup><br>1) n=18 , n=4 and 2)<br>n=12, n=6, n=4 |
| Composition | Genus level abundance | • Disease status<br>• Disease status *<br>Intestinal<br>complaints (n=58) | • CBCL total problems<br>score (n=20)<br>• ADOS comparative<br>score (n=22) |

<sup>a</sup> Tests performed are corrected by age, age<sup>2</sup>, gender, BMI, dietary setting and familyID. <sup>b</sup> Test performed are corrected by age, age<sup>2</sup>, gender and BMI. <sup>c</sup> Genetic variants are grouped into 1) pathogenic mutations + deletions < 1MB and deletions > 1MB and 2) pathogenic mutation, small deletion (< 1Mb) and large deletion (> 1Mb).

### Microbiota procedures

#### Fecal sample collection

Fecal samples were collected by the subjects at home using a validated protocol by OMNIgene•GUT kit (DNAGenotek, Ottawa, CA). The material was aliquoted into 1.5 ml Eppendorf tubes and stored in −80°C until further laboratory processing (Szopinska et al., 2018).

#### Bacterial DNA isolation and sequencing

For bacterial DNA extraction, 150ul or mg of feces was aliquoted from the Eppendorf tube. Microbial DNA was isolated including a bead-beating procedure and purified using a customized protocol by Baseclear B.V. (Leiden, the Netherlands). The V4 region of 16S ribosomal RNA (rRNA) gene was amplified in duplicate PCR reactions for each sample. The V4 region was targeted using primers 515F (5’GTGYCAGCMGCCGCGGTAA) and 806R (5’ GGACTACNVGGGTWTCTAAT). Both a PCR negative was included sample to assess contamination introduced during the PCR step as well as two positive control samples to assess correct detection of a sample with a known microbial distribution. Libraries were pooled for sequencing on the Illumina NovaSeq 6000 platform (paired-end, 250 bp) (Baseclear B.V., Leiden, the Netherlands). Sequence reads were demultiplexed and filtered using Illumina Chastity. Reads containing PhiX control signal were removed and adapter sequences were removed from all reads. An average sequencing quality score “Q” of 35,98 (range 35,17 – 36,25) was obtained indicating the sequencing error probability for a given base, where Q=30 indicates 99,9% accuracy. We observed a total number of raw read pairs of 57.749.319 with an average of 916.655 and a range of 752.140 - 1.209.564.

#### Bioinformatics

Reads were processed for statistical analysis in QIIME2. First, reads were denoised using DADA2 (Callahan et al., 2016): primers were trimmed off, sequencing errors (chimeras) were removed, combined read pairs and erroneous read combinations were removed. Read pairs were aligned into Amplicon Sequence Variants (ASVs) using mafft (Katoh, Misawa, Kuma, & Miyata, 2002). Combined reads deviating from the expected combined length of 250 bases were filtered out, allowing variation of 10 bases both directions (i.e. 240-260 range), which was 0.1% of the total of 39.777.830 reads. This resulted in a total of 39.739.397 reads corresponding to 5452 unique ASVs over the 63 samples. An average read count per sample of 635.858 was obtained, ranging between 479.606 – 896.730, see **Figure 1**. These sequences represented a total of 6.771 ASVs with an average of 477 ASVs per sample, ranging between 176-914 ASVs. Taxonomic assignment was done in q2-feature-classifier using the Naive Bayes classifier (Bokulich et al., 2018) trained on the SILVA reference database (version 132 with 99% OTU from 515/806R region of sequences) (Yarza et al., 2014). A phylogenetic tree was built using fasttree2 (Price, Dehal, & Arkin, 2010) and combined in a biom file including ASV and taxonomy information. Non-bacterial and unassigned ASVs were removed from the biom file. Data was made compositional; i.e. abundance of each ASV was expressed relative to the total amount and prevalence of ASVs. All statistical analyses were performed in R (version 3.6.2) using packages phyloseq (McMurdie & Holmes, 2013) and microbiome (Lathi & Shetty, 2017), other packages are cited when used.

**Figure 1.**
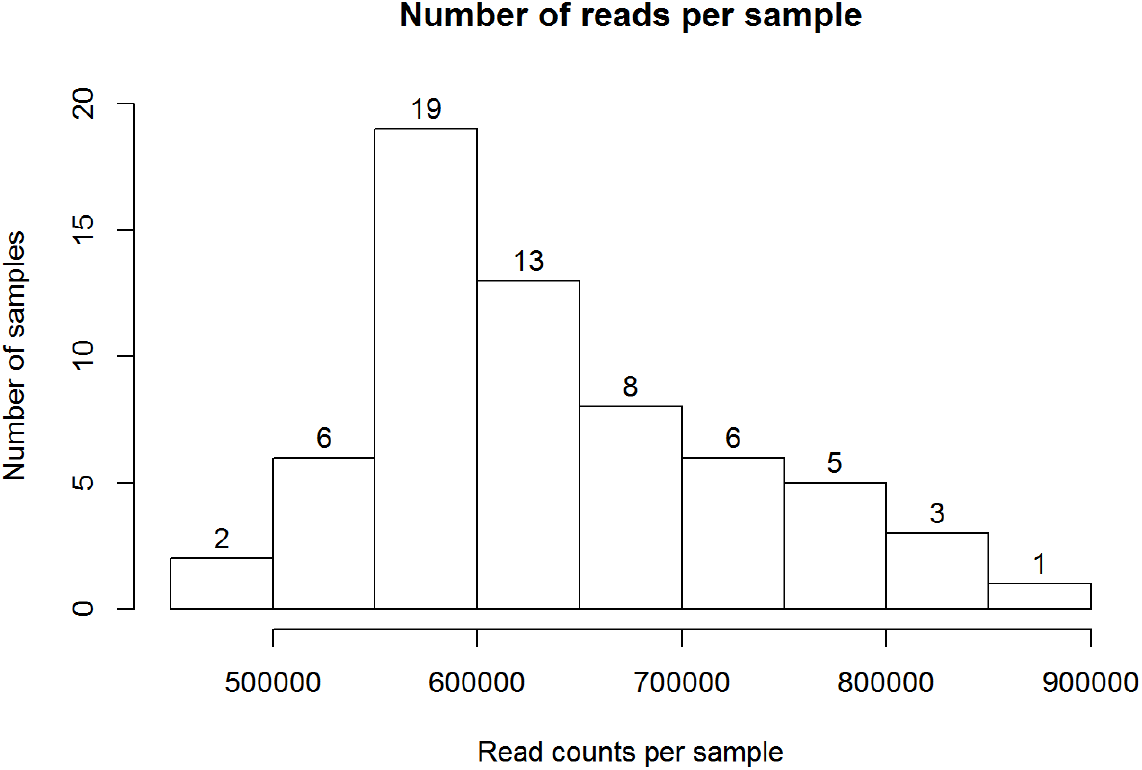
Histogram of raw read counts per sample.

### Statistical analyses

#### Analyses on gut microbiota

The analyses were divided in two parts: analysis on the whole dataset (n=63) and within the patient group only (n=23), see **Table 2** for a schematic of all analyses performed. Analysis on the whole dataset consisted of community and composition analysis on disease status (patient versus family members) and on the interaction between disease status with intestinal complaints. The patient only analysis consisted of association analyses of community and composition with phenotypic information on behavioural symptom severity (CBCL total problems score and ADOS comparative score) and on the genetic variants (mutation and deletion in the *EHMT1* gene, see above).

#### Community analyses

Global community analyses were performed without any prevalence filtering on genus and sample level to prevent inducing a bias against less prevalent genera.

##### Alpha-diversity

Shannon alpha-diversity was calculated in this study. After assessing normality using Shapiro-Wilk test, parametric ANOVAs were used.

For the analyses of the complete dataset, two separate ANOVA models were performed, one testing for effect of the factor disease status (Patient vs Family) and another including the factor intestinal complaints (yes/no), allowing us to test for an interaction between disease status and intestinal complaints. Both models were corrected for age, age^2^, BMI, gender, familyID and dietary setting.

In the patient only analyses, alpha diversity was correlated with the symptom severity measurements CBCL total score and ADOS comparative score (using Pearson’s correlation where both measures are normally distributed, Spearman’s where not). Moreover, using ANOVA we tested for differences in Shannon diversity between 1)two groups of genetic variants (mutation + deletion < 1MB, deletion > 1MB) and 2) three groups (mutation, deletion < 1MB, deletion > 1MB) both controlled for age, age^2^, BMI and gender (see **Table 2** for a an overview of analyses performed).

##### Beta-diversity

As a measure of beta-diversity, we calculated Bray-Curtis dissimilarity using the Principal Coordinate Analysis (PCoA) method (via the ordinate function in the phyloseq package). Bray-Curtis is a distance metric explaining differences in the relative abundances between microbial communities (Lozupone & Knight, 2009).

Permutation testing was performed using the adonis function in the vegan package (Oksanen et al., 2019). On the whole dataset, permutation testing was done in two models: the first one tested for the effect of disease status (Patient, Family) on beta diversity and the second one assessed an interaction between disease status and intestinal complaints (yes/no). Both models were corrected for age, age^2^, BMI, gender, dietary setting and accounted for family relatedness by using familyID in the ‘strata’ argument. This ensures permutations are constrained between and not within families.

For the patient only analyses, we assessed the effect of behavioural symptom severity (CBCL total problems score and ADOS comparative score) and genetic variants on beta diversity, correcting for age, age^2^, BMI and gender.

Besides statistical testing, beta diversity was visualized using Canonical Analysis of Principal Coordinates (CAP) (Anderson & Willis, 2003). This supervised ordination method allows visualization of variance explained by a variable of interest, rather than visualisation of the largest source of variation in the data as is the case in unsupervised ordination methods such as Principle Coordinates Analyses (PCoA).

#### Compositional analyses on the taxonomic data

Compositional analyses were performed at genus level; the 6.771 ASV’s obtained after denoising were aggregated to 417 genera. Two filtering steps removed genera containing excess zeros in the composition data, see (Szopinska-Tokov et al., 2021). First, at genus level, genera with a prevalence > 50% across all samples were selected; consisting of 138 genera (these are genera with at last 50% of non-zero values across all samples). This threshold removes noise and rare genera and includes the most abundant microbiota (see Risely et al. (2021)), therefore referring to this selection as the “core” microbiota. Second, at sample level, samples with less than 10% of genera with non-zero values are removed, which resulted in zero removed samples.

For the whole dataset, per genus we used two regression models to test for 1) a main effect of disease status and 2) an interaction effect between disease status and intestinal complaints (yes/no) on relative abundance (see **Table 2**). To abide by the skewed nature of the data, non-parametric regressions were performed using the aligned.rank.transform function of the ART package (Villacorta, 2015). In short, this function includes a pre-processing step aligning data before applying averaged ranks, and then runs a classic ANOVA on each of the aligned ranks (F-test) (Wobbrock, Findlater, Gergle, & Higgins, 2011). Rank-based statistics are very commonly used in non-parametric testing, making the statistics less vulnerable for extreme values in the distribution (Feys, 2016). Moreover, this method allows inclusion of covariates as well as interaction testing, which is not possible using other non-parametric methods such as Wilcoxson’s signed rank test. The two models included correction for age, age^2^, BMI, gender, dietary setting and familyID. In case of significant interactions between disease state and intestinal complaints, simple effects of disease state per intestinal complaints group (yes/no) were tested using Mann-Whitney U-tests.

For the patient dataset, we ran three models to test for a main effect of CBCL total score and ADOS comparative score on the relative abundance of any of the bacterial genera. These models were corrected for age, age^2^, BMI, gender. Due to the low sample size per genetic variant sub-group, no compositional analyses were performed on genetic variants. All the taxonomic analyses were FDR-corrected for the amount of tests performed (138 genera per model).

For significant results, we visually inspected the relative abundance data distribution: identifying the number of non-zero and zero observations per group. The nature of sequencing data does not allow dissociation between “true” biological zero’s from sub-threshold detection or sequencing artefacts (Kaul, Mandal, Davidov, & Peddada, 2017). Hence, interpretation of statistical results based on many zero observations should be done with caution. Working with a core microbiota already limits the chance of finding such results. The interpretation of statistical results should be done in the context of how many non-zero observations this result was based on, which is reported per group (patient and family members) in each data plot on the horizontal axis.

#### Analyses on the relation between gut microbiota and disease-related symptoms and EHMT1 genetic variation

To assist the interpretation of potential associations between disease-related characteristics (such as behavioural symptoms, intestinal complaints and genetic variants) and gut microbiota, we assessed the relation among these characteristics in the patients. We tested the relation between behavioural symptoms (CBCL and ADOS) using a Spearman correlation. CBCL and ADOS were compared between patients that did versus those that did not experience intestinal complaints using a t-test for the CBCL and a Wilcoxon test for the ADOS.

Comparison between CBCL and two groups of genetic variants was done using a t-test and for the three groups of genetic variants using an ANOVA. Comparison between ADOS and two groups of genetic variants was done using a Wilcoxon test and a Kruskal-Wallis test for the three groups of genetic variants. To test the association between intestinal complaints and the two and three groups of genetic variants, a chi-square test was used.

## Results

### Microbiota results

#### Community results

##### Alpha-diversity

###### Patients versus family members

Within the complete dataset, we observed a main effect of disease status on Shannon diversity; F(1,54)=6.400, p=0.014, where alpha diversity was lower in the patient group compared to their family members, see **Figure 2**. Intestinal complaints (yes/no) did not affect Shannon diversity; F(1,47)=0.674, p=0.416, neither did disease status interact with intestinal complaints in terms of Shannon diversity scores ; F(1,47)=0.147, p=0.703, see **Figure S1**. Both models were corrected for age, age^2^, BMI, gender, familyID and dietary setting.

**Figure 2.**
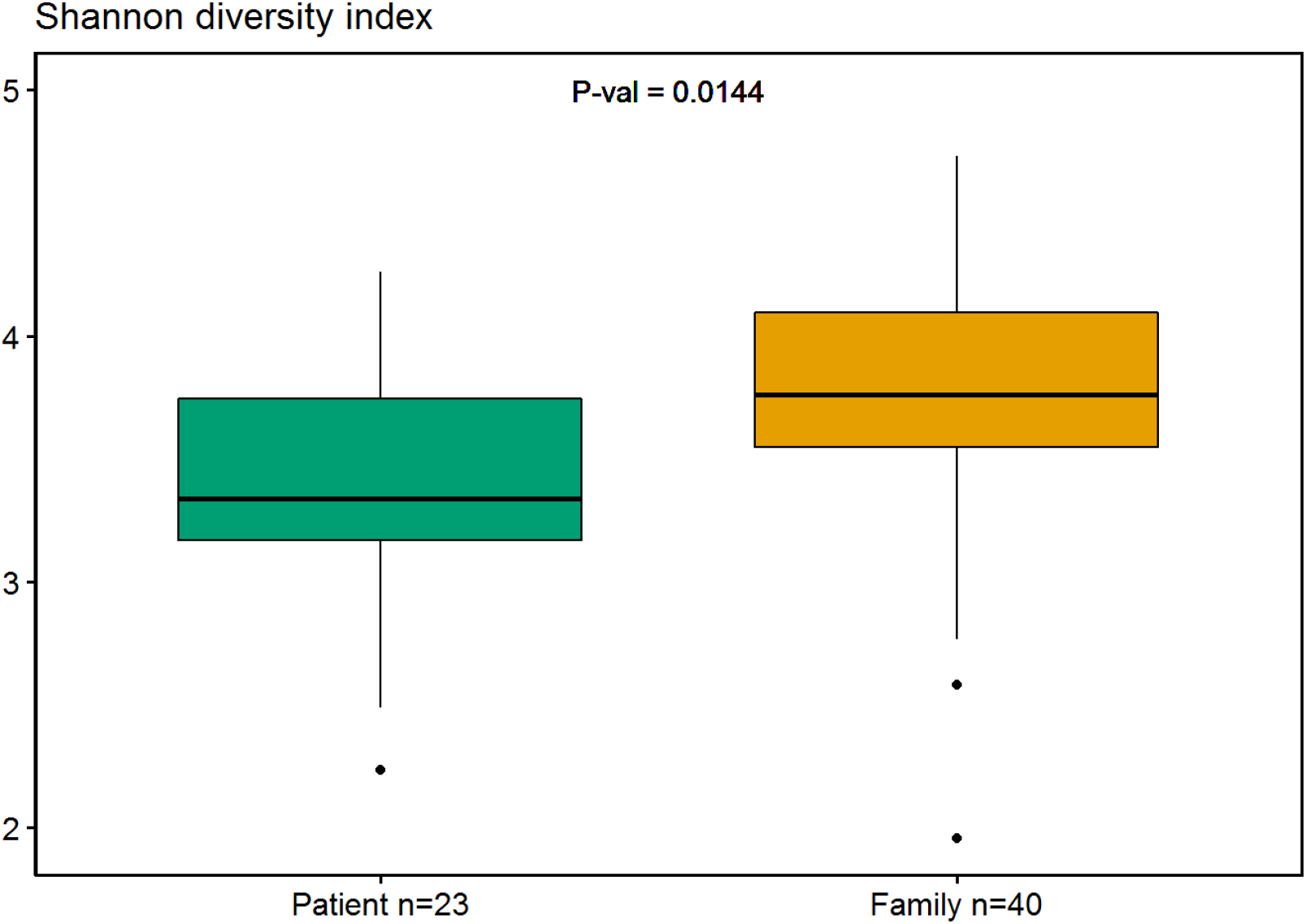
Boxplot of the main effect of disease status on Shannon diversity, with the p-value of the main effect of disease status of the ANOVA on Shannon diversity. The boxplot indicates the median (black horizontal line) and 25^th^ and 75^th^ quartiles as the outside of the box. The upper and lower horizontal lines in the boxplot represent the largest and smallest values within the 1.5 interquartile range and the dots indicate the outliers which is > 1.5 times and < 3 times the interquartile range.

###### Patient dataset

For the patients only dataset, no significant correlations were found between Shannon diversity and the CBCL (r=-0.114, p=0.614) or the ADOS (r=0.018, p=0.936). We did not observe significant differences on Shannon diversity between the two (F(1,15)=0.001, p=0.979) or the three genetic variant (F(2,14)=0.049, p=0.952) groupings see **Figure S2**.

##### Beta-diversity

###### Patients versus family members

Within the whole dataset, gut microbial community measured by Bray-Curtis dissimilarity at ASV level differed between patients and family members (a main effect of disease status F(1,55=1.873, p=0.012), see **Table 3**. Disease status explained 3% variance explained in adonis permutation testing (R2 in **Table 3**) in the model, corrected for age, BMI, gender, dietary setting and family relatedness of which none are significant confounders in this model. The CAP visualization (**Figure 3**) shows separate grouping of bacterial communities of patients and their family members.

**Figure 3.**
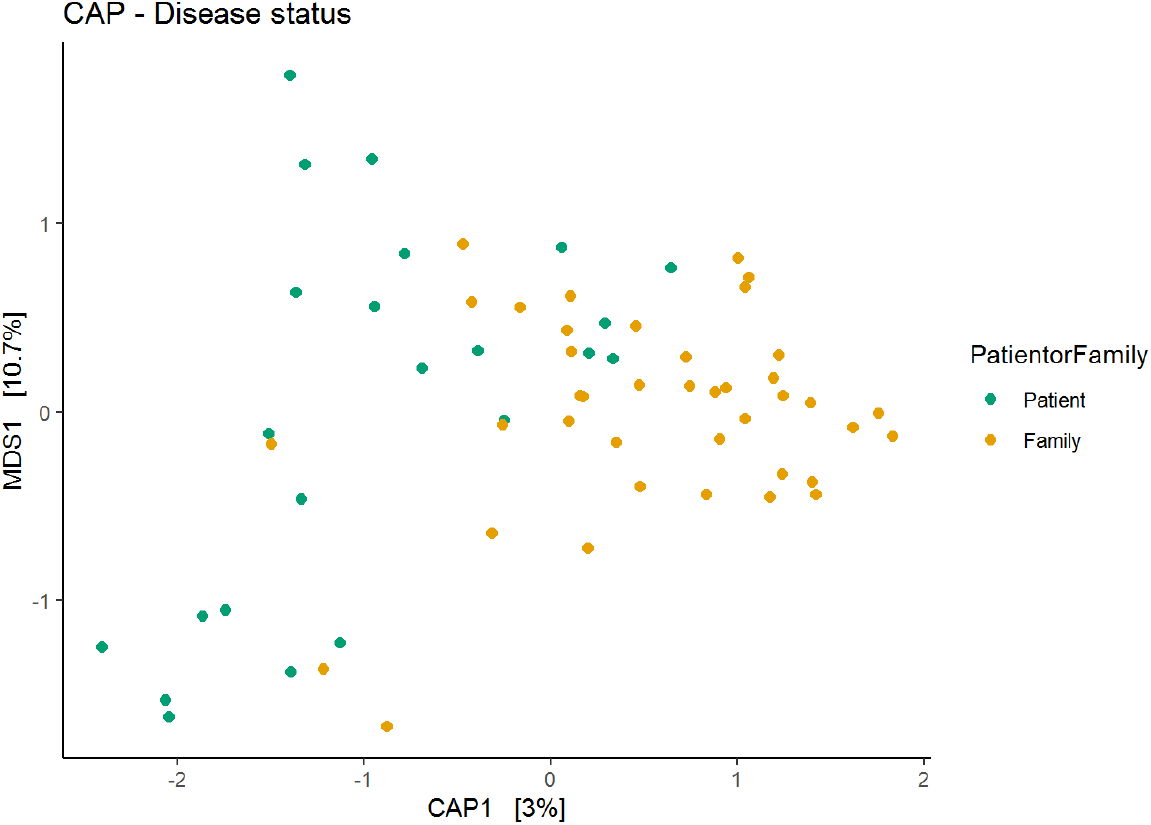
CAP plot of Bray-Curtis dissimilarity metric for disease status (CAP1). The colouring represents the disease status showing a significant effect in the permanova test, see **Table 3**.

**Table 3.** Effect disease status on Bray-Curtis beta diversity.

|  | df | SumOfSqs | R2 | Pseudo F | p-value |
| --- | --- | --- | --- | --- | --- |
| Disease status | 1 | 0.541 | 0.030 | 1.873 | 0.012 |
| Age | 1 | 0.326 | 0.018 | 1.128 | 0.258 |
| Age<sup>2</sup> | 1 | 0.195 | 0.011 | 0.676 | 0.933 |
| BMI | 2 | 0.408 | 0.023 | 0.706 | 0.965 |
| Gender | 2 | 0.355 | 0.020 | 1.228 | 0.180 |
| Dietary Setting | 1 | 0.286 | 0.016 | 0.991 | 0.445 |
| Residual | 55 | 15.879 | 0.883 |  |  |
| Total | 62 | 17.989 | 1.000 |  |  |

Legend: df: degrees of freedom, SumofSqs: sums of squares – average distances between groups, $R^2$ : variance explained, Pseudo F: signal to noise ratio of between to within group variance, p-value: fraction of permuted F ratio's that are larger than the observed F ratios.

Across all participants, intestinal complaints (yes/no) did not significantly affect beta diversity: F(1,47)=1.047, p=0.367, 1.8% variance explained, see **Figure S3**. Moreover, disease status did not interact with intestinal complaints: F(1,47)=1.135, p=0.271, 2% (R2).

###### Patients dataset

For the patient only dataset, a main effect of CBCL total score on beta diversity was observed (F(1,15)=1.654, p=0.045, 7.4% variance explained by CBCL in the model (R2), see **Table 4** and **Figure 4 - left panel**. The behavioural symptom severity measured CBCL total score associated with a differential beta diversity. No effects were observed for ADOS comparative score (F(1,14)=0.7242, p=0.815, 3.6% variance explained), see **Figure 4 - right panel**.

**Figure 4.**
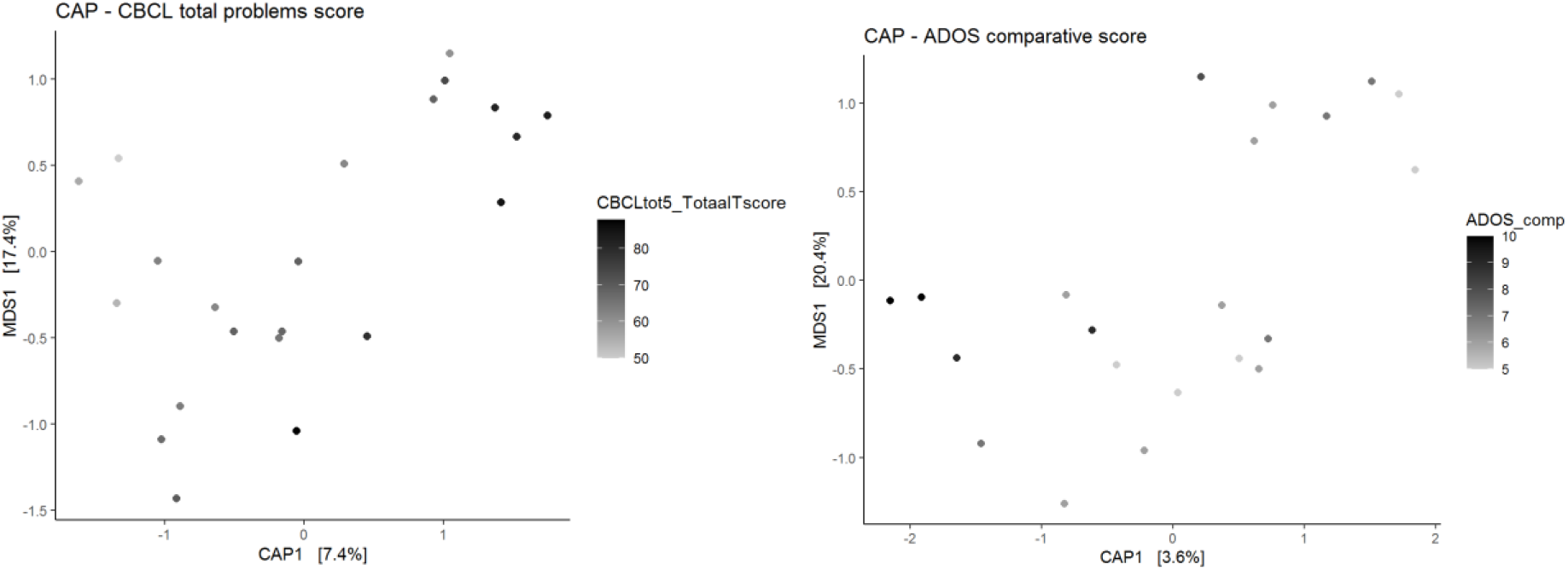
CAP plots of Bray-Curtis dissimilarity metric for CBCL total problems score (CAP1, left panel) and ADOS comparative score (CAP1, right panel). The colouring represents the CBCL total problems or ADOS comparative score where a lighter grey represents a higher score and a darker grey represents a lower score. The CBCL total problems score showed a significant difference in the permanova test, see **Table 4**.

**Table 4.**
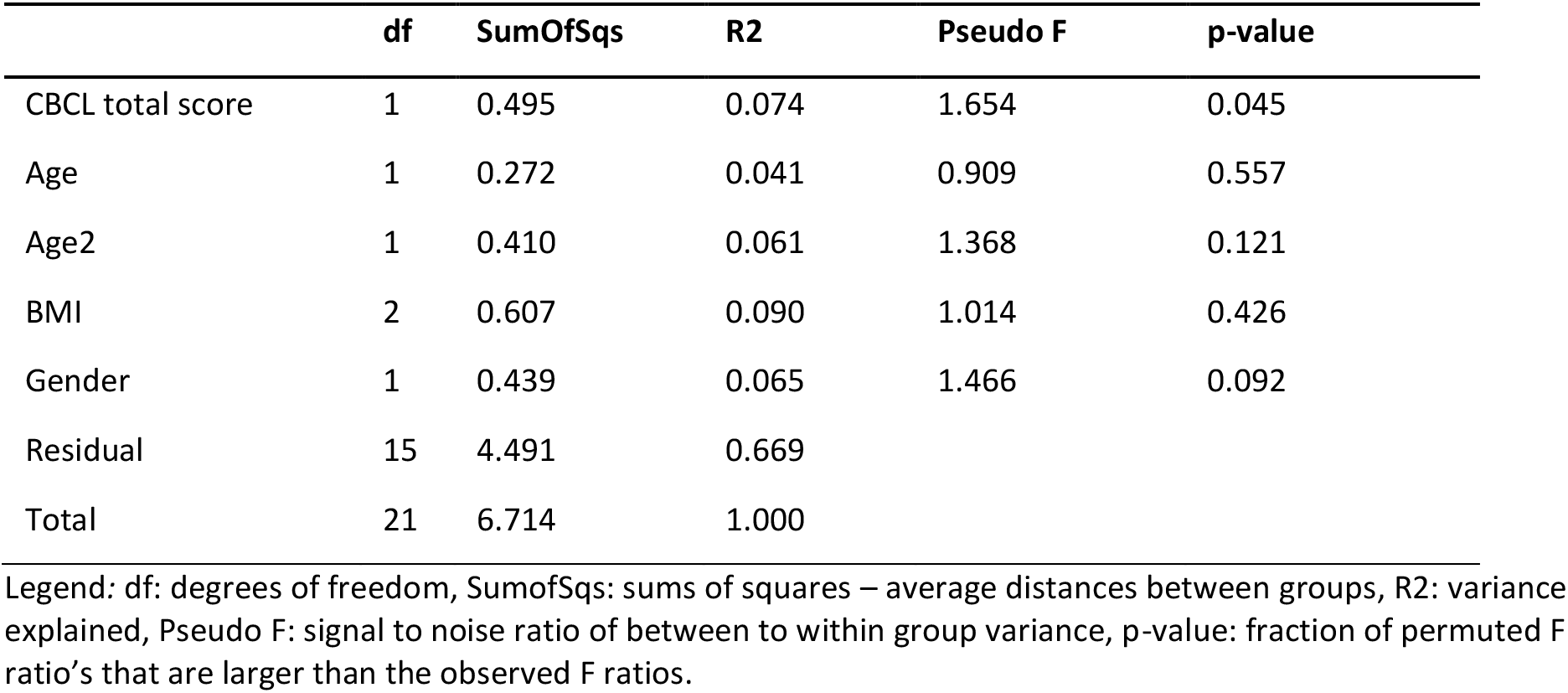
Permanova results of Bray-Curtis beta diversity for CBCL total problems score.

|  | df | SumOfSqs | R2 | Pseudo F | p-value |
| --- | --- | --- | --- | --- | --- |
| CBCL total score | 1 | 0.495 | 0.074 | 1.654 | 0.045 |
| Age | 1 | 0.272 | 0.041 | 0.909 | 0.557 |
| Age2 | 1 | 0.410 | 0.061 | 1.368 | 0.121 |
| BMI | 2 | 0.607 | 0.090 | 1.014 | 0.426 |
| Gender | 1 | 0.439 | 0.065 | 1.466 | 0.092 |
| Residual | 15 | 4.491 | 0.669 |  |  |
| Total | 21 | 6.714 | 1.000 |  |  |

Legend: df: degrees of freedom, SumofSqs: sums of squares – average distances between groups, R2: variance explained, Pseudo F: signal to noise ratio of between to within group variance, p-value: fraction of permuted F ratio's that are larger than the observed F ratios.

We did not find any significant differences in beta diversity between the 1) two genetic variant groups (F(1,15)=1.106, p=0.302, 4.8% variance explained (R2); **Figure 5 – right panel**) or the 2) three genetic variant groups (F(2,14)=0.876, p=0.683, 7.6% variance explained(R2); **Figure 5 – left panel**).

**Figure 5.**
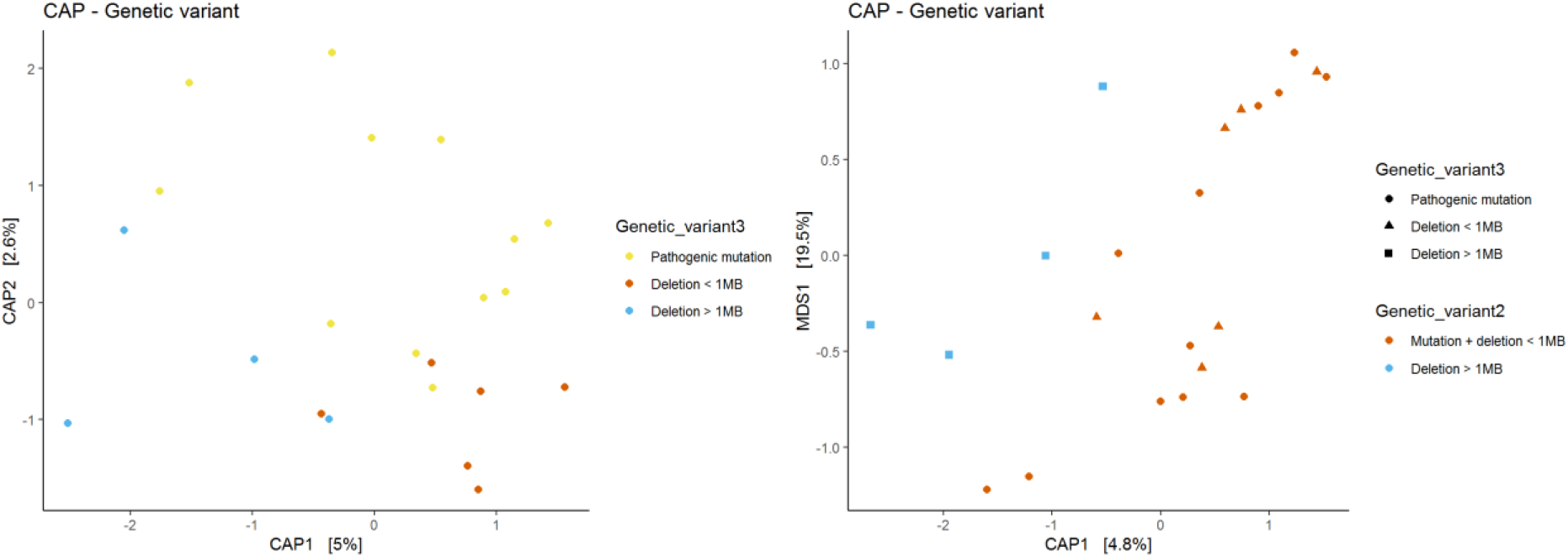
CAP plots of Bray-Curtis dissimilarity metric for genetic variants. The coloring represents the different genetic variants in three groups (pathogenic mutation, deletion < 1MB or deletion > 1MB) or in two groups (pathogenic mutation + deletion < 1MB or deletion > 1MB). In the right panel, the shapes are based on the three groups (pathogenic mutation, deletion < 1MB or deletion > 1MB) to see how the pathogenic mutation and deletion < 1MB variants cluster.

##### Compositional analyses

###### Patients versus family members

Out of 138 genera, one genus, *Coprococcus 3*, was significantly associated with disease status, relative abundance of this genus was higher in the family group compared to the patient group (F=16.303, p_uncorrected_=0.0003, p_corrected_=0.045), see **Figure 6**.

**Figure 6.**
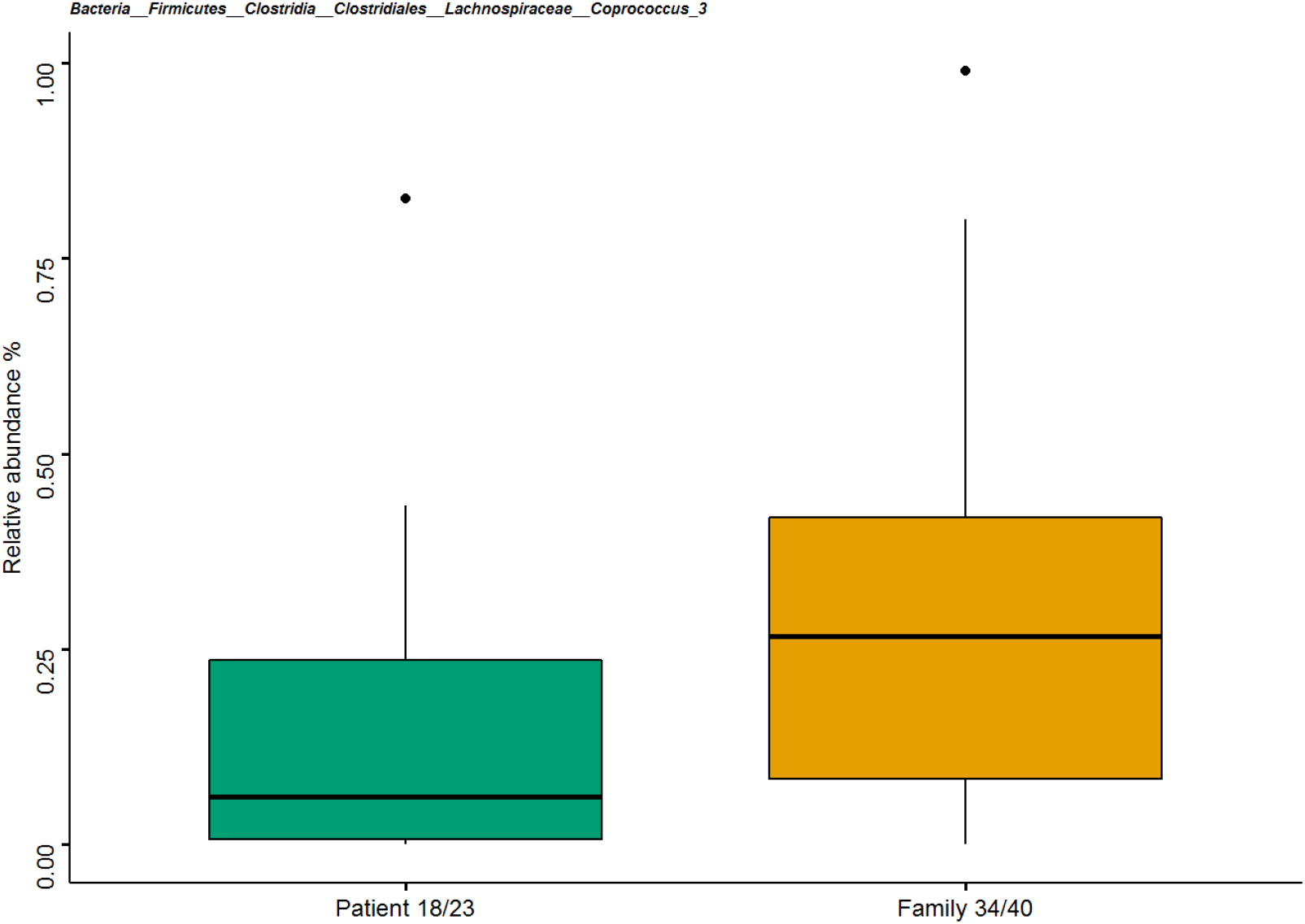
Boxplot showing the effect of disease status on relative abundance of genus *Coprococcus 3.* The boxplot indicates the median (black horizontal line) and 25^th^ and 75^th^ quartiles as the outside of the box. The lines represent the largest and smallest values within the 1.5 interquartile range and the dots represents the outliers which is > 1.5 times and < 3 times the interquartile range.

For 15 genera we observed an FDR-uncorrected effect of disease status, see Supplementary Results for more information on these, amongst others **Table S4** on the statistical results per genus and **Figure S4** for plots per genus.

###### Disease status * intestinal complaints

One genus, *Merdibacter*, showed a significant interaction effect of disease status*intestinal complaints (F=15.313, p_uncorrected_=0.0003, p_corrected_=0.040), see **Figure 7**. In those participants reporting intestinal complaints, *Merdibacter* relative abundance was lower in the patient group compared to their family members. Whilst in those participants reporting no intestinal complaints, *Merdibacter* relative abundance was higher.

**Figure 7.**
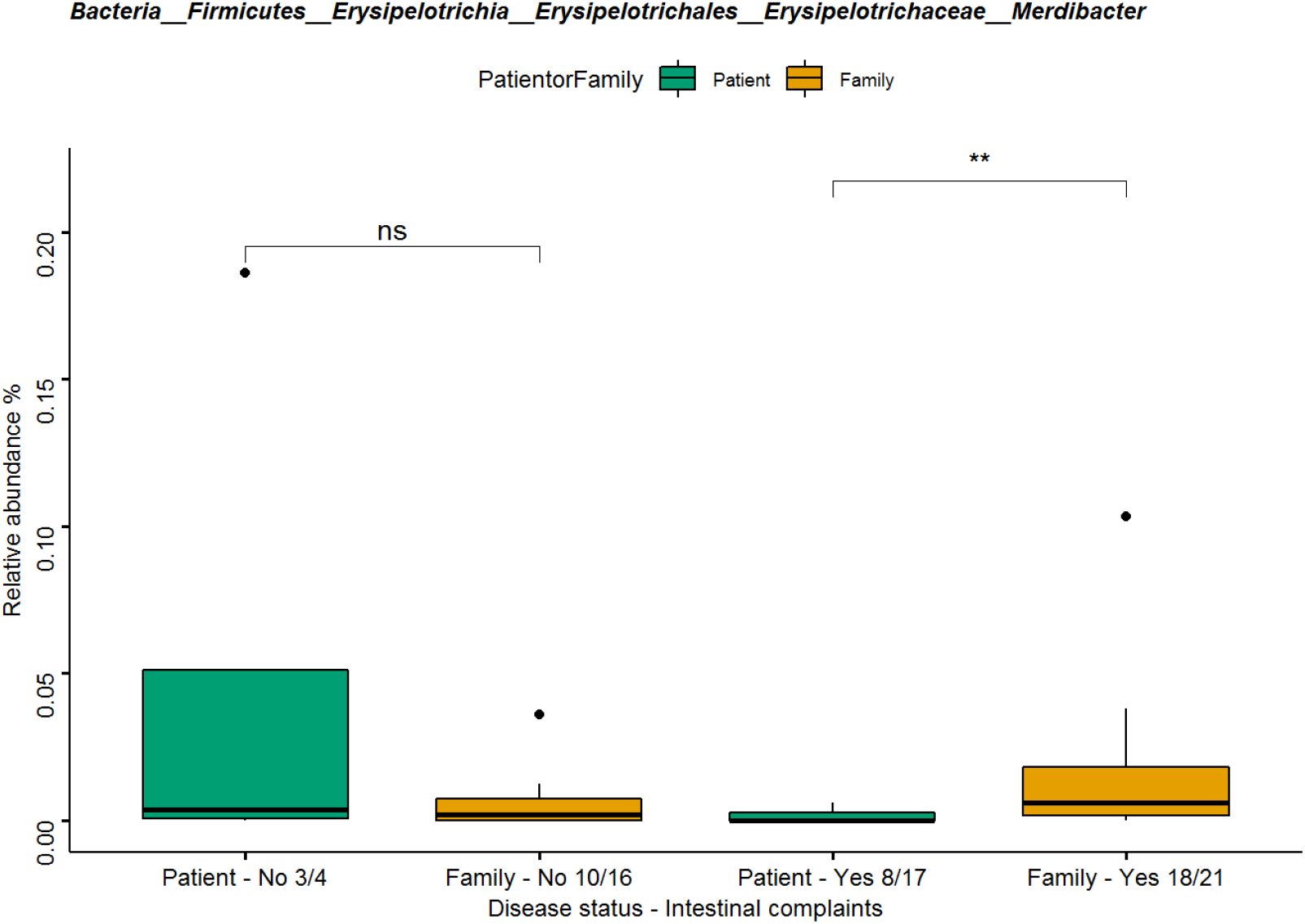
Boxplot showing the interaction effect of disease status*intestinal complaints on relative abundance of genus *Merdibacter.* The boxplot indicates the median (black horizontal line) and 25^th^ and 75^th^ quartiles as the outside of the box. The lines represent the largest and smallest values within the 1.5 interquartile range and the dots represents the outliers which is > 1.5 times and < 3 times the interquartile range. The asterisks show the results of the within test using a Mann-Whitney U-test: ns = P > 0.05, * = P < 0.05 and ** P < 0.01.

We observed an FDR un-corrected interaction effect of disease status*intestinal complaints for 17 genera,. In the Supplementary Results more information is provided, amongst others **Table S4** for the statistical results per genus and **Figure S5** for plots per genus.

##### Patients dataset

###### CBCL total score

Higher relative abundance of genus *Atopobiaceae – uncultured* was found to be significantly associated with a higher CBCL total score (F=21.141, p_uncorrected_ =0.0003, p_corrected_ =0.048), see **Figure 8**.

**Figure 8.**
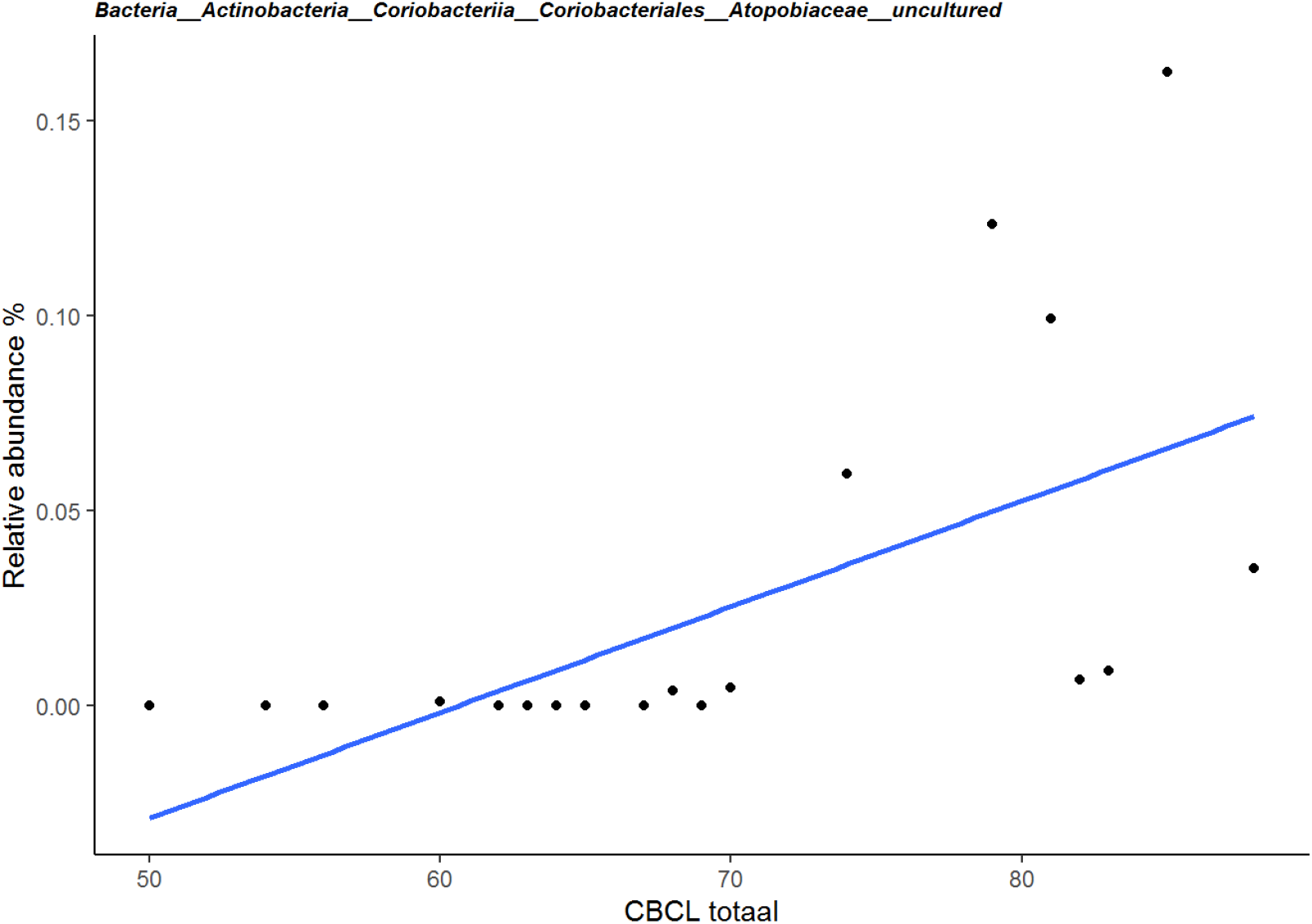
Scatter plot showing the relation between relative abundance of genus *Atopobiaceae - uncultured* and CBCL total score.

Relative abundance of seven genera showed a non-significant (FDR-uncorrected) association with CBCL total score. See the Supplementary Results, **Table S4** and **Figure S6** for more information.

##### ADOS comparative score

We did not detect any significant associations between any of the genera with the ADOS. However, four genera, *Gemella, Atopobiaceae – uncultured, Butyricicoccus* and *Veillonella*, showed suggestive evidence for association with the ADOS comparative score. See Supplementary Results, **Table S4** and **Figure S7** for more information.

### Relation between disease-related symptoms and genetic variants

Scores on ADOS and CBCL scales did not correlate significantly (r= −0.113, p= 0.625). Regarding the relation with intestinal complaints, we observed a significant difference in CBCL total score (t=-3.171, p=0.018), showing that high behavioural symptom severity was associated with frequent intestinal complaints, see **Figure S8.** ADOS scores did not differ between patients experiencing and those not experiencing intestinal complaints (W=28, p=0.732).

None of the genetic variant groups showed differences in CBCL or ADOS scores (CBCL: two groups (t=-0.181, p=0.872, three groups (F(2,17)=0.114, p=0.893), ADOS comparative score: two groups (W=29.5, p=0.712), three groups (χ^2^=5.002, p=0.082)). No significant relation was found between intestinal complaints and two groups (χ^2^=1.523e-30, p=1) or three groups of genetic variants (χ^2^=1.593, p=0.451).

## Discussion

### Summary

In this first exploratory study on the gut microbiota in KS we compared the gut microbiota of 23 patients to 40 of their family members (n=14 fathers, n=16 mothers and n=10 siblings). Alpha diversity was lower in KS patients compared to their family members and beta diversity of the patients clustered separately from family members. CBCL scores associated significantly with beta diversity, explaining 7% of variance in the microbial community. We detected significant taxonomic differences (at genus level) in relative abundance between patients with KS and their family members. Relative abundance of genus *Coprococcus 3* was significantly lower in patients compared to their family members. Moreover, an interaction between disease status and intestinal complaints was observed for genus *Merdibacter.* That is, for those participants reporting intestinal complaints, *Merdibacter* relative abundance was lower in patients (versus family members), while the opposite pattern was seen in participants without intestinal complaints. Within the patient group, we found higher levels of an uncultured *Atopobiaceae* genus associated with more behavioral symptoms on the CBCL.

### Gut microbial community differences in Kleefstra Syndrome

The gut microbial community of KS patients is less diverse and clusters separately from that of their family members, indicating the microbial community consists of different types and lower abundance of bacteria. Such findings are found in several types of NDDs, both monogenic as well as multifactorial. For example, ASD as well as Down Syndrome, Rett Syndrome and ADHD are associated with reduced alpha and altered beta diversity compared to controls or family members (Biagi et al., 2014; Borghi & Vignoli, 2019; Bundgaard-Nielsen et al., 2020), although few experimental studies are yet available. Comparison to these related NDDs is relevant as there are no studies on the gut microbiota in KS. Moreover, patients with KS often also display ASD-like symptoms and meet ASD criteria, hence gut microbial differences and underlying mechanisms may partially overlap.

Within the patient group, no significant effect of genetic variants on the gut microbial community was observed. Despite this and a small sample size, visually separate clustering was seen between these different genetic variants (**Figure 5**). This suggests a potential “dose effect” of *EHMT1* genetic variants on the microbiota diversity. Such dose effects are known in the clinical phenotype (i.e. larger deletions are associated with worse behavioral and somatic symptoms and overweight (BMI > 25) and high birth weight is more prevalent in KS patients carrying a mutation rather than a deletion (Kleefstra et al. 2009, Willemsen et al., 2012)) and may hence exist also at gut microbial level.

### Gut microbial composition differences in Kleefstra Syndrome

When zooming into the different bacteria, abundance of the genus *Coprococcus 3* was lower in KS patients versus their family members. Interestingly, also in children with ASD *Coprococcus* was lowered (Kang et al., 2013). Moreover, *Coprococcus* abundance associated with altered gut-brain networks in patients with irritable bowel syndrome (Labus et al., 2019). Specifically, in healthy controls *Coprococcus* abundance showed a triangular relation with gastrointestinal pain experienced during a nutrient and lactulose challenge test and resting state activity in the left caudate nucleus. These relations were absent in irritable bowel syndrome patients. In this context, genus *Coprococcus 3* may be part of mechanisms underlying ASD pathology and gastro-intestinal inflammation, associated with the neural processing of gastro-intestinal signals, which highlights the role of the gut-brain axis in NDDs. This is relevant especially as gastro-intestinal symptoms are frequently reported in KS patients. Note that in the above mentioned studies *Coprococcus* was not further specified into *Coprococcus 3* as is the case for the finding in our study. This is a difference in taxonomic classification likely caused by differences in classifier (version) used between studies. Hence the findings in the studies by Kang et al. and Labus et al. may concern a grouping of related, but not identical *Coprococci* bacteria. In terms of function, *Coprococci* strains produce butyrate (Duncan, Barcenilla, Stewart, Pryde, & Flint, 2002; Pryde, Duncan, Hold, Stewart, & Flint, 2002), a short-chain-fatty acid essential for gut health, immune functioning and neurodevelopment (Dalile, Van Oudenhove, Vervliet, & Verbeke, 2019). Blasting of the most frequent observed sequence classified into *Coprococcus 3* resulted in an unpublished hit and in one published study on the pharmacodynamics of the diabetes medication metformin (Kim et al., 2021). More information is listed in the Supplementary Discussion. Lastly, the non FDR-significant results of interest in the context of this exploratory study are reviewed in the Supplementary Materials (**Table S4** and Supplementary Discussion).

### Associations between gut microbiota, behavioral symptoms and intestinal complaints

Within the patient group, we showed that the total t-score on the CBCL explains a significant part of the variance in the microbial community (7% variance explained in beta diversity). At taxonomic level, the abundance of the uncultured genus *Atopobiaceae* correlated positively with behavioural symptom severity.

Note that this result is build up by several zero-observations (n=12 zero versus n=7 non-zero observations), hence we cannot verify whether these zero observations are true biological zeros or reflect under-sampling. As this is still an uncultured bacterium, not much is known about its function in the human intestinal tract and GBA. Blasting of the most frequent observed sequence resulted in one unpublished hit and one published hit on the gut microbiota in patients with schizophrenia (S. Li et al., 2020). More information is listed in the Supplementary Discussion.

Another positive (uncorrected) correlation with CBCL was observed for *Coprococcus 3*, which was also observed to be lower in the patients versus family member comparison (see above). This combination of results suggests this genus may be a relevant microbial signature belonging to KS.

In terms of intestinal complaints, the abundance of the genus *Merdibacter* was lower in KS patients compared to their family members, but only in those participants reporting intestinal complaints. *Merdibacter* is a relatively new discovered genus (Anani et al., 2019). Blasting of the most frequent sequence resulted in eight hits, describing this bacterium as a butyrate producer in chickens (Eeckhaut et al., 2011). Moreover, it is found in studies on obesity (Ley, Turnbaugh, Klein, & Gordon, 2006), recurrent antibiotic induced Clostridium difficile infection (Chang et al., 2008) and ulcerative colitis (Lepage et al., 2011). These results could be viewed as early indication that gut microbial subgroups exist within the KS diagnosis based on intestinal complaints, where *Merdibacter* interacts with the developmental alterations (of the gut) associated with KS.

### Strengths and limitations

This is the first exploratory study into the gut microbiota of KS. Access to a clinical sample is rare due to the low disease prevalence, hence the characterization of 23 KS patients is a relevant first step in the identification of gut microbial signature of the syndrome. Our sample has a broad range of characterization including behavioral information, medication, diet and intestinal complaints. The control group consisting of family member is very suitable, as almost all KS cases are caused by de novo genetic aberrations, hence the EHMT1 genotype is the biggest source of variation between them. Moreover, their dietary intake will largely be similar. Although some patients may have a more restricted diet due to their behavioral symptoms, indeed we did not see big differences between intake of the largest food groups. The use of family members also had some limitations: there was a strong imbalance in age between patients (mean age 18.8) and family members (mean age 36.8) included. For this reason we included age and linear additive effects of age (age^2^) as covariates in our analysis. Besides the age difference between family members and patients, there is a large age range *within* the patient group. The age of a KS patient is relevant for their disease status as during adolescence a majority suffers from regressive episodes. This may also interact with their gut microbial profile. To illustrate, Schwarz et al. (2018) observed that subjects with larger differences in gut microbiota composition in their first psychotic episode compared to healthy controls, had poorer mental health over 12 months follow-up. Similarly, older KS patients may display more severe regression based on their microbial profile and fluctuations. The sample size in the current study does not allow comparison within the patient group between those that went through such an episode to those that have not, stratified by age. Also within the patient group analyses included age and age^2^ covariates to correct for the broad age range.

Another imbalance is medication use; 16 out of 23 patients versus 7 out of 38 family members reported medication use. Patients also have a much longer medication list. We acknowledge that medication impacts the gut microbiota and can contribute to differences between patients (with KS) and comparison groups (Jackson et al., 2018; Maier et al., 2018; McGovern, Hamlin, & Winter, 2019). Nevertheless, medication use was not controlled for in the analyses, as all such strategies in this dataset will be flawed especially given our sample size. For example, excluding patients with medication use will only leave 6 samples in this group to compare with 36 family members. Moreover, medication use in KS patients often include psychotropics such as anti-psychotics and sedatives, which are therefore in themselves an indication of disease and symptom severity. This means that when testing for an association with behavioral symptom severity, medication use is colinear with disease severity and cannot be properly controlled for. When testing for effects of disease status, depending on the type of medication, patients may never go without and hence one can argue that the effect of medication on the gut microbiota is part of the KS patient phenotype.

### Future directions

The current findings should be tested for replication in a larger sample. For example, including more siblings will form a separate, better age-matched, comparison group. Moreover, higher sample size per genetic variant will extend analyses on microbial profile differences between genetic variants. Including a broader age range of patients will allow testing of the stability of gut microbial signatures of KS across age or with regression stage. Furthermore, intestinal complaints may be more extensively characterized with validated questionnaires, moving beyond the rough semi-quantitative inventory that was currently done for feasibility reasons. It is noteworthy that a study by Strati et al. (2016) in patients with Rett Syndrome did define constipation according to the official Rome III criteria and also found no effect on the global community measures (alpha and beta diversity). Similarly, dietary patterns may be studied more in depth, even though a proper assessment requires an extensive time and effort both from the participant as well as a certified nutritionist scoring the dietary diary or Food Frequency Questionnaire. Illustrating the relevance of proper dietary control in gut microbial case control comparisons is the recent paper on the bidirectional relation between diet and gut microbial differences associated with ASD (Yap et al., 2021).

Tapping more closely into mechanisms driving gut-brain alterations, in future studies we would like to assess metabolic characteristics of KS patients. The EHMT gene affects metabolic processes; such as (brown) fat metabolism (Nagano et al., 2015) and energy metabolism under stress conditions, as observed in EHMT/G9a knockout fruit flies (Riahi et al., 2019). Mitochondrial metabolism is disturbed in ASD and other NDDs (Panneman, Smeitink, & Rodenburg, 2018; Rossignol & Frye, 2012). These defects in mitochondrial fat/energy metabolism in KS and related NDDs could also interact with the gut microbiota, who’s metabolites modulate metabolic health (Franco-Obregón & Gilbert, 2017). By studying such host-microbiota interactions in KS patients we may identify suitable pathways for microbial interventions such as probiotics.

In relevant initiatives, overlapping microbial signatures across various types of NDDs (monogenetic such as Rett and Down Syndrome and multifactorial such as ASD) are reviewed (Borghi & Vignoli, 2019; Orozco, Hertz-Picciotto, Abbeduto, & Slupsky, 2019). With more studies on microbiota in NDDs emerging, these initiatives can be up-scaled and extended to other NDDs including KS, characterizing shared as well as unique microbial alterations in NDDs. The ultimate goal is this research is to flag gastro-intestinal complaints in an early stage and design interventions targeted at the gut microbiota such as probiotics to improve developmental trajectories and quality of life of KS patients and their caregivers.

### Conclusion

In conclusion, our results suggest that gut microbial alterations are an integral part of the KS phenotype, perhaps via processing of gastro-intestinal signals, which could exacerbate behavioral complaints. Understanding potential alterations in the gut microbiota of individuals with KS may bring improved therapeutic guidance for this patient group. For example, novel treatment strategies, such early life probiotics supplementation may support better gut microbial health with the ultimate aim to relive their GI symptoms and, through the GBA, their cognitive abilities and behavioral symptoms.

## Acknowledgments

The authors thank Maran Koolen, Melike Şimşek, Jeff Severijns and Hajar Rotbi for their help with data processing and literature search. The research leading to these results received funding from the European Community’s Horizon 2020 research and innovation programme through the Eat2beNICE (grant agreement no. 728018), CoCA (grant agreement no. 667302) and CANDY (grant agreement no. 847818) projects, by the Dutch Research Council grant (015.014.036 and 1160.18.320) and Netherlands Organization for Health Research and Development (91718310). Several authors of this publication are members of the European Reference Network on congenital malformations and rare intellectual disability (ITHACA).

