## Supplementary Materials for "The role of the gut microbiota in patients with Kleefstra syndrome"

**Index**

**Supplementary Materials**

**Supplementary Results**

- *Lifestyle questionnaire data*
- *Microbiota data*
  - *Community analyses*
    - *Alpha-diverstity*
      - *Patients versus family members*
      - *Patients dataset*
    - *Beta-diversity*
      - *Patients versus parents*
  - *Taxonomic results*
    - *Patients versus family*
    - *Patients dataset*
- *Relation between disease-related symptom severity and genetic variants*
- *Results overview*

**Supplementary Discussion**

**Supplementary Materials**

**Table S1. Missing values across all measures, indicated with an “x”, in the dataset.**

| **SampleID** | **Family** | **BMI*** | **Intestinal complaints** | **CBCL total problems score** | **ADOS comparative score** | **Genetic variant** |
| --- | --- | --- | --- | --- | --- | --- |
| KSMB3 | Patient |  | X |  |  |  |
| KSMB10 | Patient |  |  | X |  |  |
| KSMB22 | Patient |  |  |  | X |  |
| KSMB23 | Patient | X | X |  |  | X |
| KSMB27 | Patient | X |  |  |  |  |
| KSMB11003 | Sibling | X | X | NA | NA | NA |
| KSMB12003 | Sibling |  |  | NA | NA | NA |
| KSMB15001 | Father | X | X | NA | NA | NA |
| KSMB27003 | Sibling | X |  | NA | NA | NA |
| KSMB27004 | Sibling | X |  | NA | NA | NA |
| KSMB3003 | Sibling | X | X | NA | NA | NA |

* the variable BMI is imputed with the median of the family group the sample belongs in.

**Supplementary Results**

*Lifestyle questionnaire data*

**Table S2.** **Descriptive statistics for the lifestyle questionnaire data.** The mean and range for the food groups are derived from the answers to the questions; 4) Daily, 3) Weekly, 2) Monthly, 1) Never. When the mean is high, the frequency of consumption is also high. None of the patients were tube fed.

| **Variables** | **Options** | **Patients** | **Family** |
| --- | --- | --- | --- |
| **N** |  | 22 | 38 |
| **Processed meats** | Mean | 3.57 | 3.45 |
|  | Range | 3-4 | 2-4 |
| **Fish** | Mean | 2.59 | 2.37 |
|  | Range | 1-3 | 1-3 |
| **Fruit** | Mean | 4 | 3.76 |
|  | Range | 4-4 | 2-4 |
| **Legumes** | Mean | 3.18 | 3.00 |
|  | Range | 3-4 | 2-4 |
| **Vegetables** | Mean | 3.86 | 3.74 |
|  | Range | 3-4 | 3-4 |
| **Sweetened beverages** | Mean | 3.45 | 3.70 |
|  | Range | 2-4 | 2-4 |
| **Alcohol during the week** | Mean | 1 | 1.15 |
|  | Range | 1-1 | 1-3 |
| **Alcohol during weekends** | Mean | 1 | 1.38 |
|  | Range | 1-1 | 1-3 |
| **Chocolate** | Mean | 1.82 | 2.08 |
|  | Range | 1-3 | 1-3 |
| **Milk** | Mean | 2.14 | 1.82 |
|  | Range | 1-4 | 1-4 |
| **Smoking** | Yes | 0 | 7 |
|  | No | 22 | 23 |
|  | In the past | 0 | 8 |
| **Medication** | Yes | 16 | 7 |
|  | No | 6 | 31 |
| **Allergies** | Yes | 4 | 4 |
|  | No | 18 | 33 |
| **Complaints Physical health** | Always | 0 | 1 |
|  | Most of the time | 2 | 1 |
|  | Part of the time | 4 | 1 |
|  | Small part of the time | 5 | 14 |
|  | Never | 11 | 20 |
| **Mental health** | Always | 3 | 0 |
|  | Most of the time | 2 | 1 |
|  | Part of the time | 3 | 3 |
|  | Small part of the time | 6 | 9 |
|  | Never | 8 | 24 |
| **Intestinal complaints** | Never | 4 | 17 |
|  | Less than 1 day a month | 2 | 6 |
|  | 1 day a month | 3 | 5 |
|  | 2-3 days a month | 4 | 5 |
|  | 1 day a week | 4 | 2 |
|  | More than 1 day a week | 4 | 2 |
|  | Every day | 0 | 1 |
| **Nausea** | Always | 0 | 0 |
|  | Most of the time | 0 | 0 |
|  | Part of the time | 3 | 1 |
|  | Small part of the time | 3 | 7 |
|  | Never | 11 | 13 |

**Table S3. Overview of medication use for patients and their family members**

| **SampleID** | **Family** | **Medication** | **Dosage** |
| --- | --- | --- | --- |
| KSMB1 | Patient | Topicorte  Latanoprost  Olanzapine  Aripiprazol | If needed  50 microgram/day  25 mg/day  5 mg/day |
| KSMB2 | Patient | No | - |
| KSMB3 | Patient | No | - |
| KSMB4 | Patient | Orthica Orthiflor  Davitamon complete  Metamucil  Doxycicline  Olanzapine  Pantoprazol  Magnesiumhydroxide  Keppra  Frisium | Daily  Daily  Twice a day  100 mg/day  10mg / day  40mg / day  Daily  1000 mg / day  20 mg / day |
| KSMB5 | Patient | Otiflox UD  Temazepam  Lorazepam  Olanzapine  Natrium Valproaat  Euthyrox  Desmopressin acet  D-cura drink  Macrogol  Dulcolax tablets  Magnesium Had tablets | If needed 9mg/ml / day  If needed 10mg  3 mg/day  20mg / day  1500 mg / day  50 mcg / day  0.3 mg / day  Every three months 10000IE  NA  If needed 10mg  2896 mg / day |
| KSMB6 | Patient | Omeprazol  Movicolon  Dipiperon | 40mg / day  2 sachets / day  40mg, 2 daily 8 drops |
| KSMB7 | Patient | Florax  Rivotril  Omeprazol  Lorazepam  Dipiperon | 3 sachets / day  0.75mg / day  80mg / day  3mg / day  80mg / day |
| KSMB10 | Patient | Lueva  Keppra  Akineton  Levocetricine | 75mg / day  90ml / day  6mg / day  3.75mg / day |
| KSMB11 | Patient | Forlax  Misalasine | 20mg / day  3gr / day |
| KSMB12 | Patient | Ethinylestradiol/Levonorgestrel  Cyklokapron | 0.03/0.15  200mg |
| KSMB13 | Patient | s-adenosyl methionine  Microgynon  Fluoxetine  Circadin  Omeprazol | 800mg  50mg  20mg  2mg  20mg |
| KSMB16 | Patient | Aripiprazol  vitamine D  Macrogol  Citalopram | NA  NA  NA  NA |
| KSMB17 | Patient | No | - |
| KSMB20 | Patient | Forlax | 49 / day |
| KSMB22 | Patient | Pantoprazol  Diazepam klysma  Macrogol  Olanzapine  Carbamazepine | 80mg / day  If needed 10mg  1 sachet  20mg / day  20 ml and 15ml |
| KSMB23 | Patient | Colecalciferol  Ethinylestradiol/Levonorgestrel  Olanzapine  Omeprazol | NA  NA  5 mg morning and 10mg evening |
| KSMB24 | Patient | Esomeaprazol  Depakine | 20mg / day  1500mg / day |
| KSMB25 | Patient | Fusidin  Probiotics  Aripiprazol  Omeprazol | NA  Twice a day  10mg  20mg |
| KSMB26 | Patient | No | - |
| KSMB27 | Patient | Forlax | 4gr |
| KSMB28 | Patient | Macrogol | NA |
| KSMB29 | Patient | No | - |
| KSMB30 | Patient | No | - |
| KSMB1001 |  | No | - |
| KSMB1002 |  | Atorvastatine  Metoprolol succinaat retard | 20mg / day  25mg / day |
| KSMB2002 |  | No | - |
| KSMB2003 |  | No | - |
| KSMB3001 |  | No | - |
| KSMB3002 |  | No | - |
| KSMB3003 |  | No | - |
| KSMB6001 |  | No | - |
| KSMB6002 |  | No | - |
| KSMB6003 |  | No | - |
| KSMB10001 |  | No | - |
| KSMB10002 |  | No | - |
| KSMB10003 |  | No | - |
| KSMB11003 |  | No | - |
| KSMB12001 |  | No | - |
| KSMB12002 |  | Spironolacton  Furosemide  Fenprocoumon  Calci chew  Opsumit  Thyrax  Azatioprine  Omeprazol | 25mg  40mg  3mg  500mg/800IE  10mg  0.1mg  50mg  20mg |
| KSMB12003 |  | No | - |
| KSMB15001 |  | No information available | - |
| KSMB15002 |  | Anticonceptics | NA |
| KSMB17001 |  | No | - |
| KSMB17002 |  | Anticonceptics | NA |
| KSMB20001 |  | No | - |
| KSMB20002 |  | No | - |
| KSMB21002 |  | No | - |
| KSMB22001 |  | No | - |
| KSMB22002 |  | No | - |
| KSMB26001 |  | No | - |
| KSMB26002 |  | No | - |
| KSMB26003 |  | No | - |
| KSMB27001 |  | No | - |
| KSMB27002 |  | No | - |
| KSMB27003 |  | No | - |
| KSMB27004 |  | No | - |
| KSMB28001 |  | No | - |
| KSMB28002 |  | No | - |
| KSMB28003 |  | No | - |
| KSMB30001 |  | No | - |
| KSMB30002 |  | No | - |
| KSMB11a |  | No | - |
| KSMB2a |  | No information available | - |

*Microbiota data

Community analyses

Alpha-diverstity

Patients versus family members* **Figure S1. Boxplot of the interaction effect of disease status*intestinal complaints on Shannon diversity**. The boxplot indicates the median (black horizontal line) and 25^th^ and 75^th^ quartiles as the outside of the box. The lines represent the largest and smallest values within the 1.5 interquartile range and the dots represents the outliers which is > 1.5 times and < 3 times the interquartile range. The p-value represents the main effect of disease status of the ANOVA on Shannon diversity.

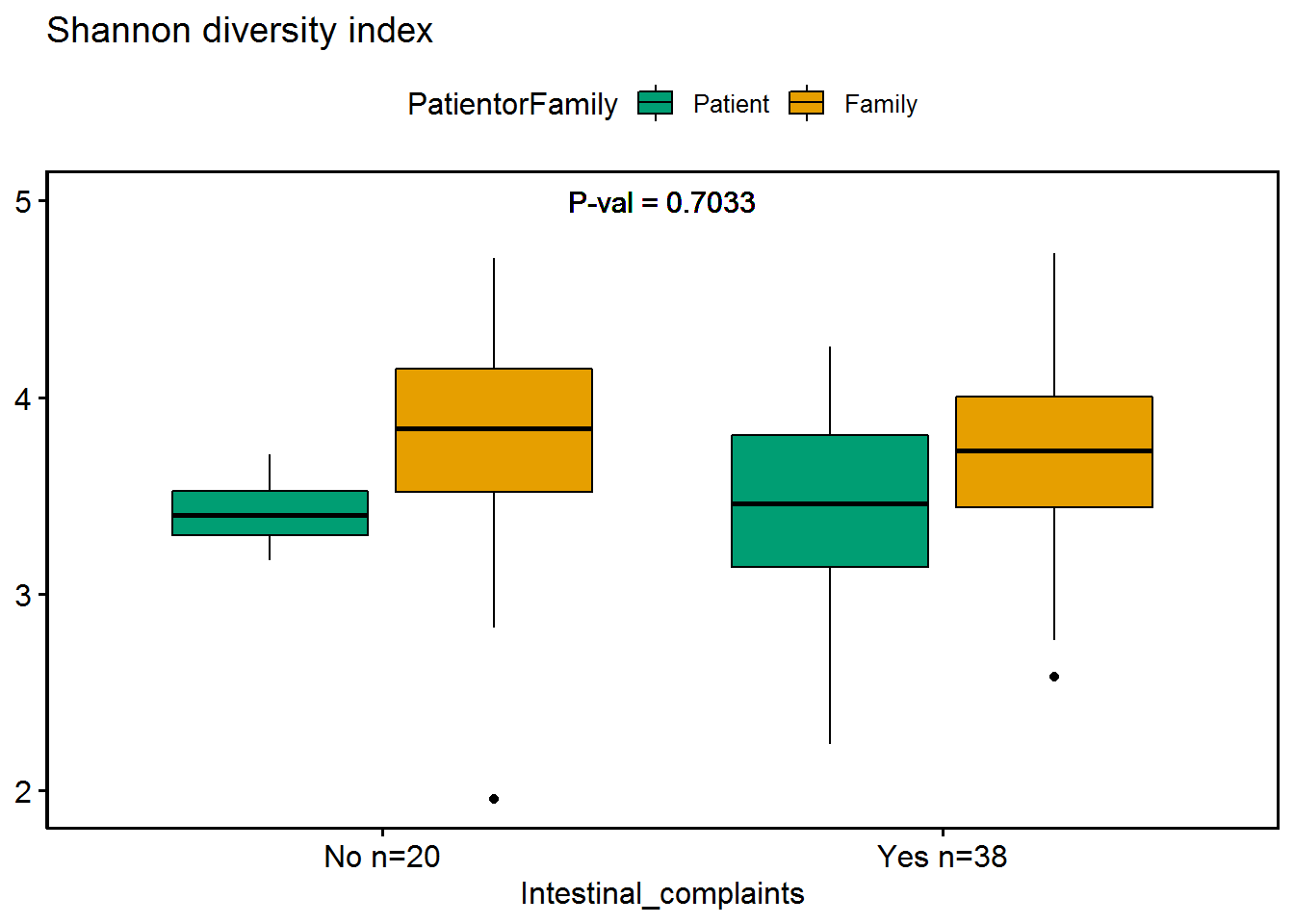

*Patients dataset*

**Figure S2.** **Boxplots of Shannon diversity for main effect of the genetic variants using three groups (pathogenic mutation, deletion < 1MB and deletion > 1MB) and two groups (pathogenic mutation + deletion < 1MB and deletion > 1MB, in the patients only dataset (n=22).** The boxplot indicates the median (black line) and 25^th^ and 75^th^ quartiles as the outside of the box. The lines represent the largest and smallest values within the 1.5 interquartile range and the dots represents the outliers which is > 1.5 times and < 3 times the interquartile range. The p-value represents the main effect of disease status of the ANOVA on Shannon diversity. The within effects are tested with a Mann-Whitney U-test. NS means p > 0.05.

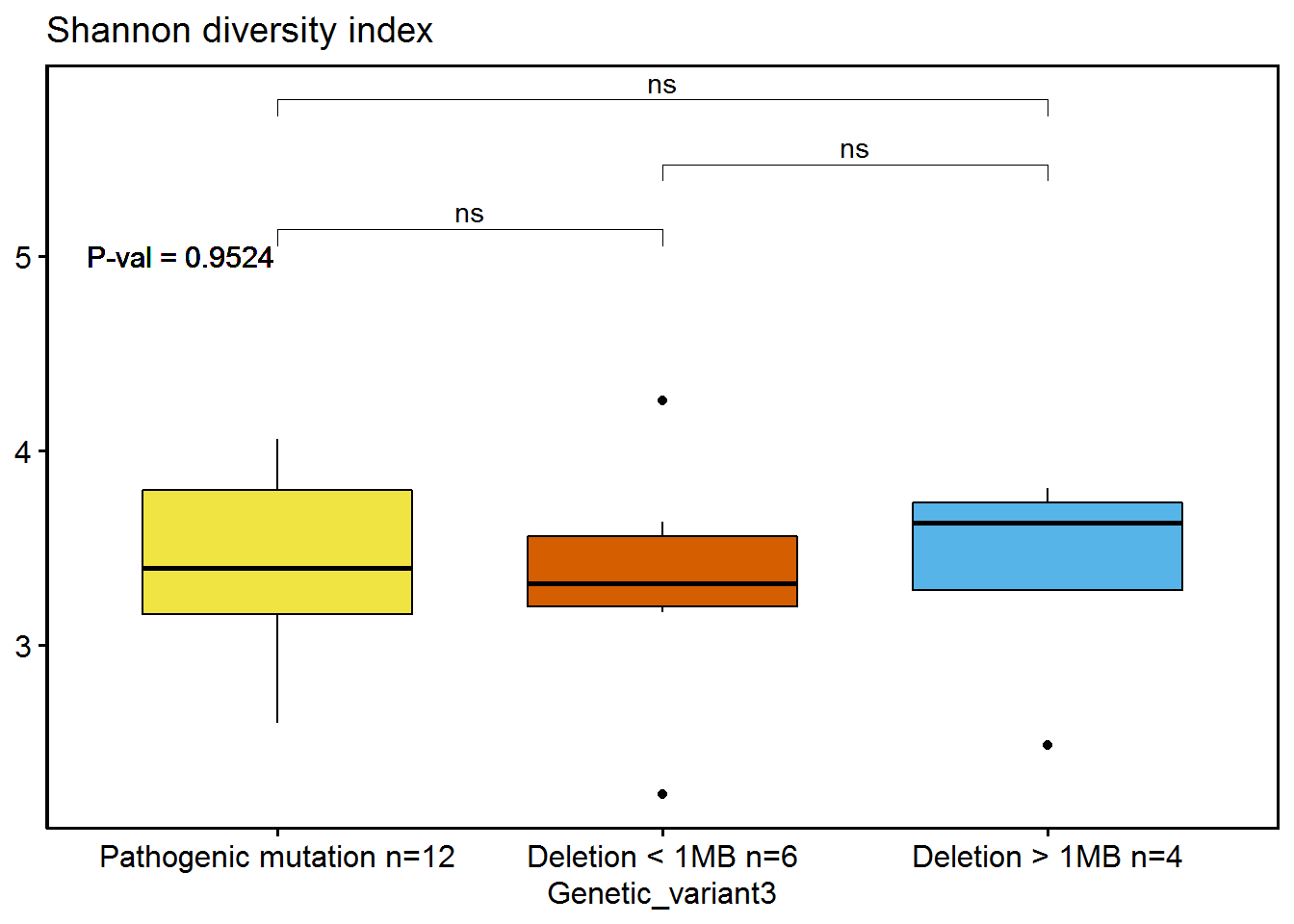

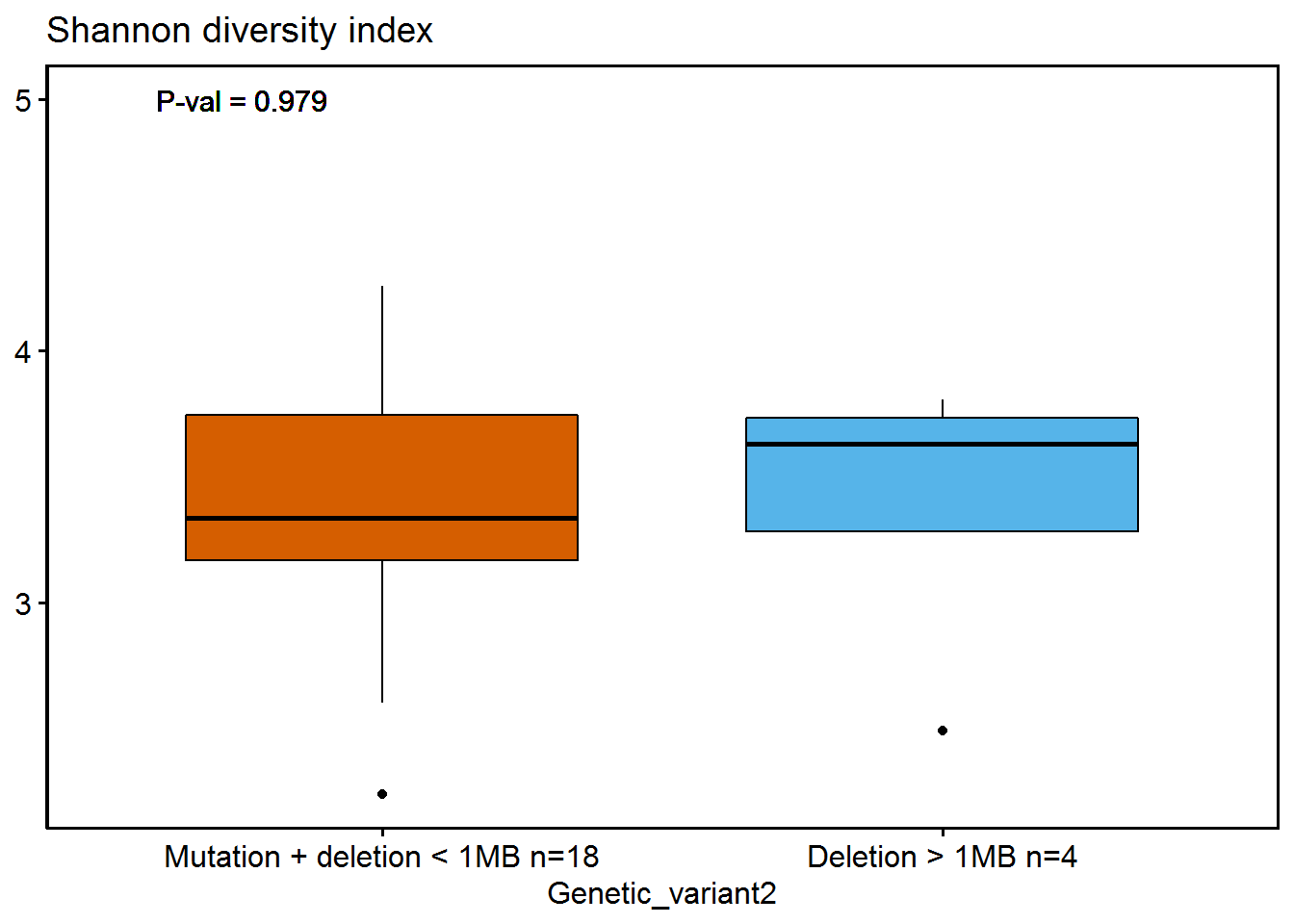

*Beta-diversity

Patients versus parents*

**Figure S3. CAP plot on the effect of intestinal complaints on beta diversity, grouped by disease status and level of intestinal complaints.**

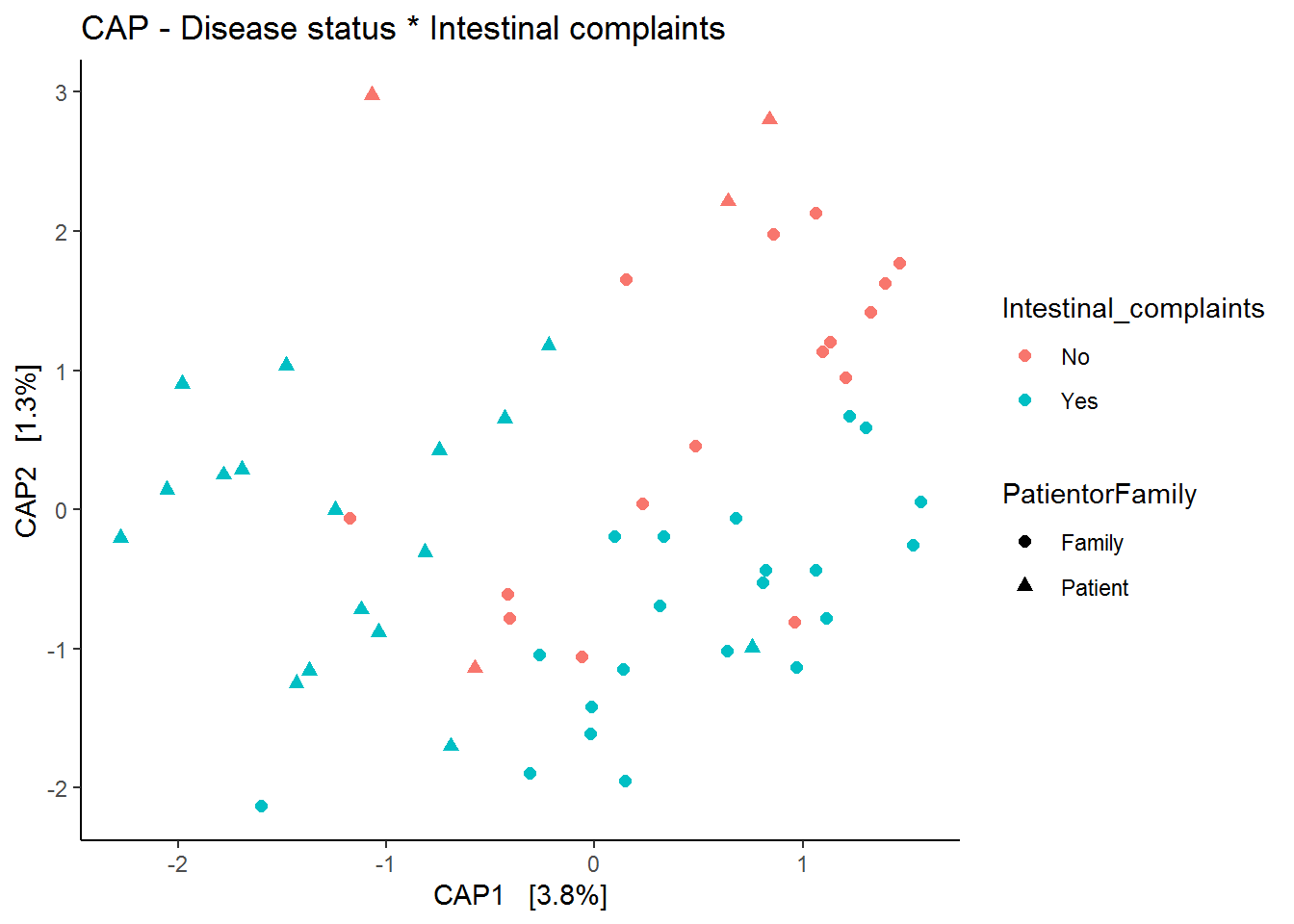

*Taxonomic results

Patients versus family*

For 15 genera we observed an FDR-uncorrected effect of disease status, of which seven genera are part of the *Lachnospiraceae* family, three of the *Ruminococcaceae* family, two of the *Clostridiales vadinBB60 group* family, and one genus was found for other families namely *Peptococcus, Rikenellaceae* and *Eubacteriaceae*. Fourteen genera showed a decrease in relative abundance in the patients compared to their family members. One genus *Peptococcus* was increased in the patient group compared to the family members, see **Figure S4.

Figure S4.** **Fifteen boxplots showing the main effect of disease status for the nominal significant genera.** The x-axis shows the patient vs family group including sample sizes in the format: the number of non-zero observations/total number of samples per group. The boxplot indicates the median (black line) and 25^th^ and 75^th^ quartiles as the outside of the box. The lines represent the largest and smallest values within the 1.5 interquartile range and the dots represents the outliers which is > 1.5 times and < 3 times the interquartile range.

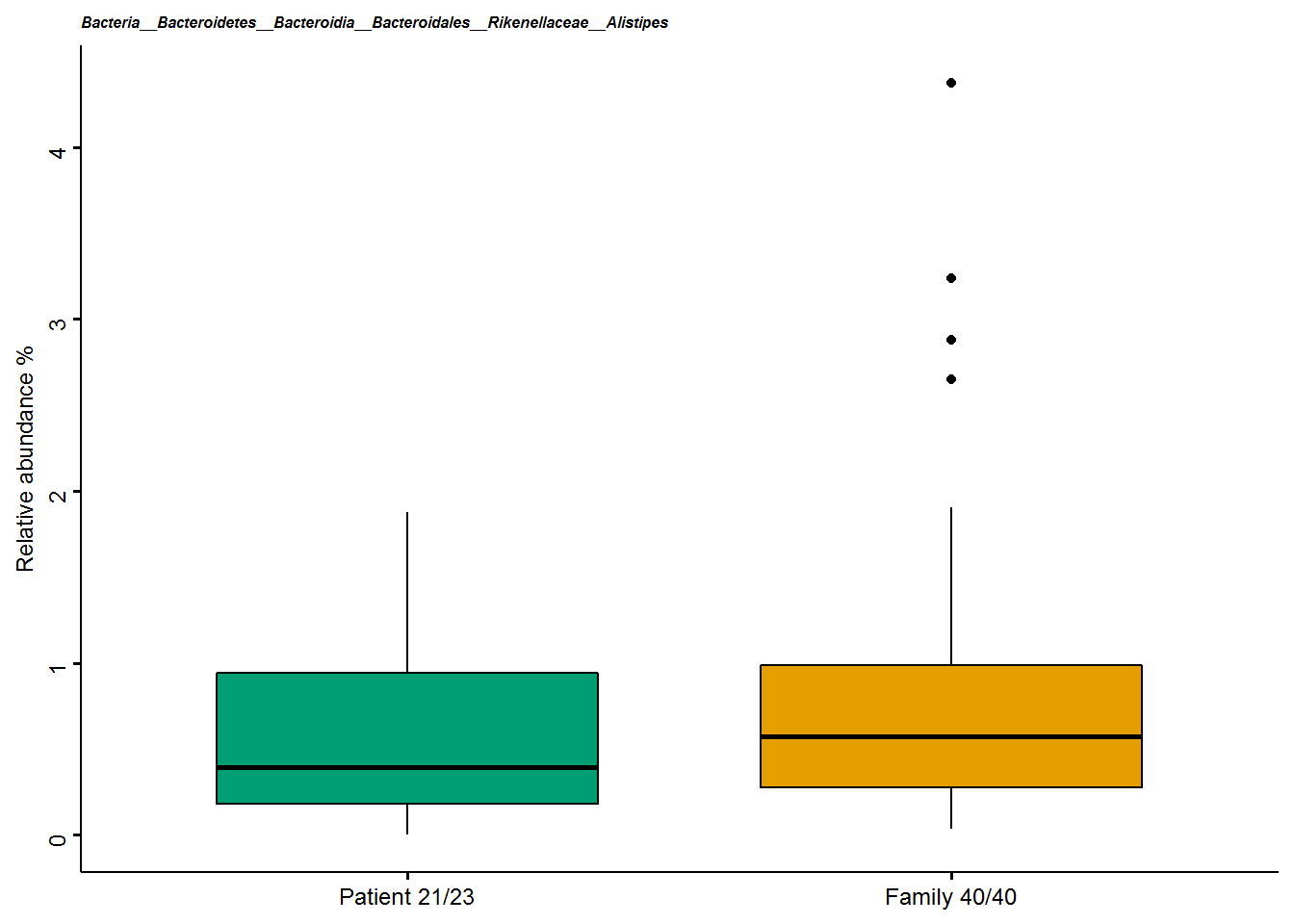

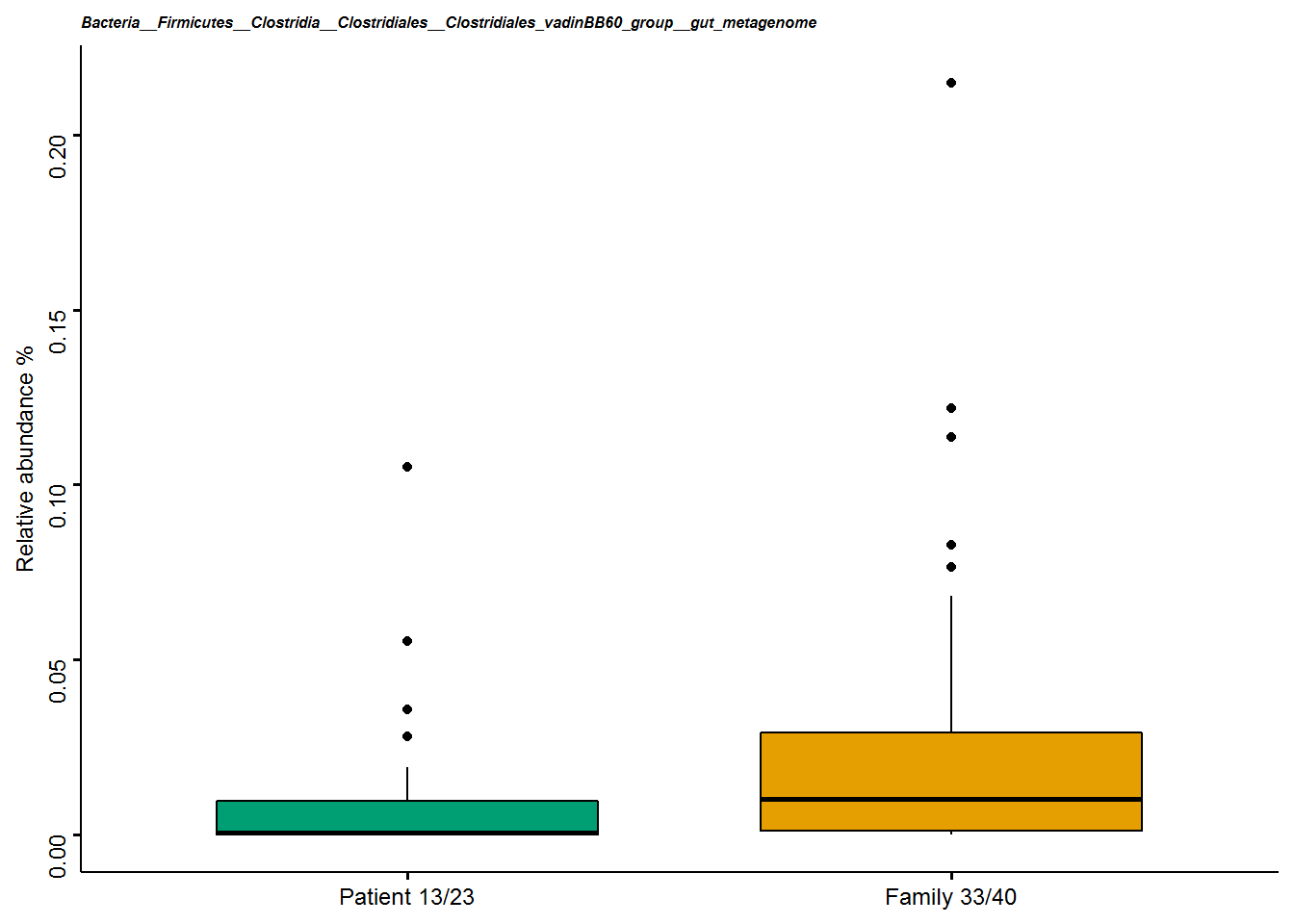

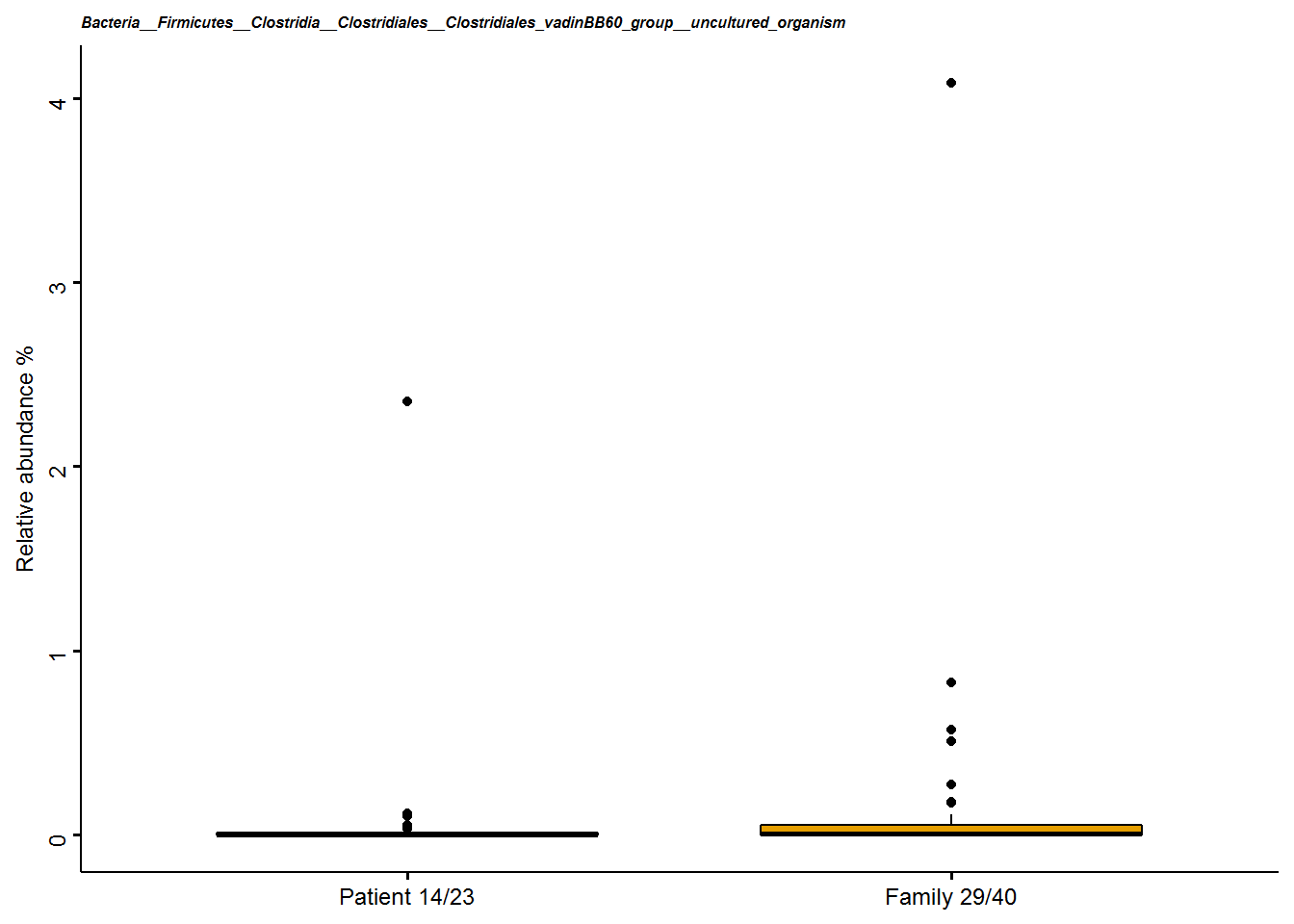

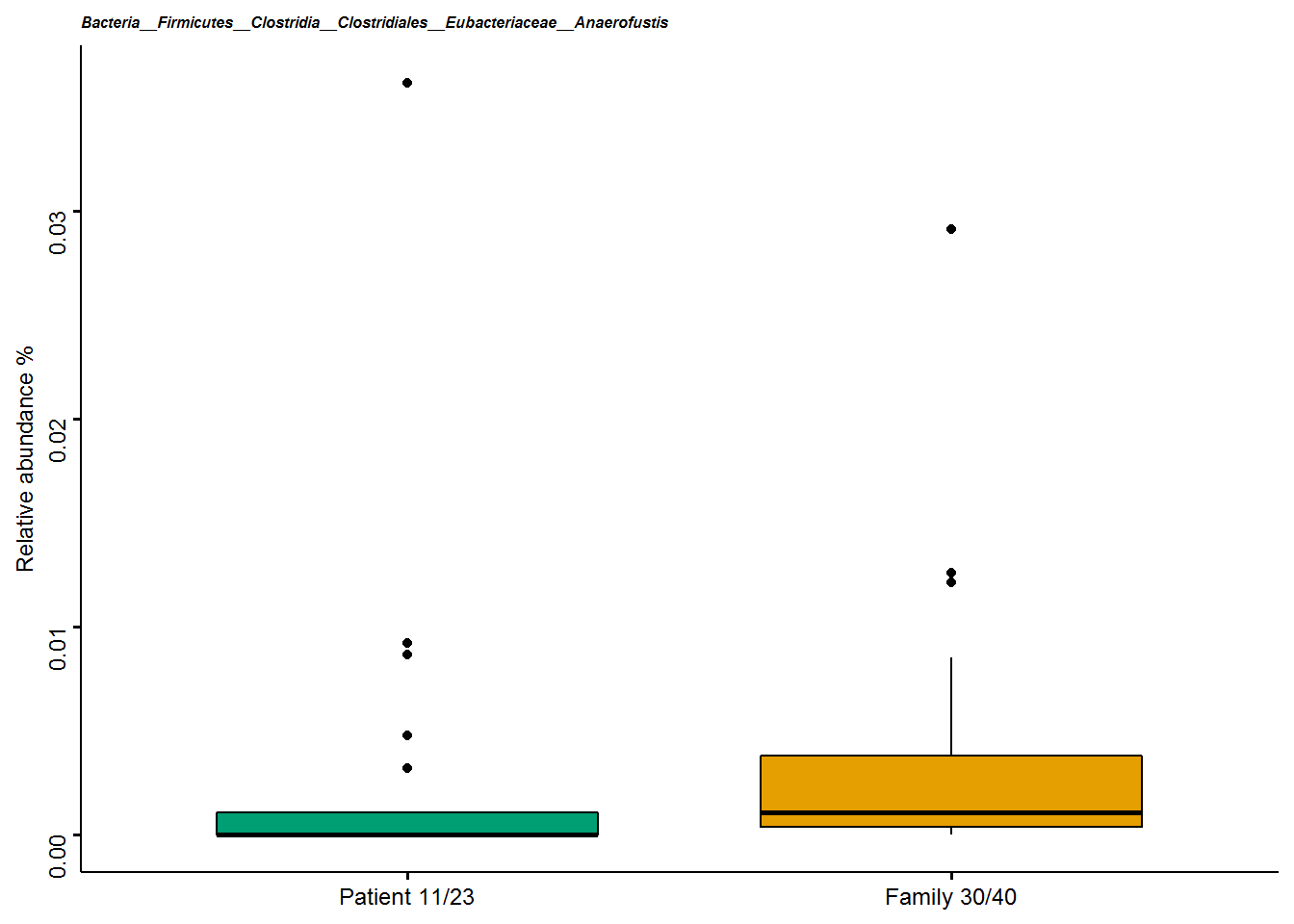

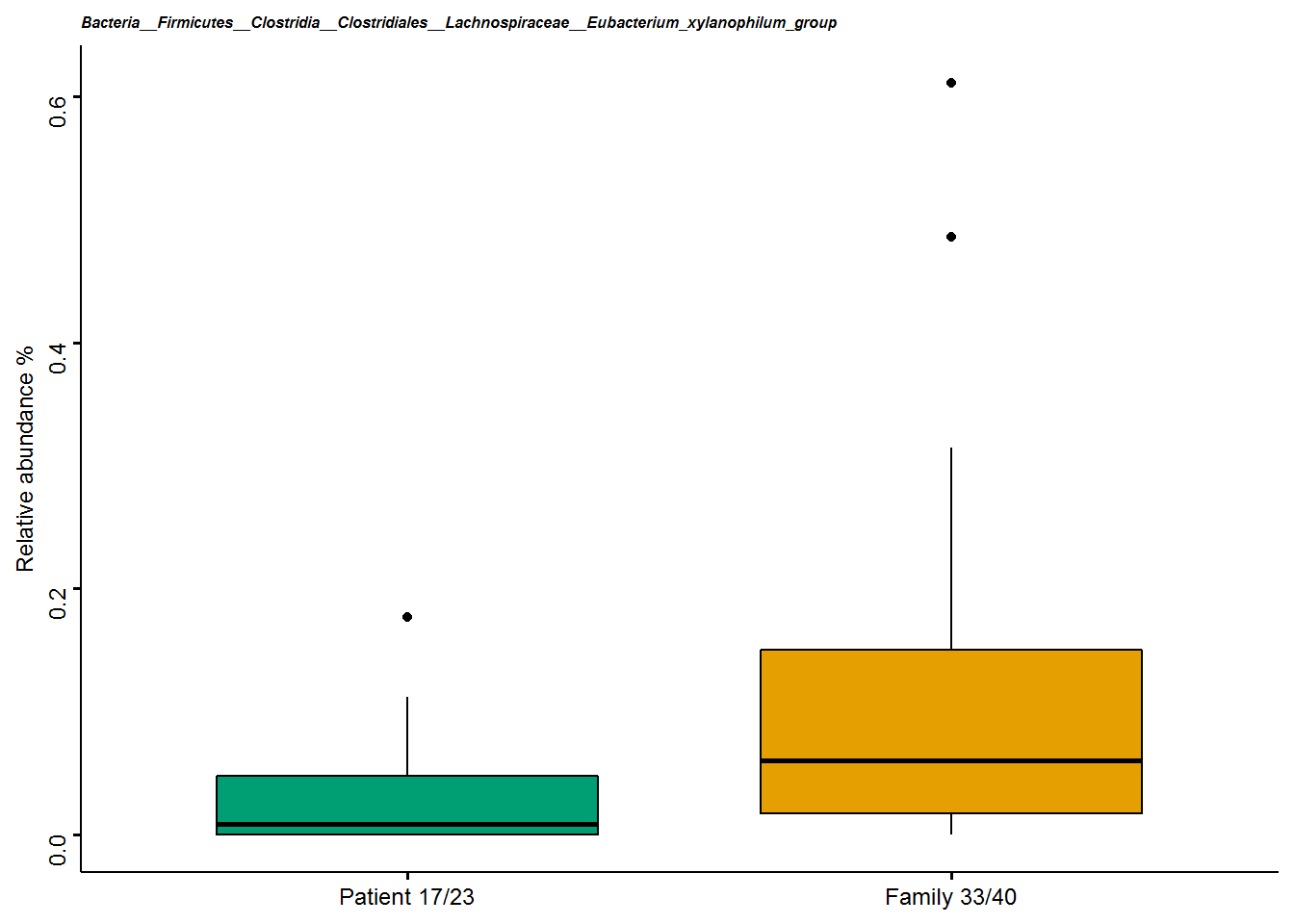

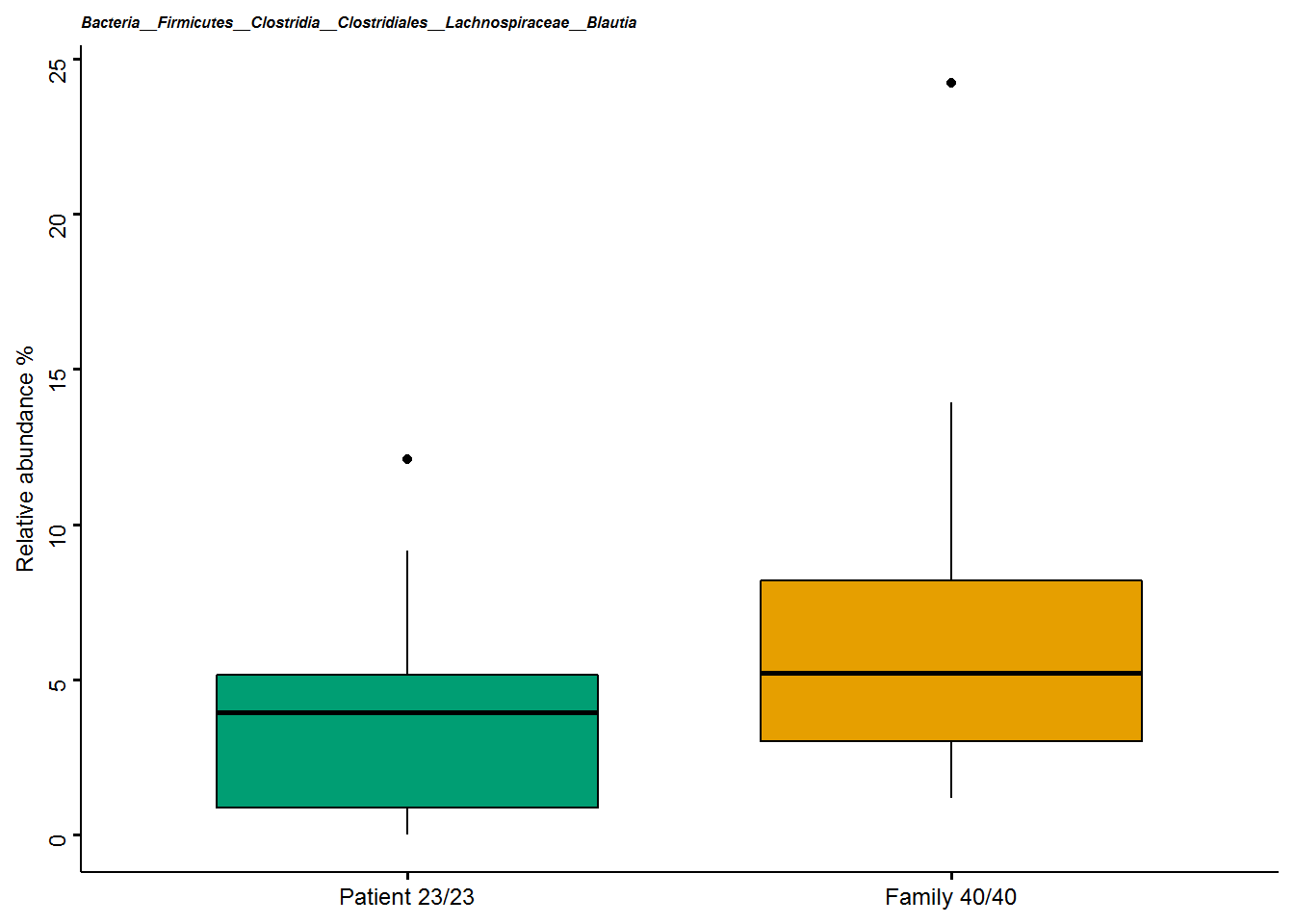

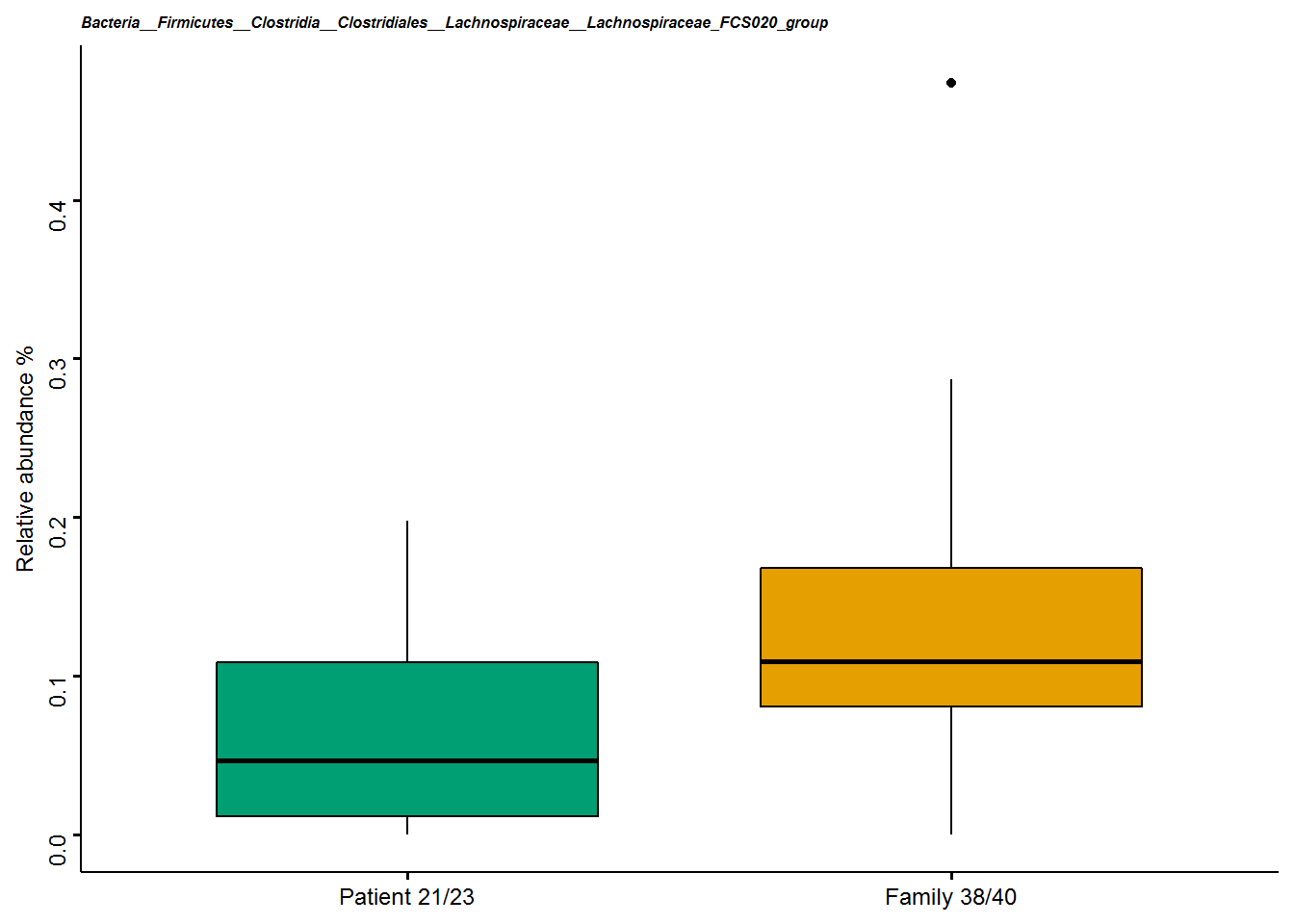

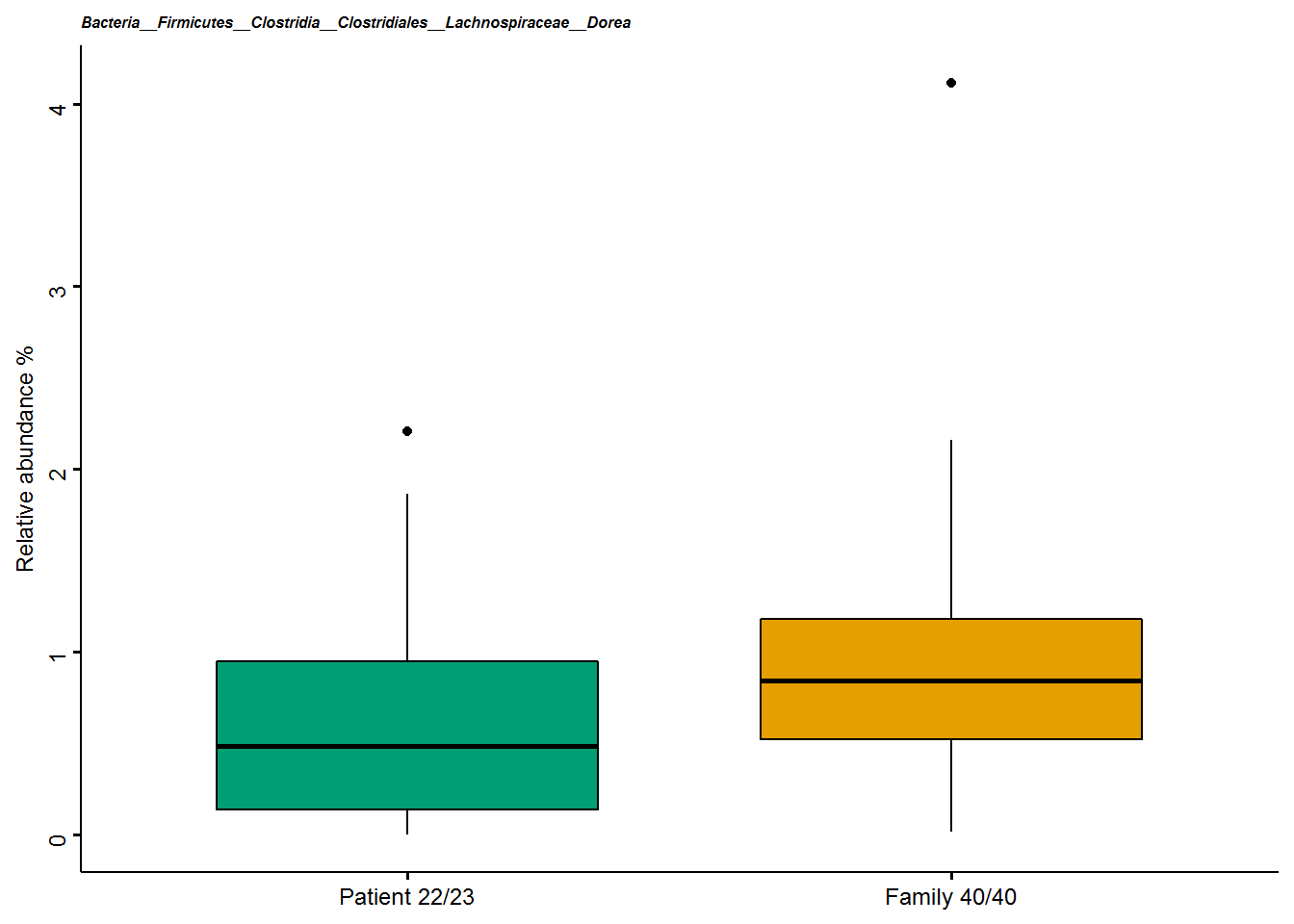

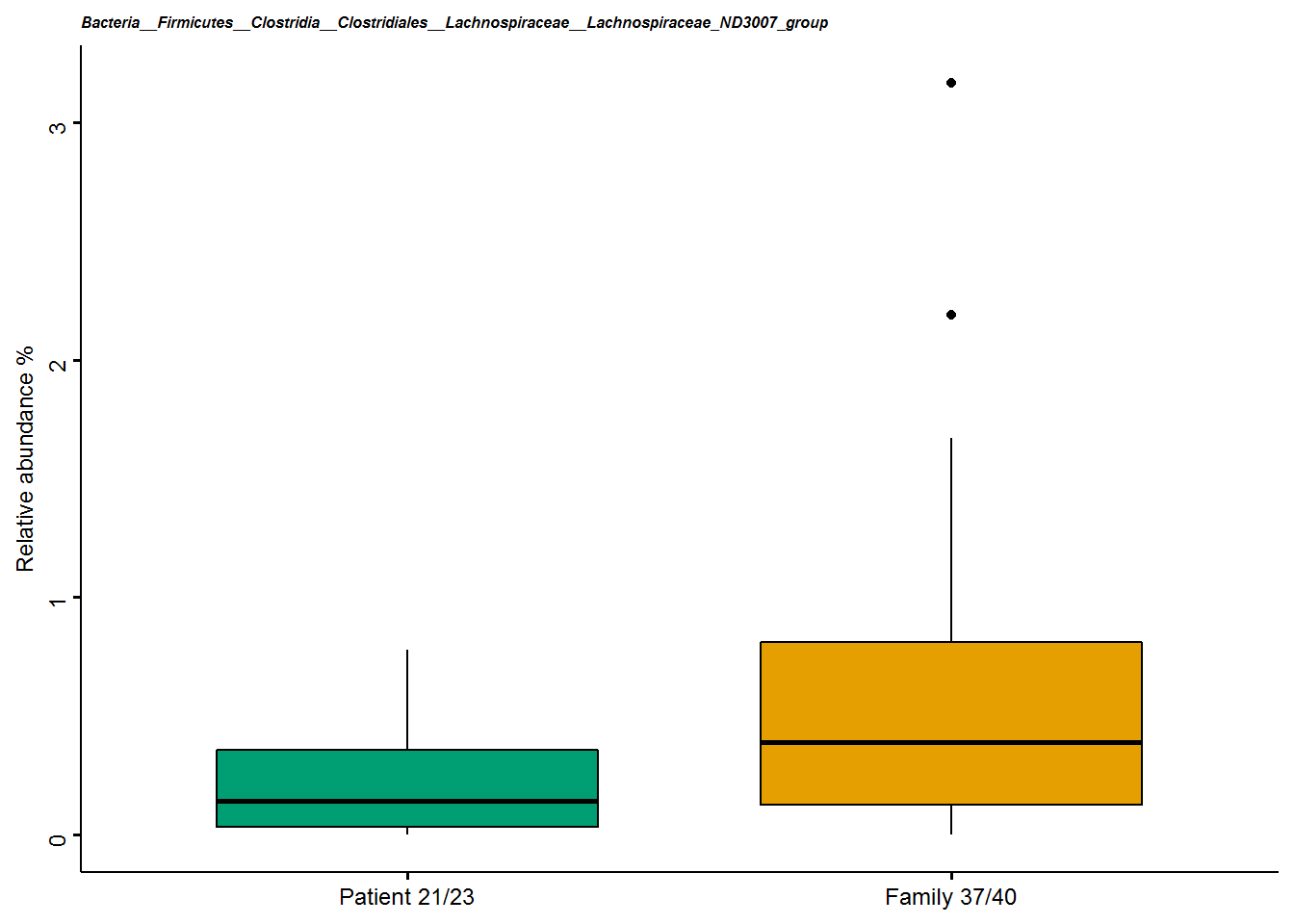

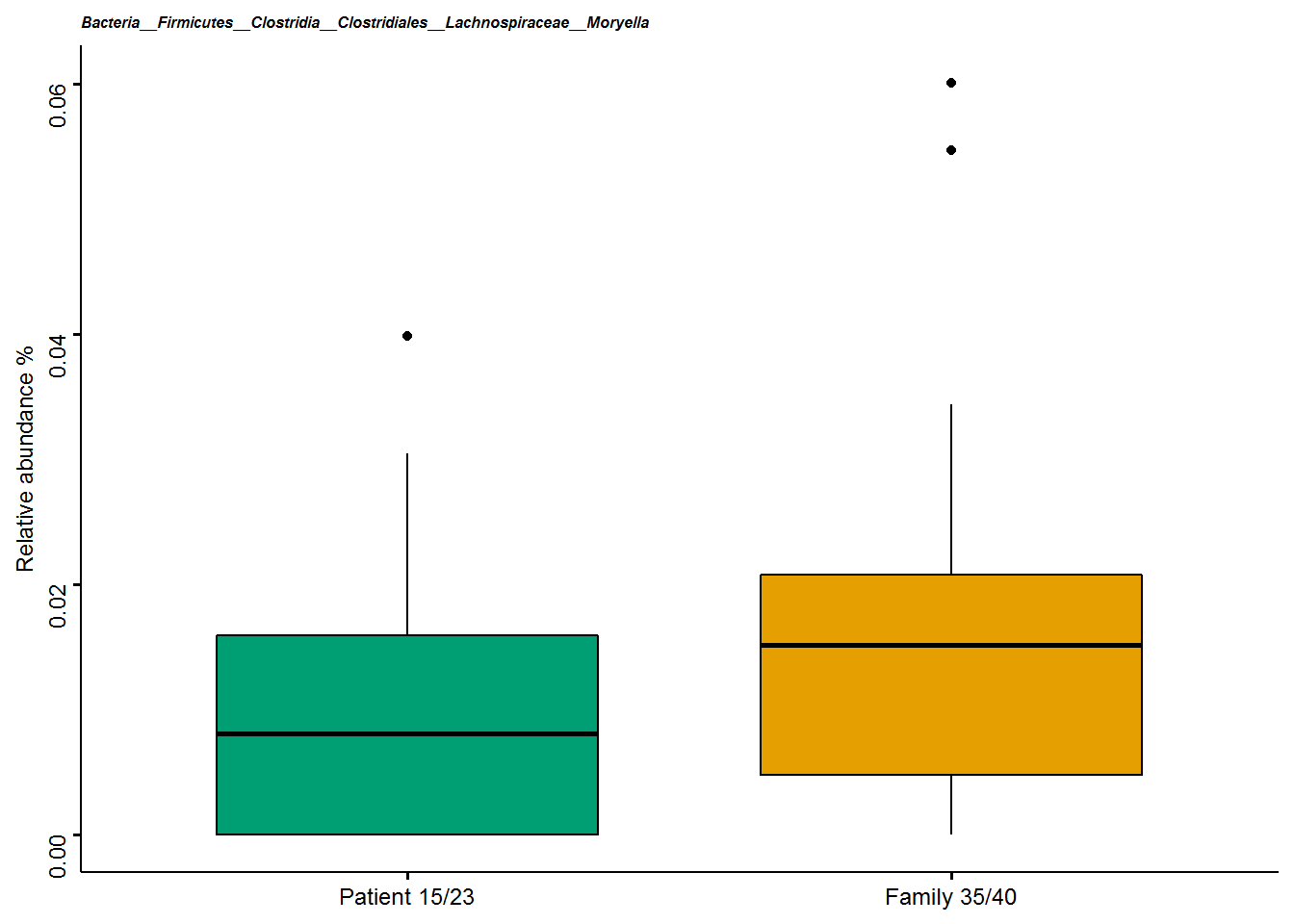

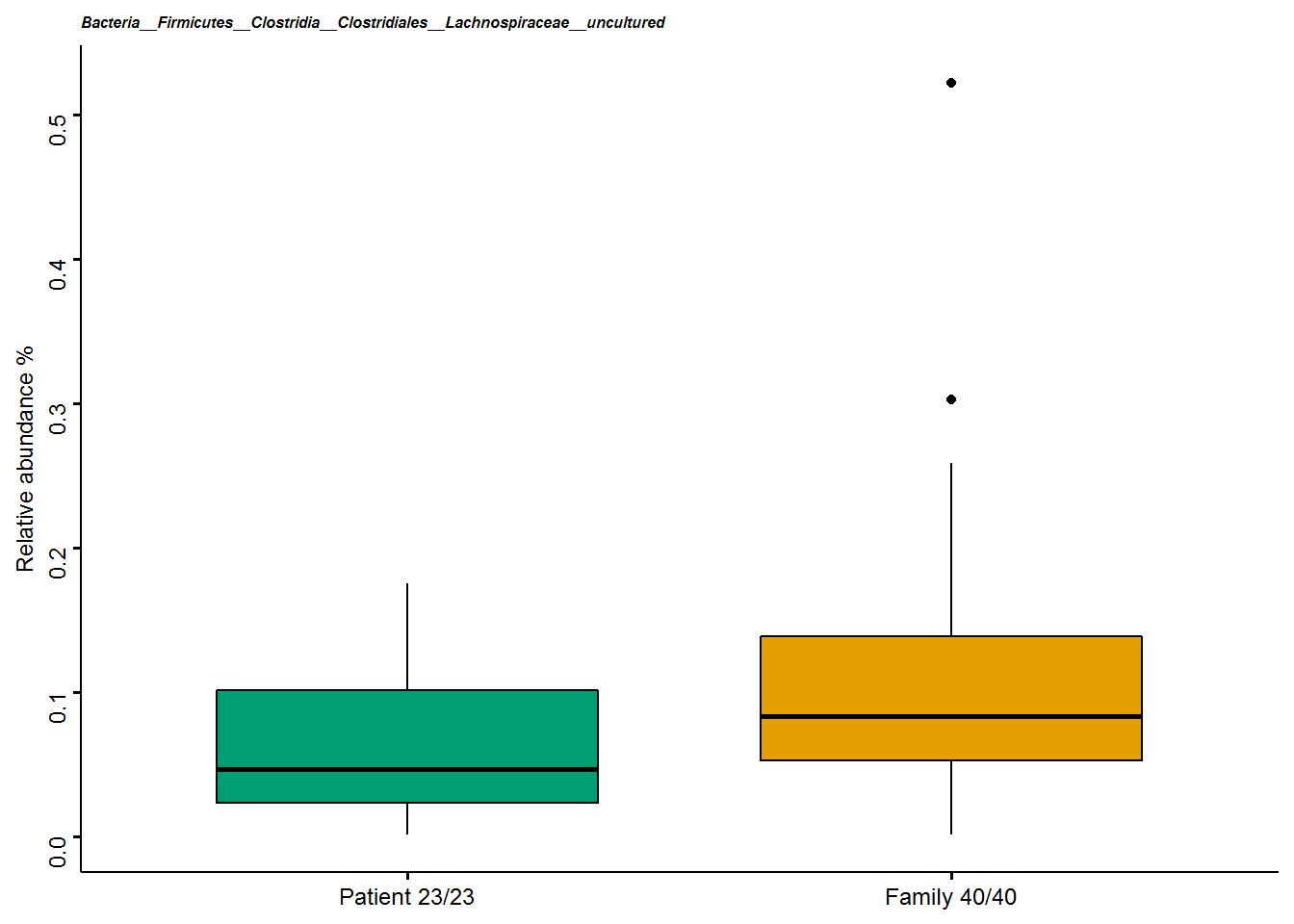

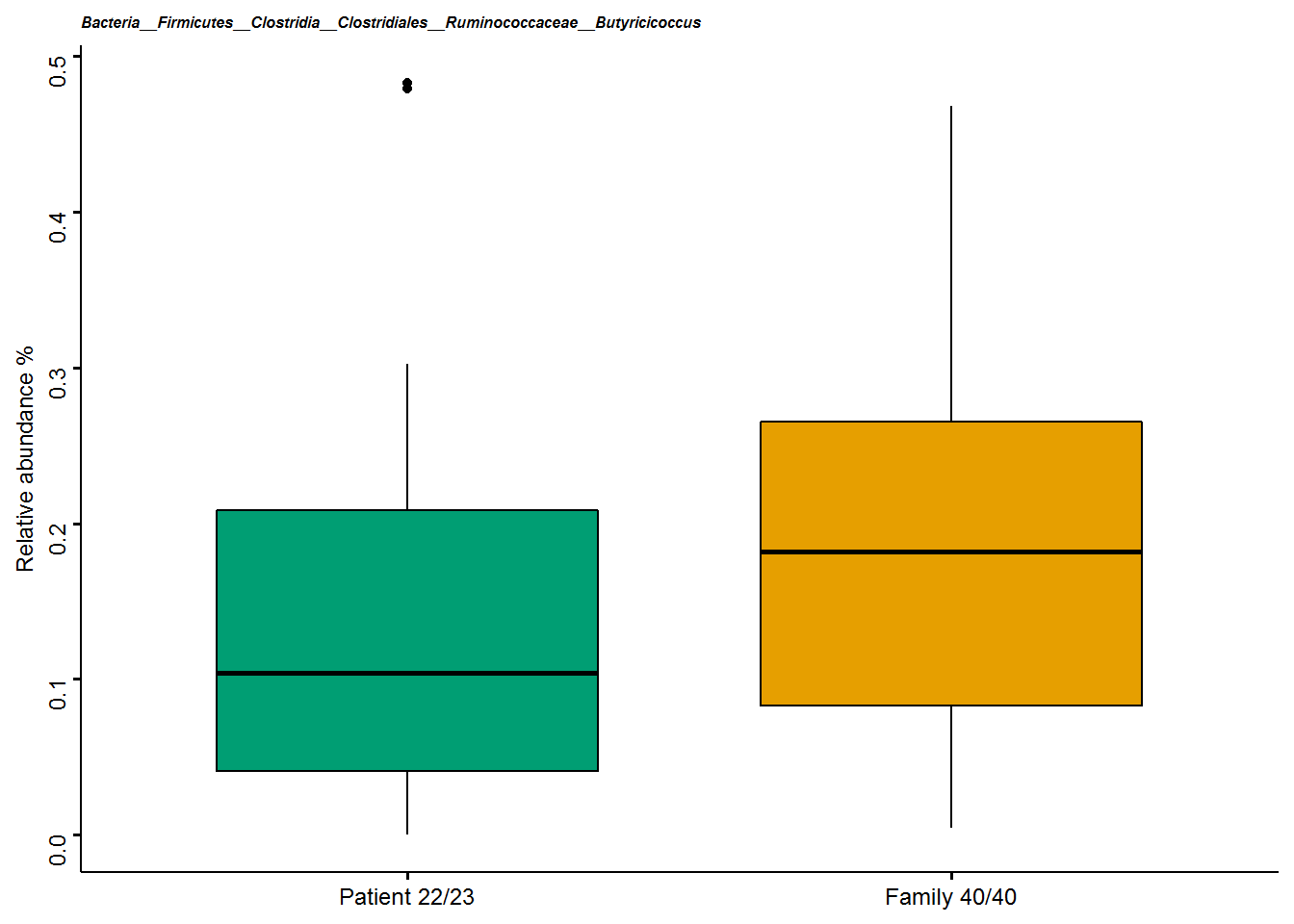

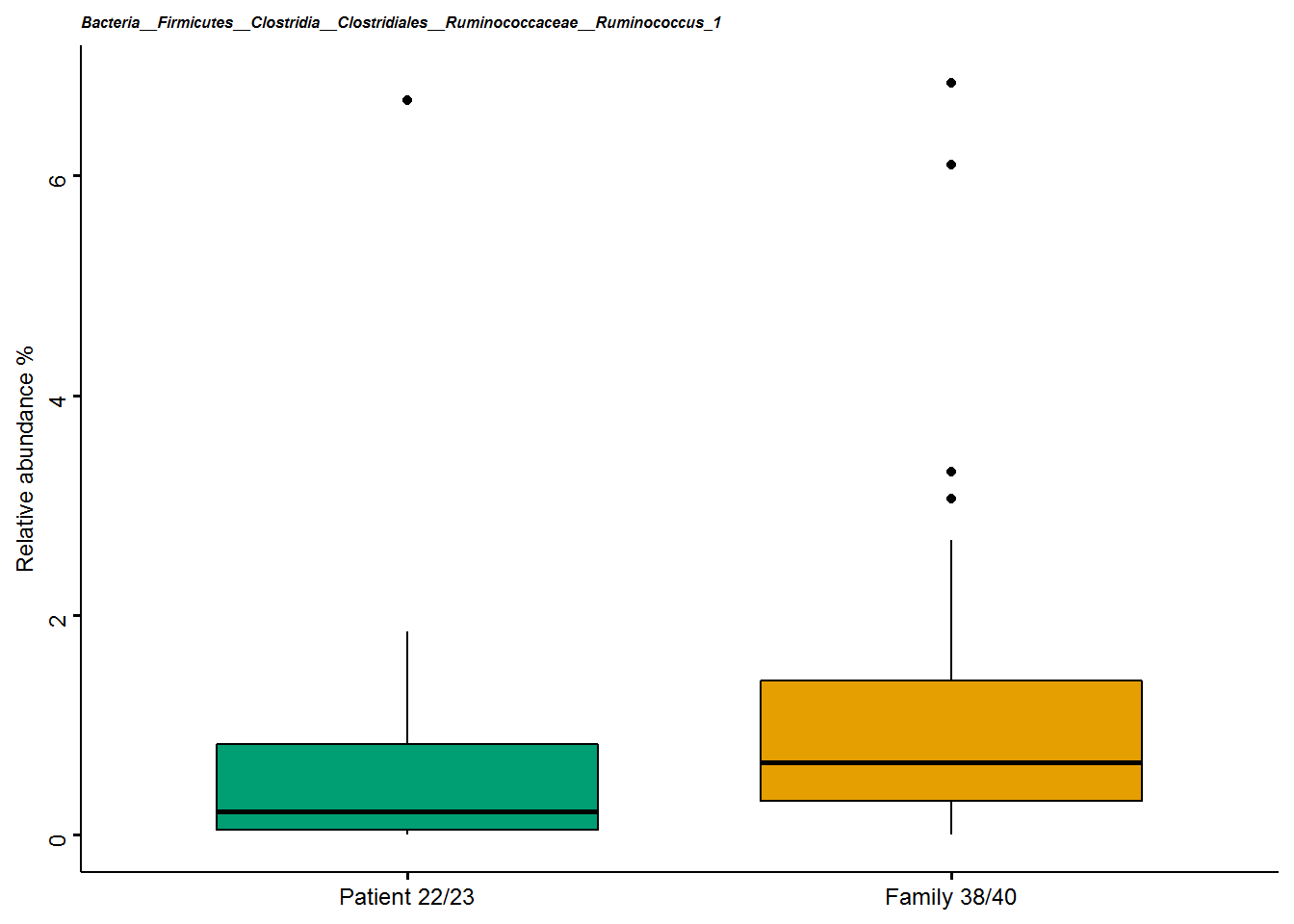

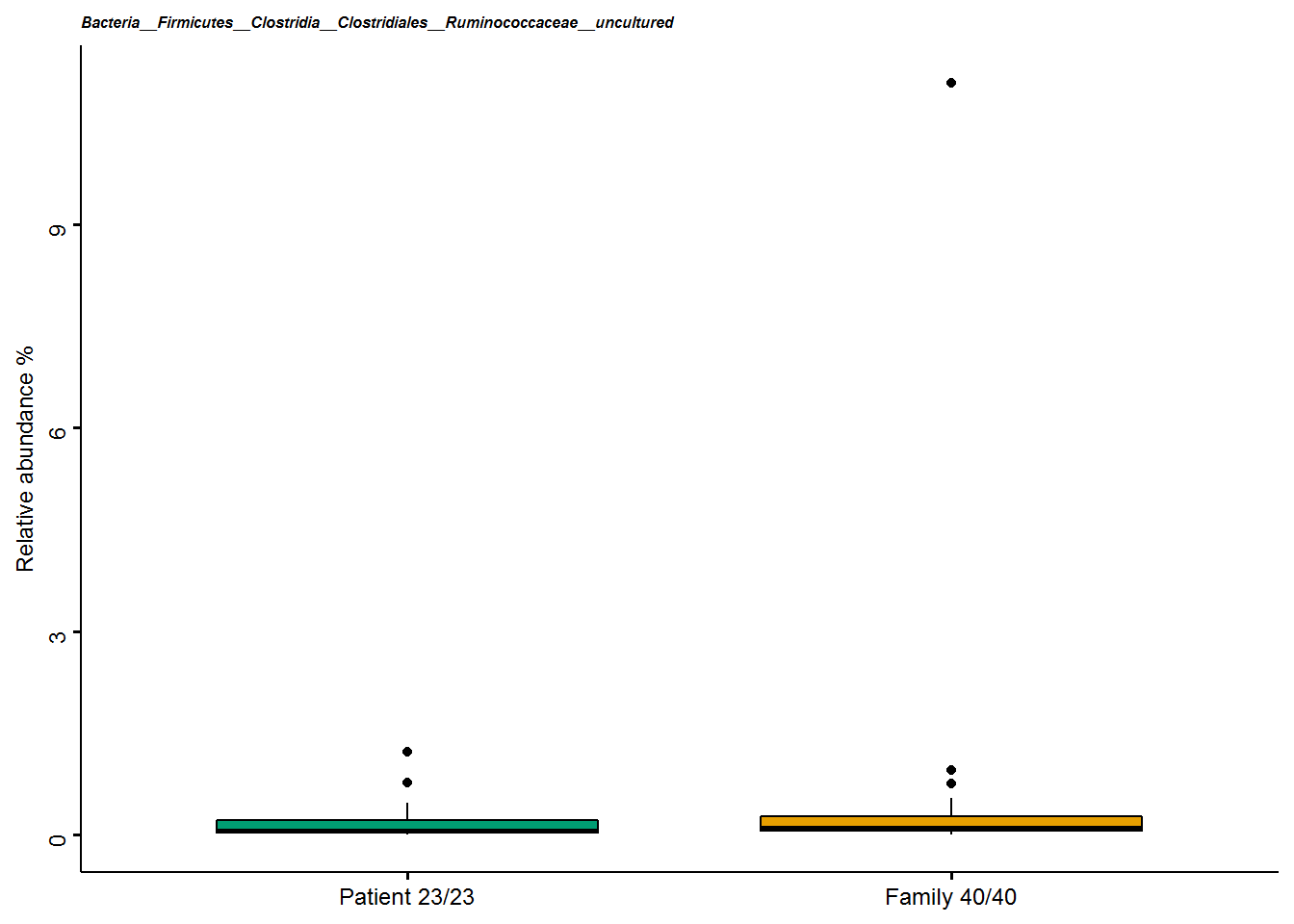

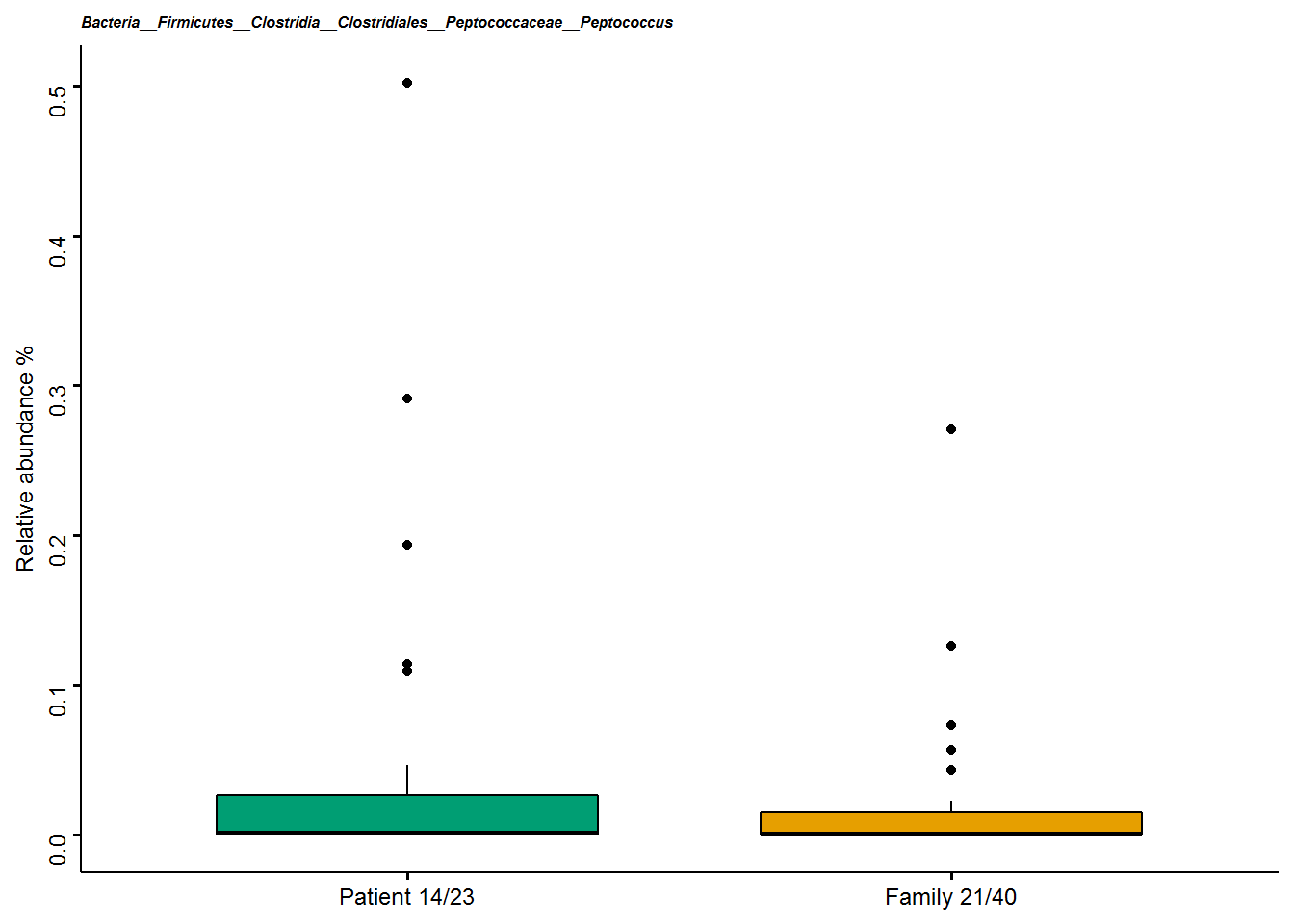

*Disease status * intestinal complaints*

For seventeen genera, we observed an FDR un-corrected interaction effect of disease status*intestinal complaints. Of these, four genera are part of the *Ruminococcaceae* family, three genera are part of the *Erysipelotrichaceae* family, two genera are part of the *Lachnospiraceae* family, and one of each genus belonged to other families namely *Eggerthellaceae, Bacteroidaceae, Peptostreptococcaceae, Rikenellaceae, Clostridiaceaea 1, Eubacteriaceae, Veillonellaceae* and *Pseudomonadaceae*. Nine genera (*Gordonibacter, Bacteroides, Alistipes, Eubacterium fissicaneta group, Eisenbergiella, Acetanaerobacterium, Ruminococcaceae-UBA1819, Clostridium innocuum group* and *Holdemania)* showed the same pattern as *Merdibacter:* in those participants reporting no intestinal complaints relative abundance was higher in the patient group compared to their family members, and in those participants reporting intestinal complaints, lower relative abundance was observed in the patient group compared to their family members. Three genera (*Ruminococcaceae UCG003, Pseudomas* and *Romboutsia*) showed the reverse pattern: when not experiencing intestinal complaints relative abundance is lower in patients compared to their family members. Those participants reporting intestinal complaints displayed higher relative abundance in patients compared to their family members. For five genera the contrasting pattern between disease status and intestinal complaints was less apparent. In two of these (*Anaerofustis* and *Ruminococcus 1*) the difference in lower relative abundance in patients versus family members is strongest in those reporting versus not reporting intestinal complaints. In three of these (*Dialister*, C*lostridium sensu stricto 1* and *Turicibacter*), the difference in relative abundance in patients versus family members is strongest in those not reporting versus reporting intestinal complaints. The plots in **Figure S5** are sorted based on this visual pattern. It must be noted that of these seventeen genera, only *Bacteroides*, *Alistipes*, *Acetanaerobacterium, Ruminococcaceae UCG003,* *Anaerofustis* and *Ruminococcus 1* show a significant difference based on disease status in any of the two intestinal complaints groups.

**Figure S5.** **Seventeen boxplots showing the interaction effect of disease status*intestinal complaints for the nominal significant genera.** The x-axis shows the intestinal complaints group including sample sizes in the format: the number of non-zero observations/total number of samples per group. The boxplots are coloured by disease status and indicates the median (black line) and 25^th^ and 75^th^ quartiles as the outside of the box. The lines represent the largest and smallest values within the 1.5 interquartile range and the dots represents the outliers which is > 1.5 times and < 3 times the interquartile range. A Mann-Whitney U-test is performed to test the within effects showing the asterisks for the p-value ns= p>0.05, * p<0.05, ** p<0.01.

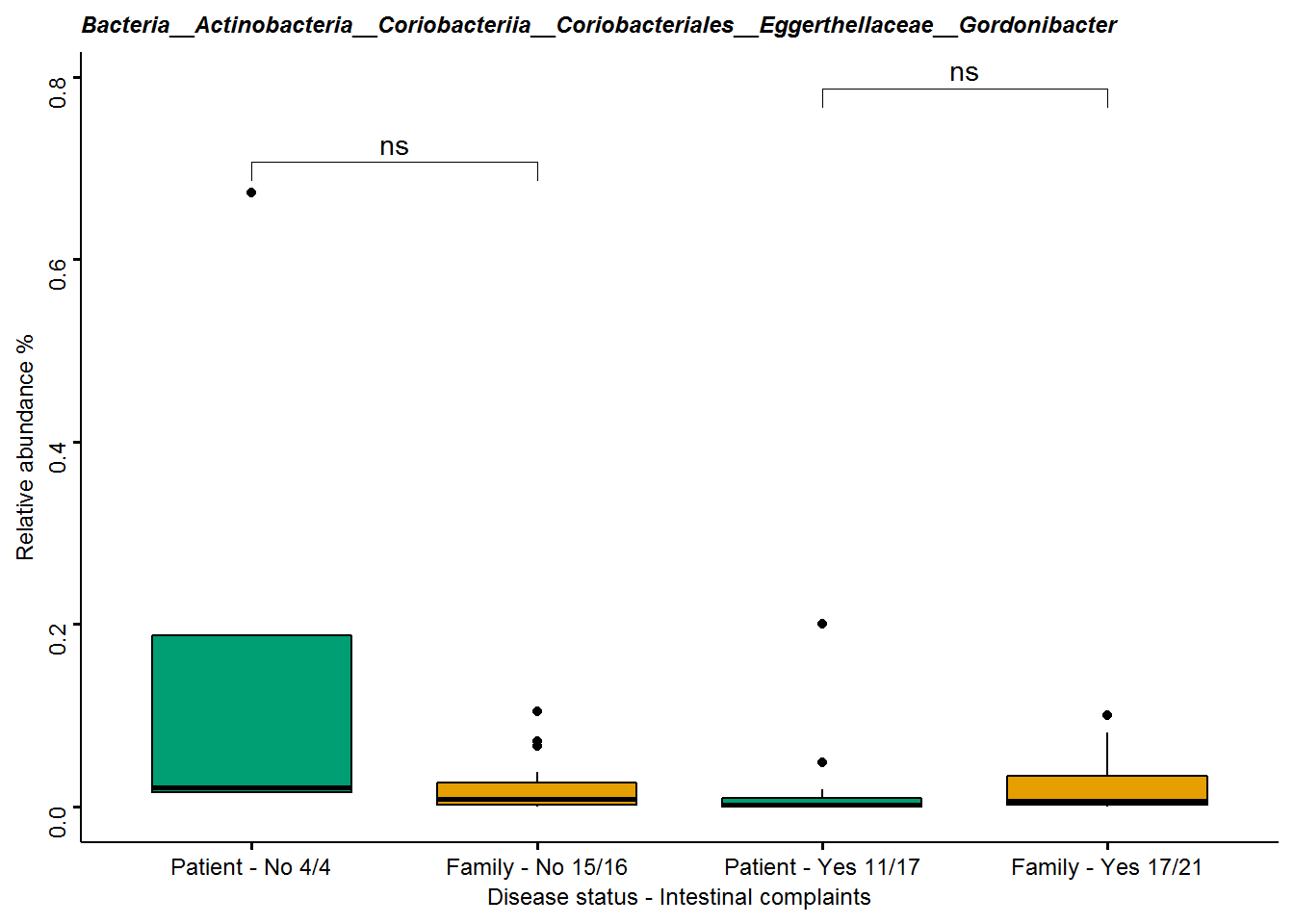

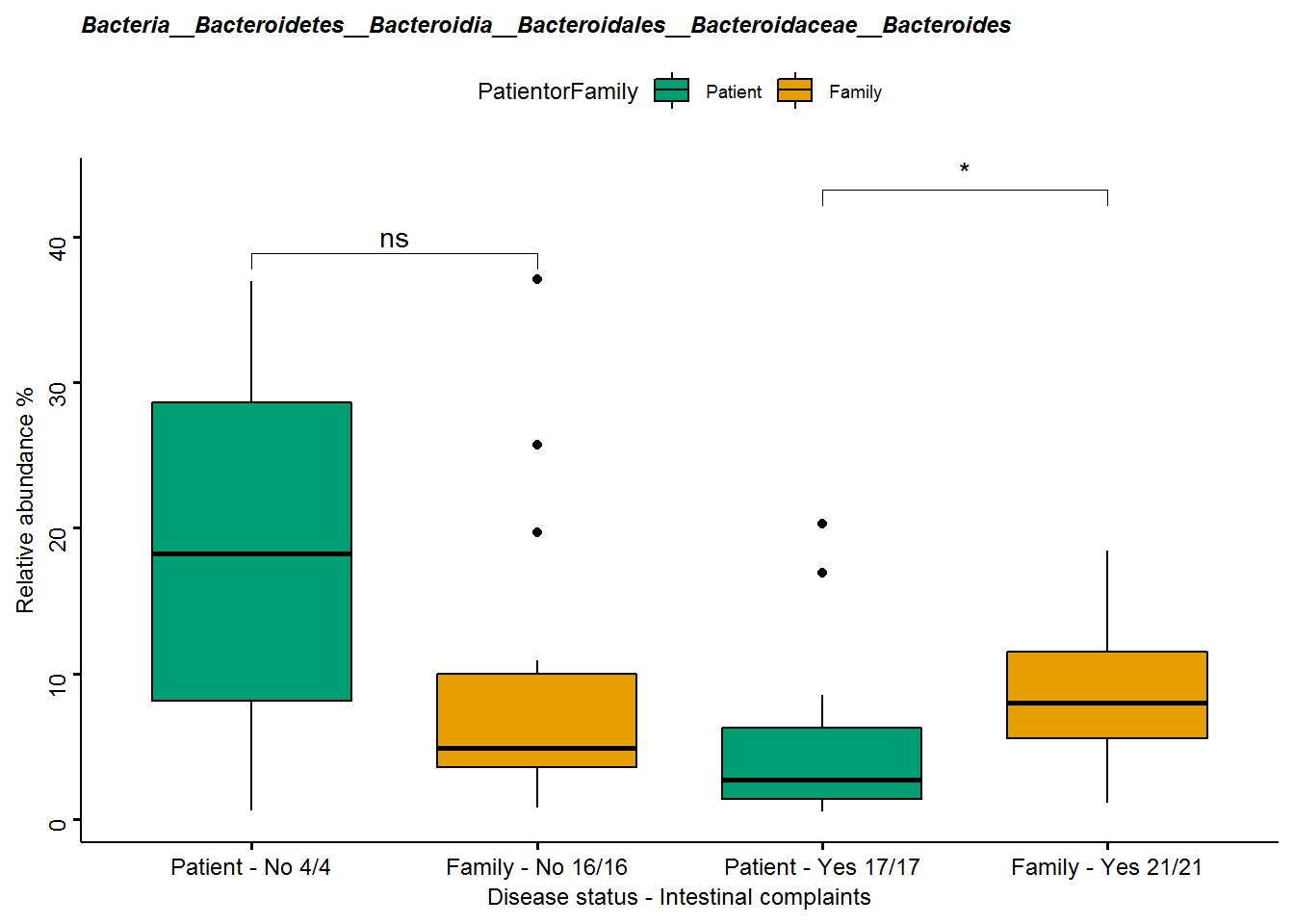

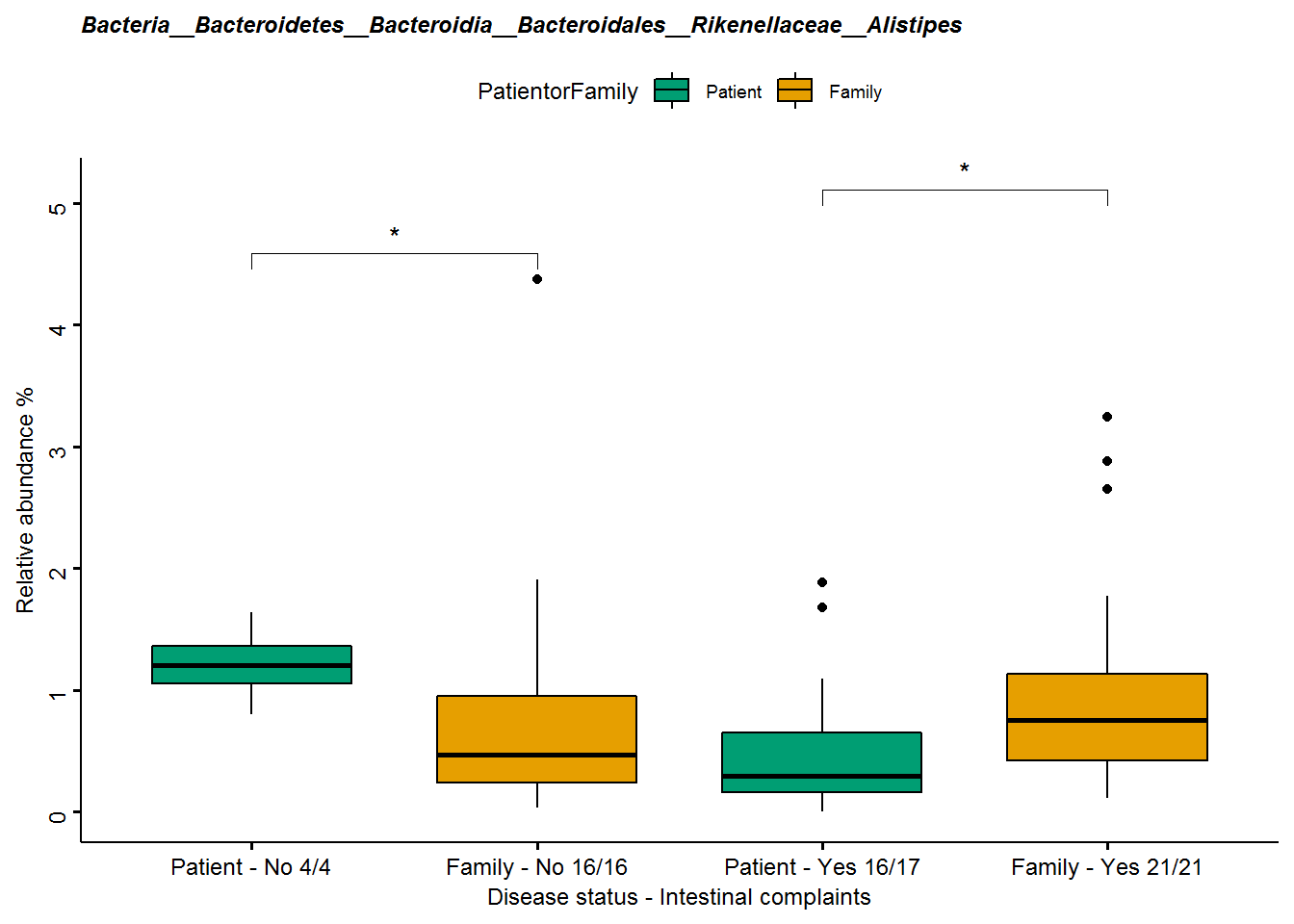

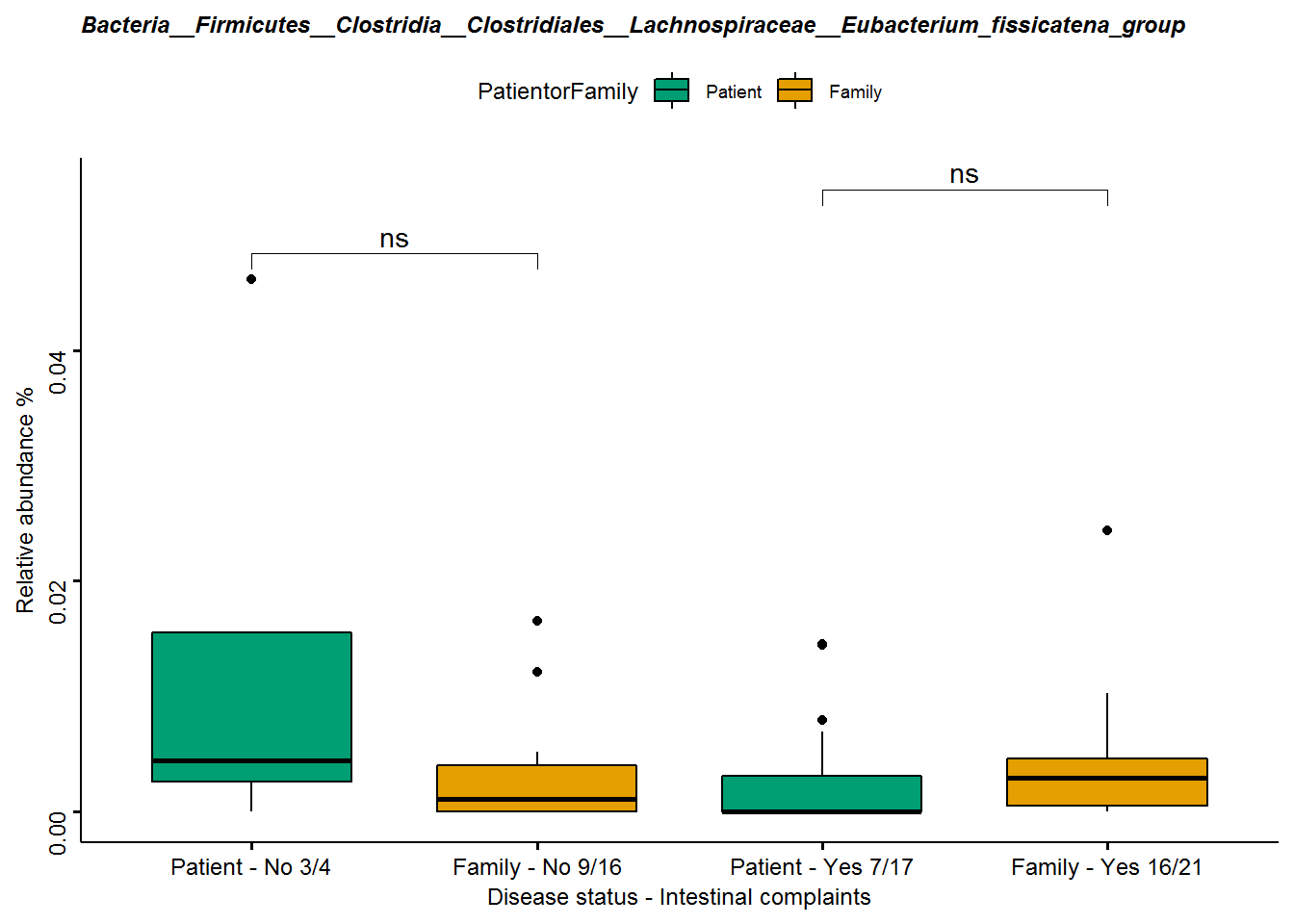

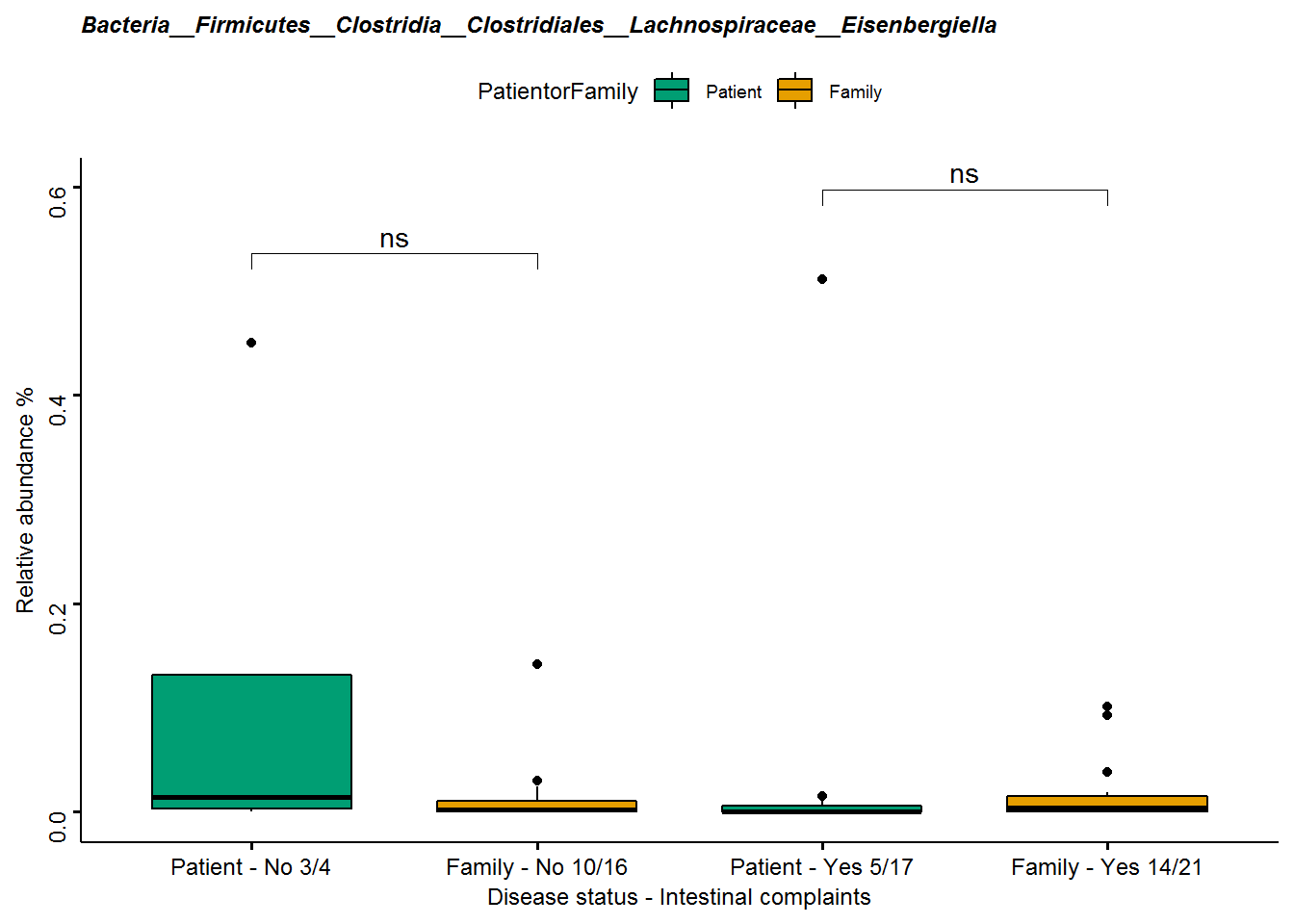

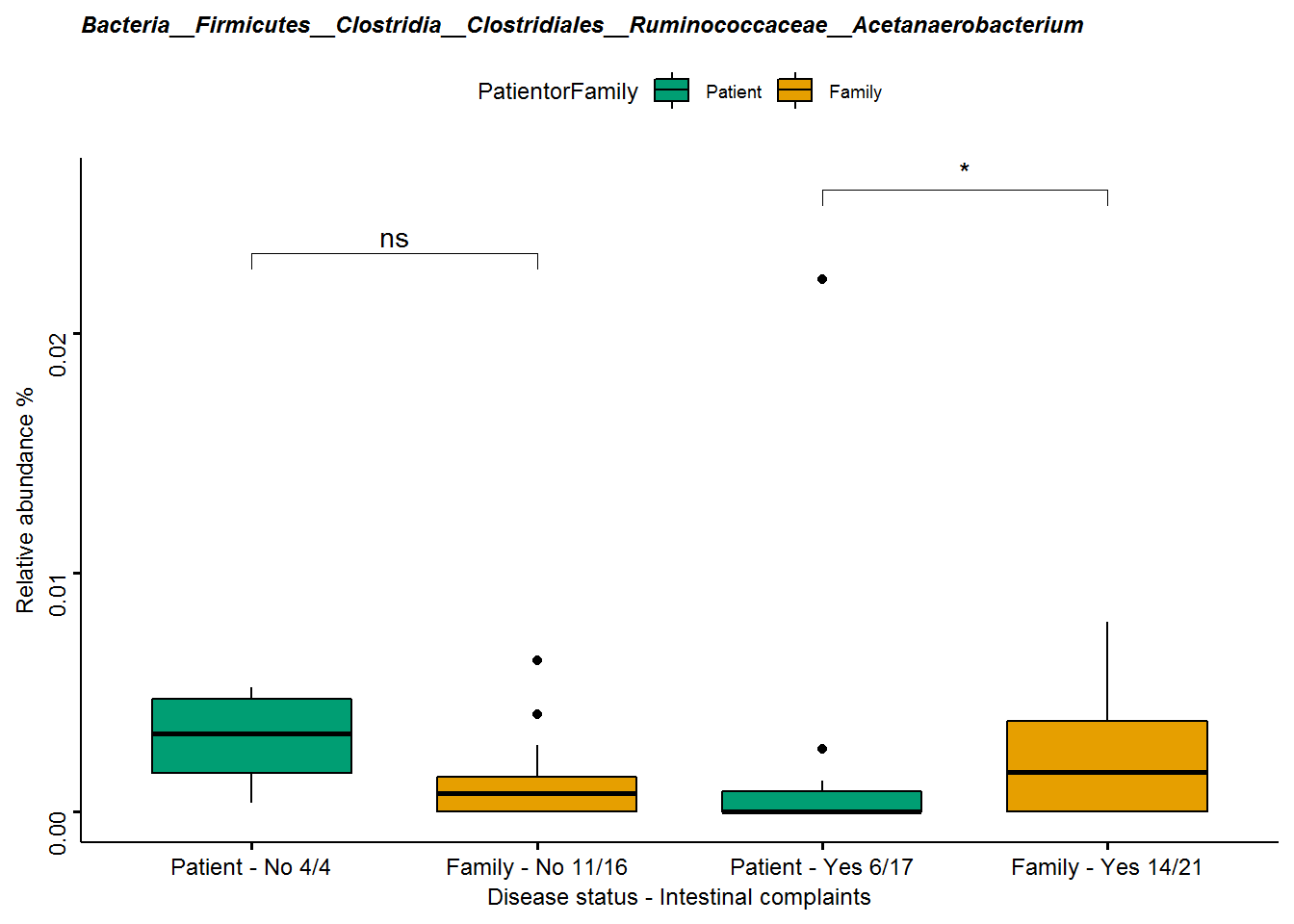

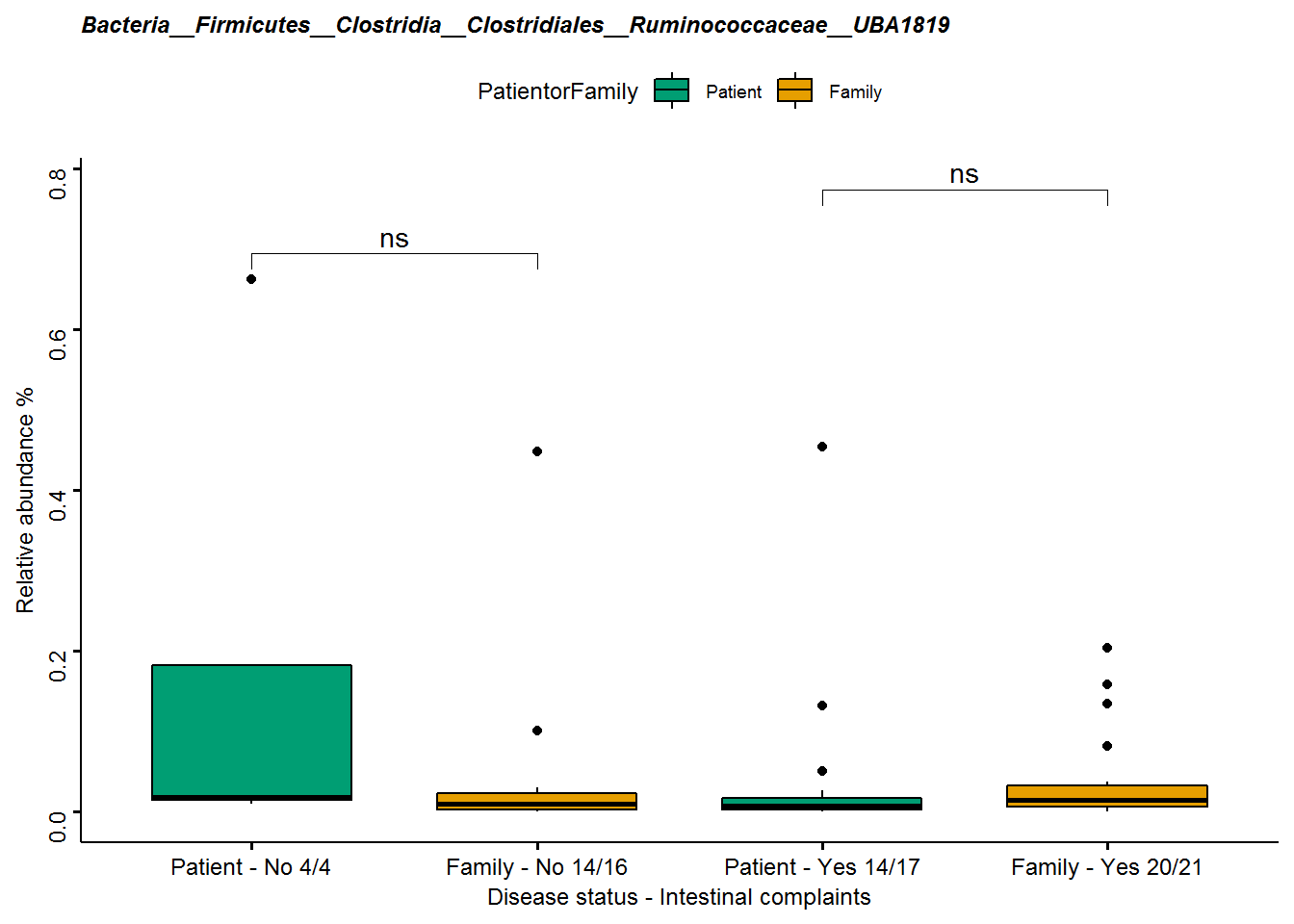

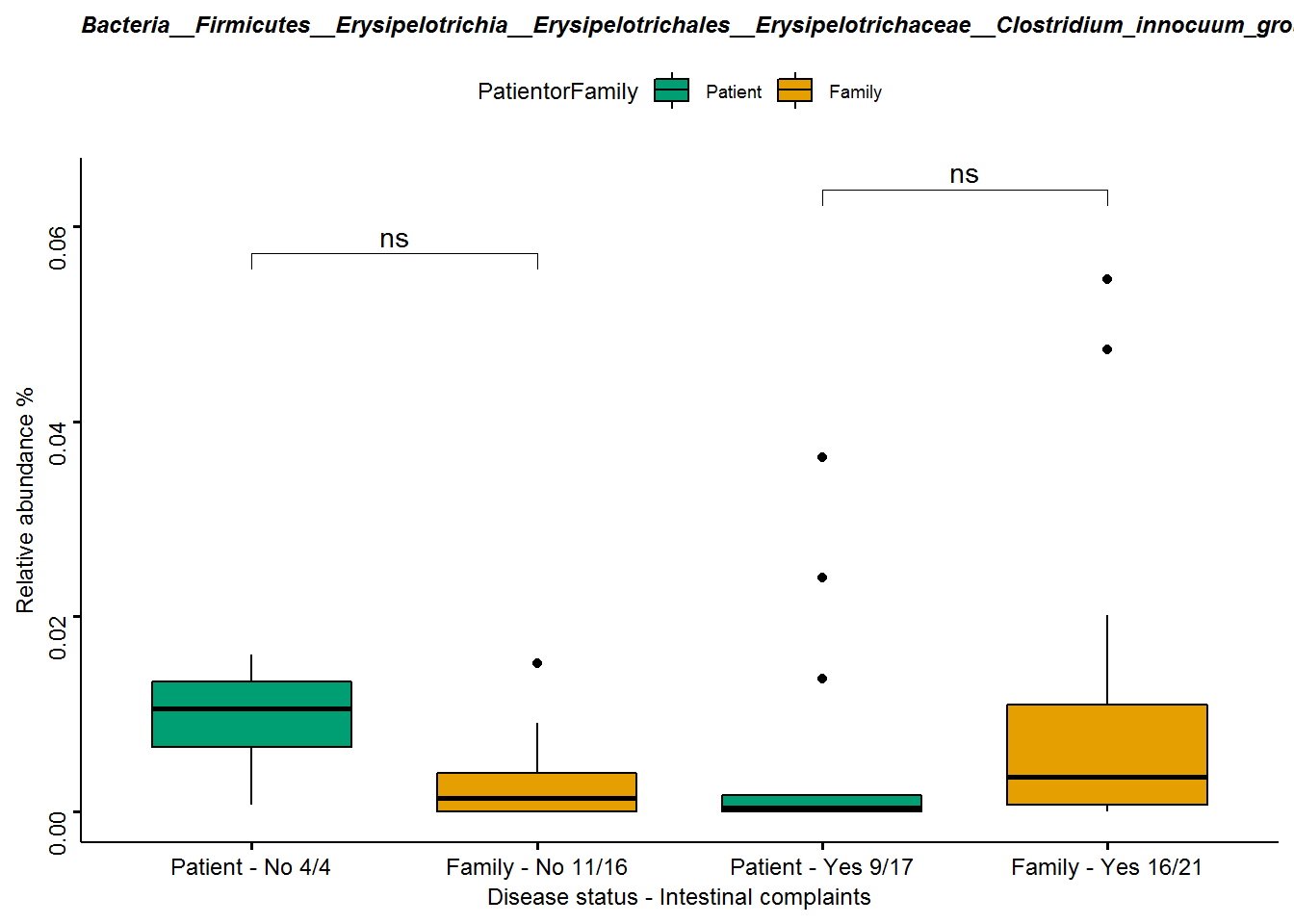

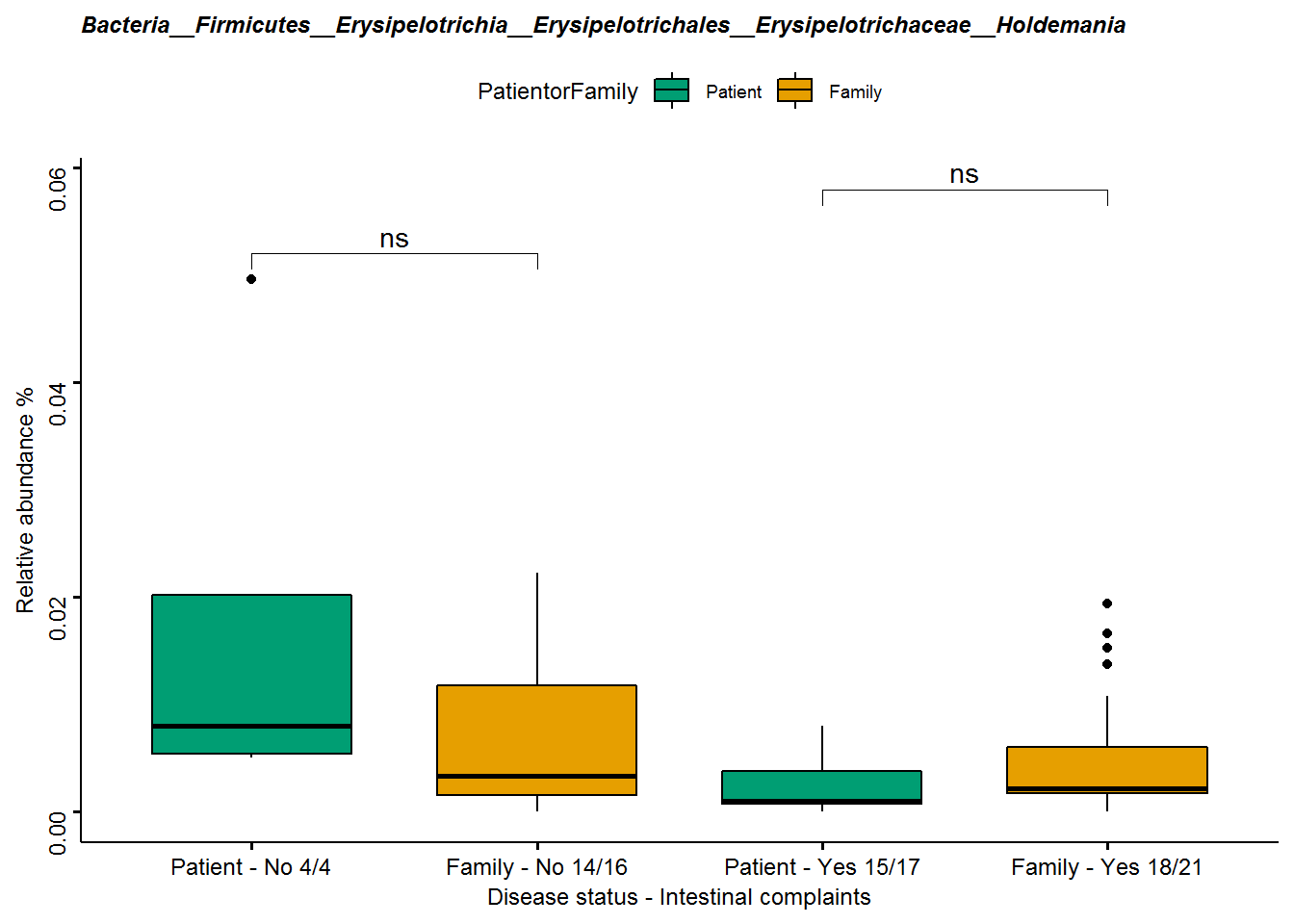

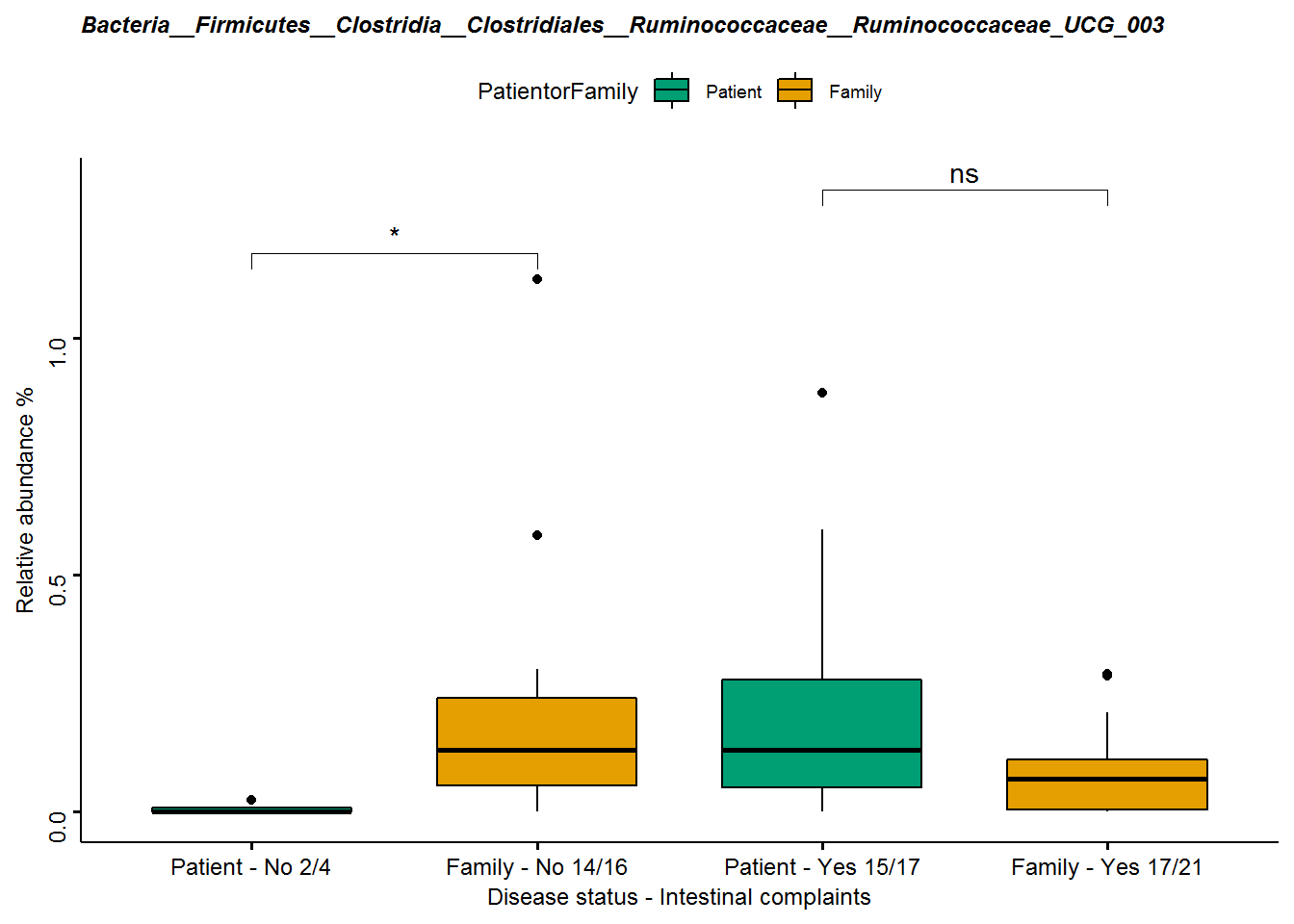

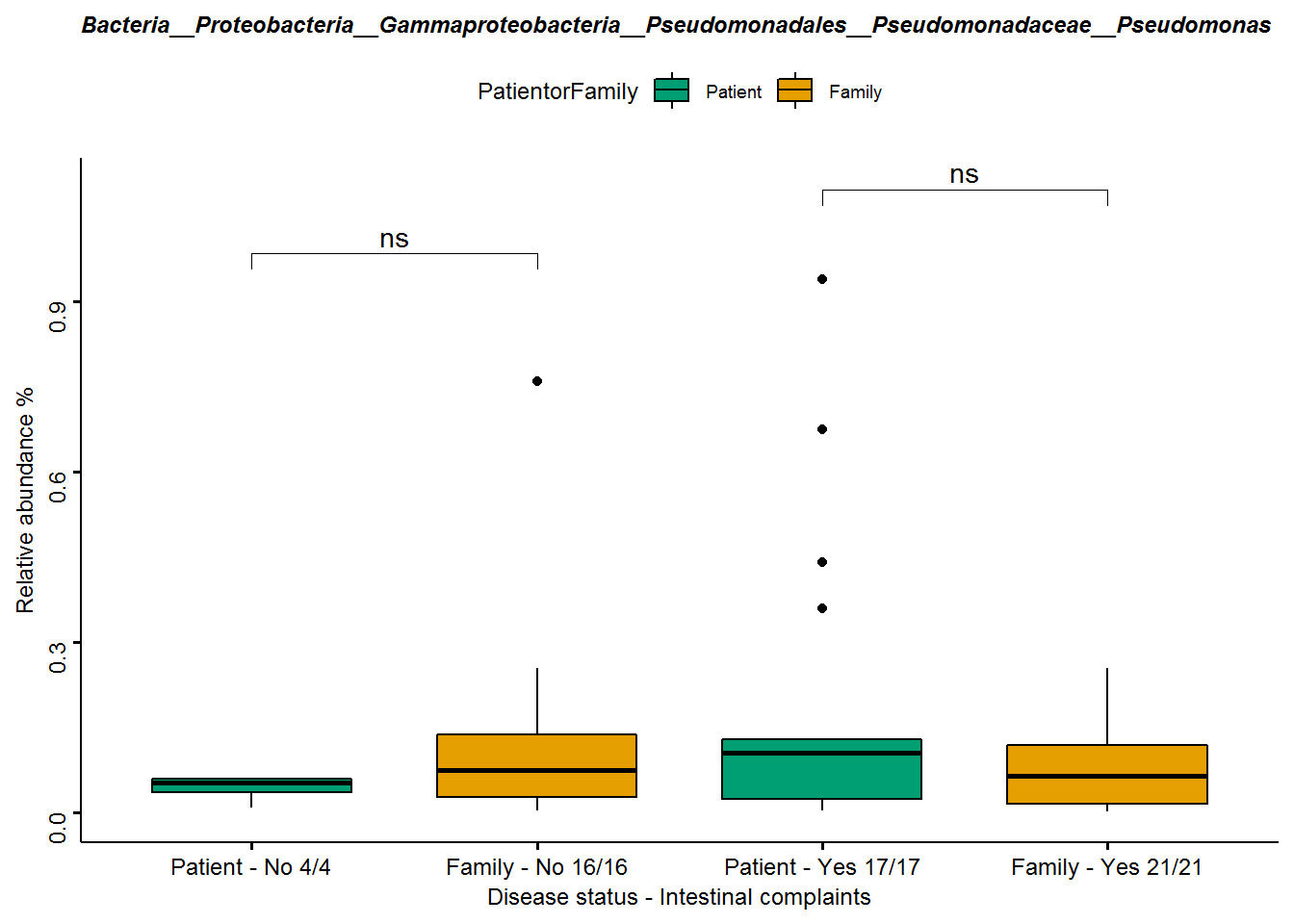

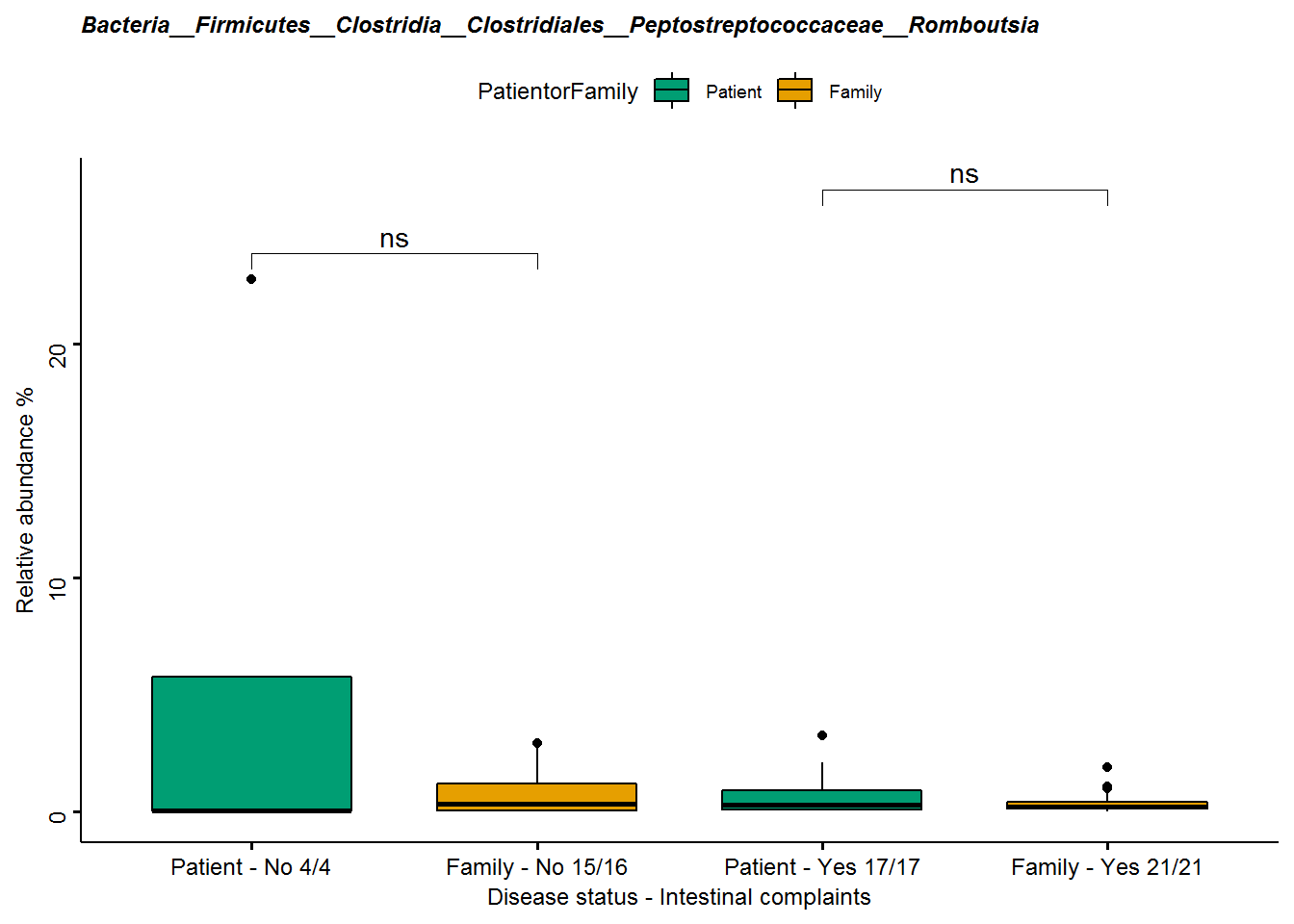

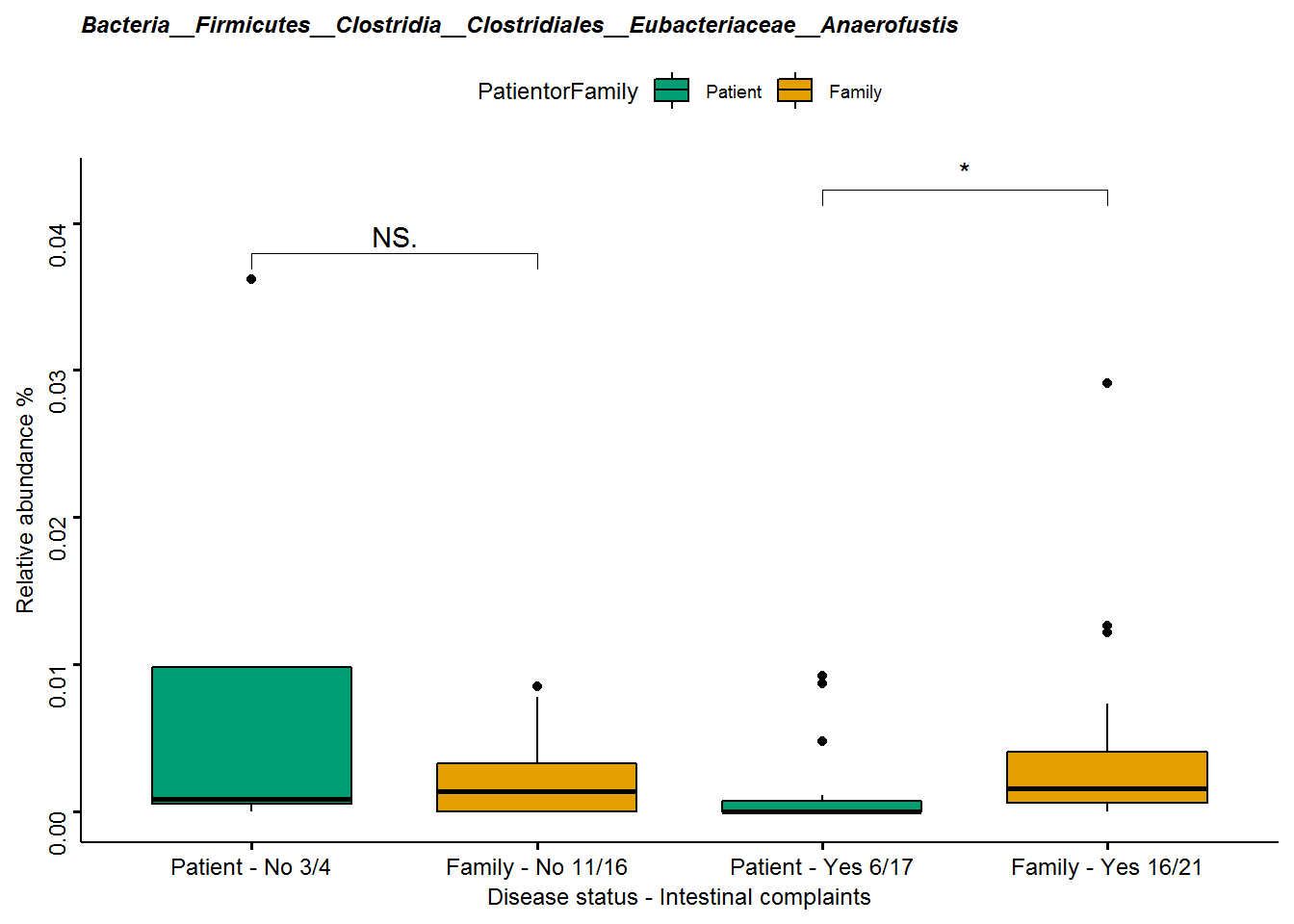

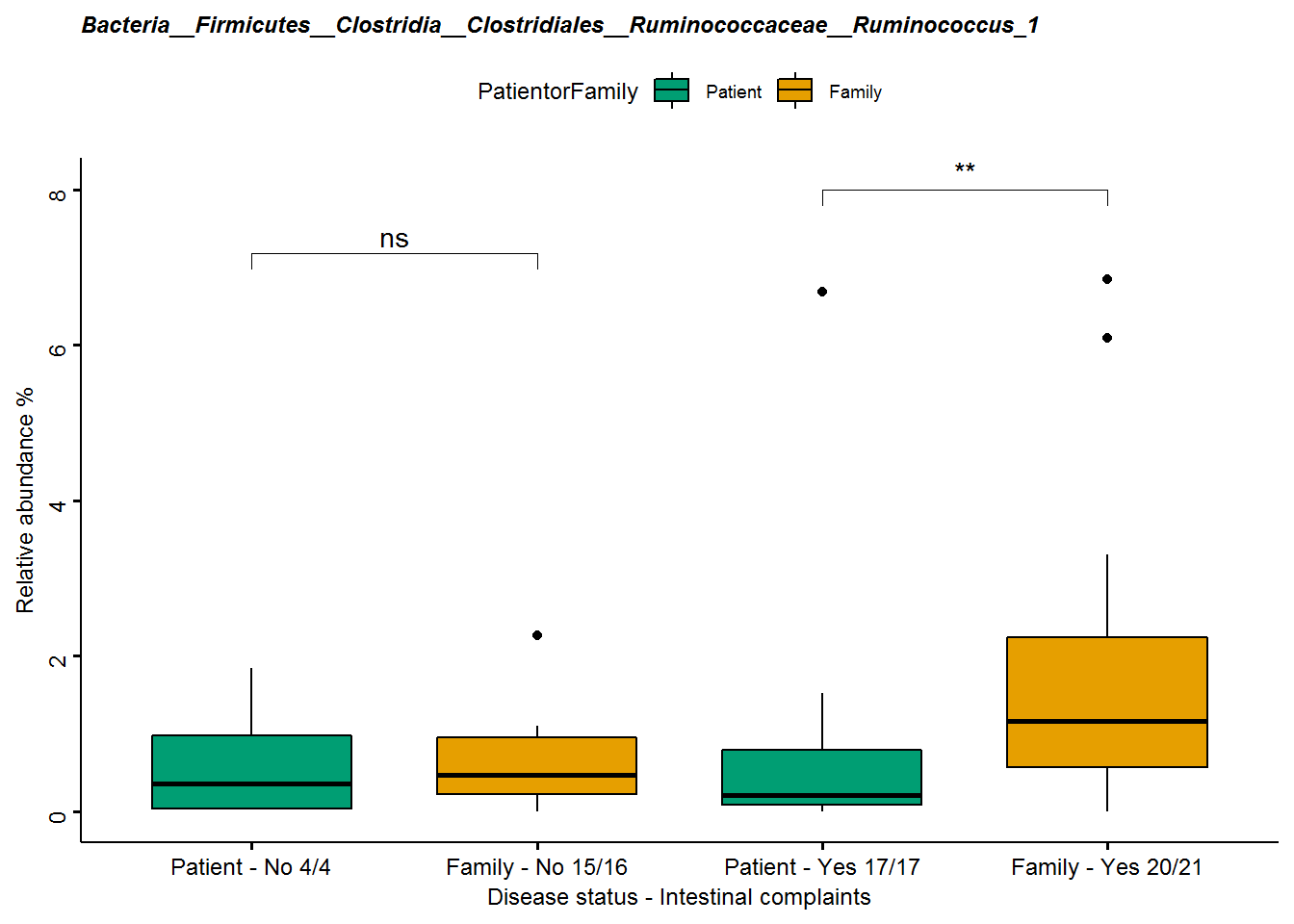

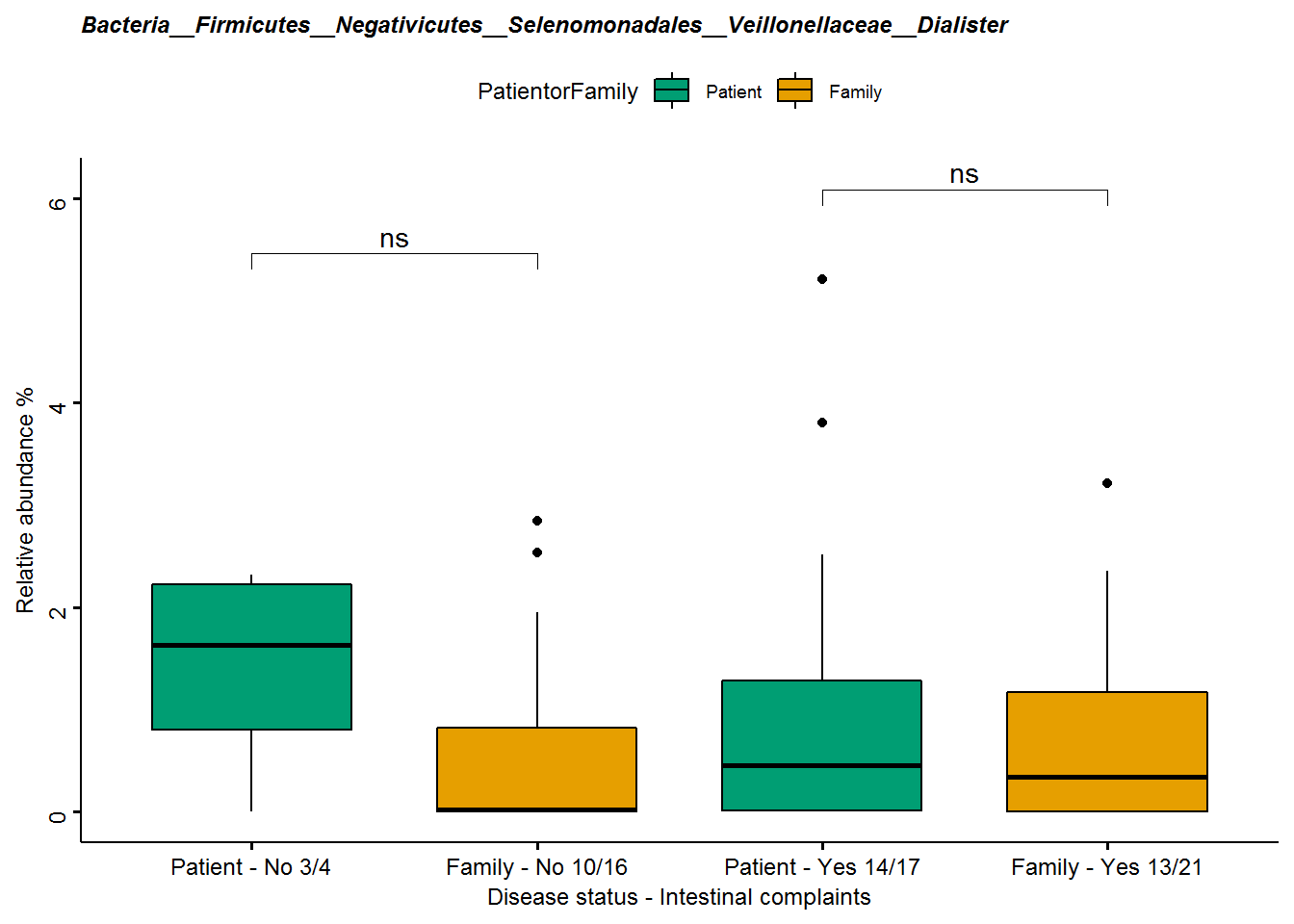

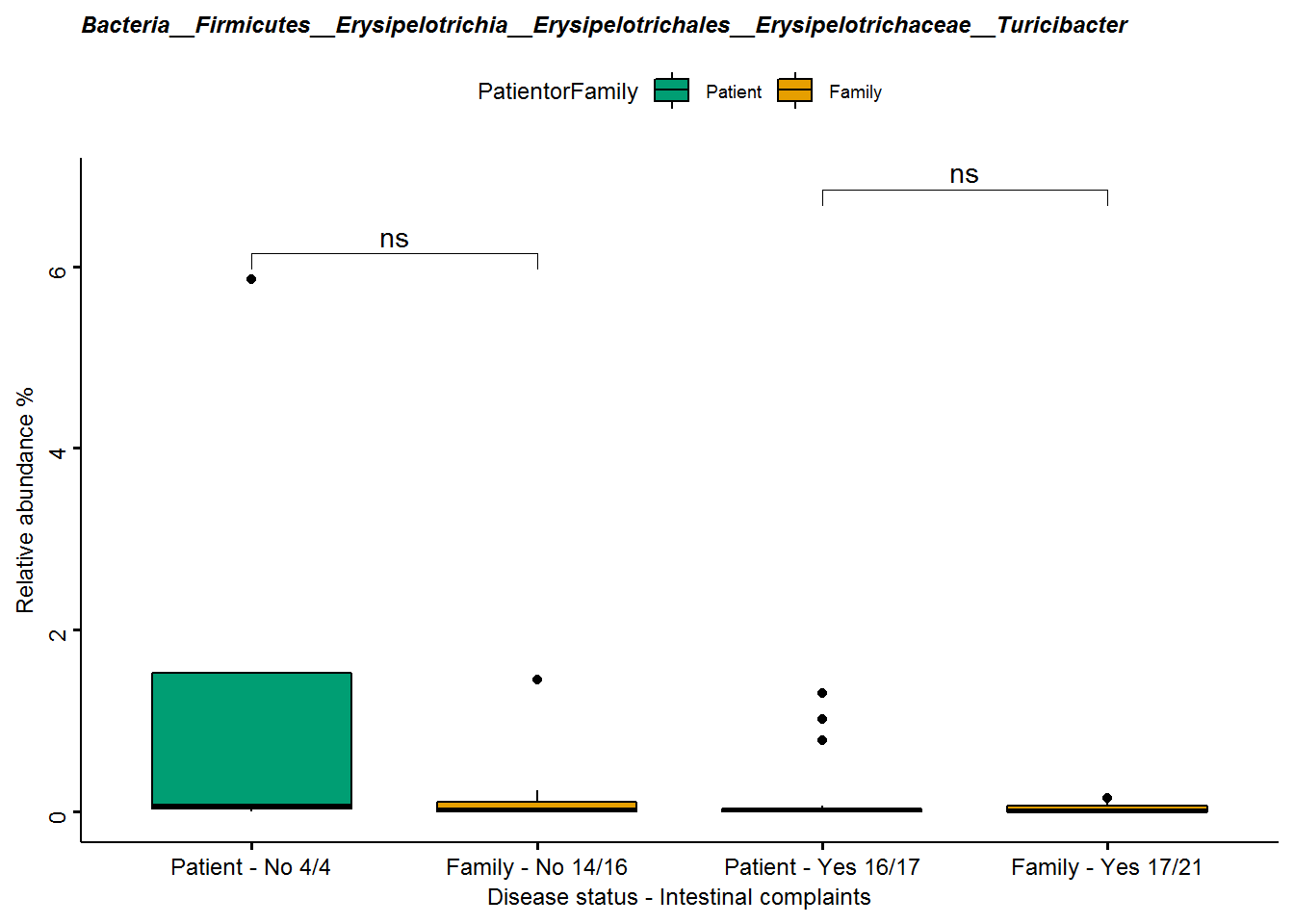

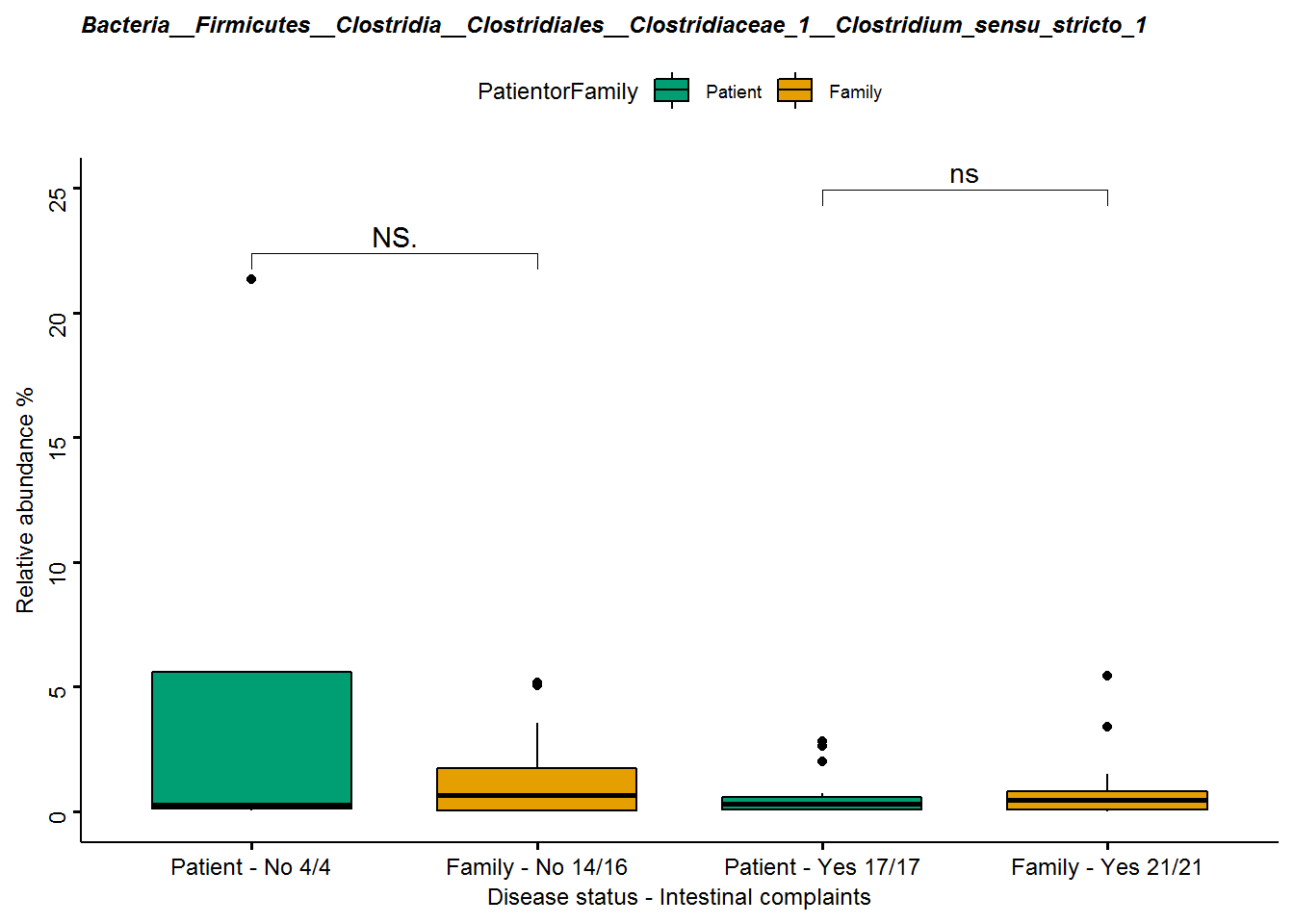

*Patients dataset

CBCL total score*

Relative abundance of seven genera associated FDR-uncorrected with CBCL total score. Three genera are part of the family *Lachnospiraceae* and one genus was found for other families namely *Ruminococcaceae, Eggerthellaceae, Bacteroidaceae* and *Desulfovibrionaceae*, see **Table S4** for statistical results per genus. Five genera*, Coprococcus 3, Desulfovibrionaceae – uncultured, Marvinbryantia, Moryella* and *Subdoligranulum*, showed an increase in relative abundance in patients with a higher CBCL total score, indicating more severe symptoms. Two genera, *Bacteroides* and *Eggerthella,* showed a decrease in patients with more severe symptoms, **see Figure S6**.

**Figure S6.** **Seven scatterplots showing the main effects of CBCL total problems score on the relative abundance of the nominal significant genera from the compositional analysis of the patients only dataset (n=22).** The blue lines represent a linear regression showing the direction of the effect.

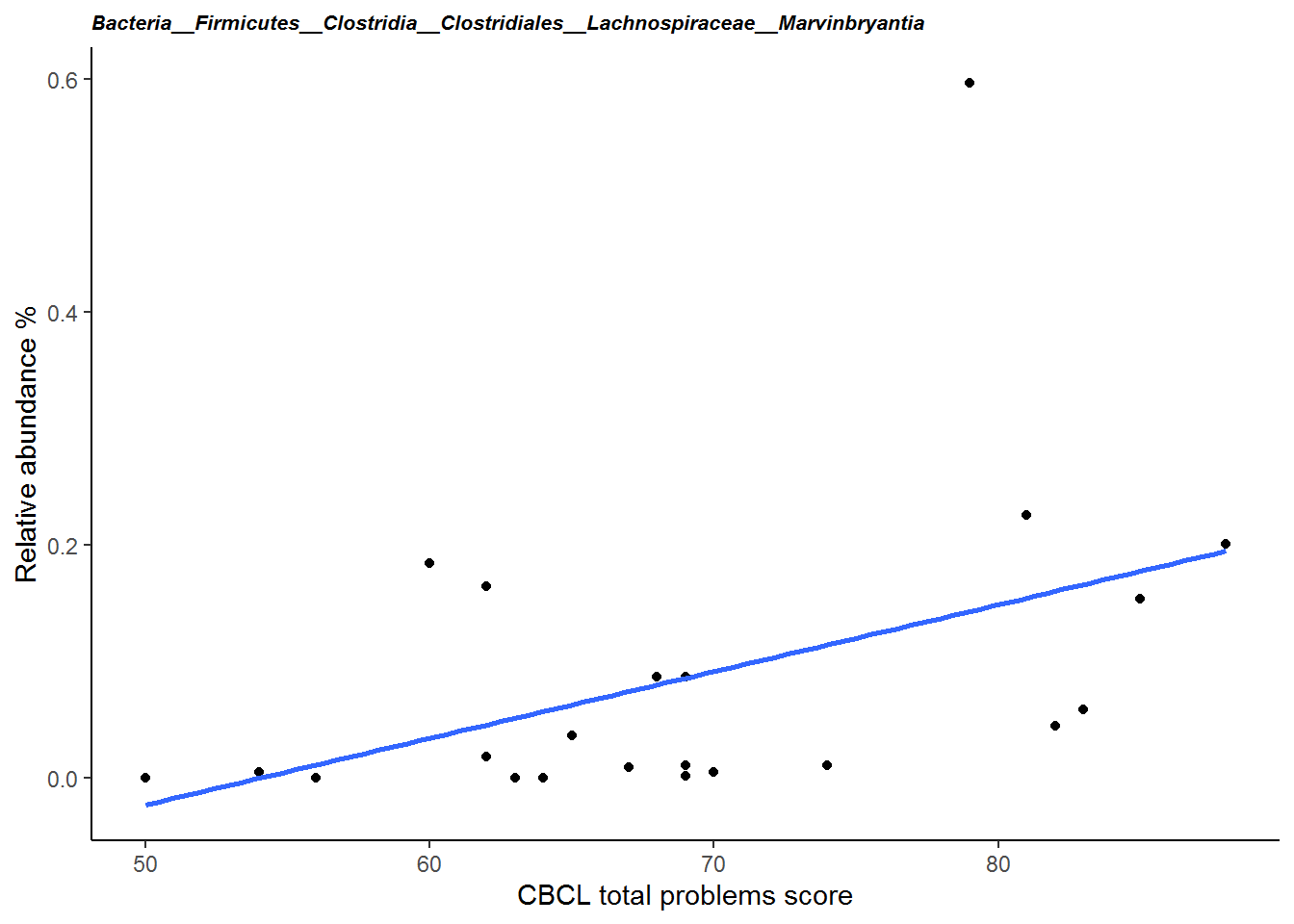

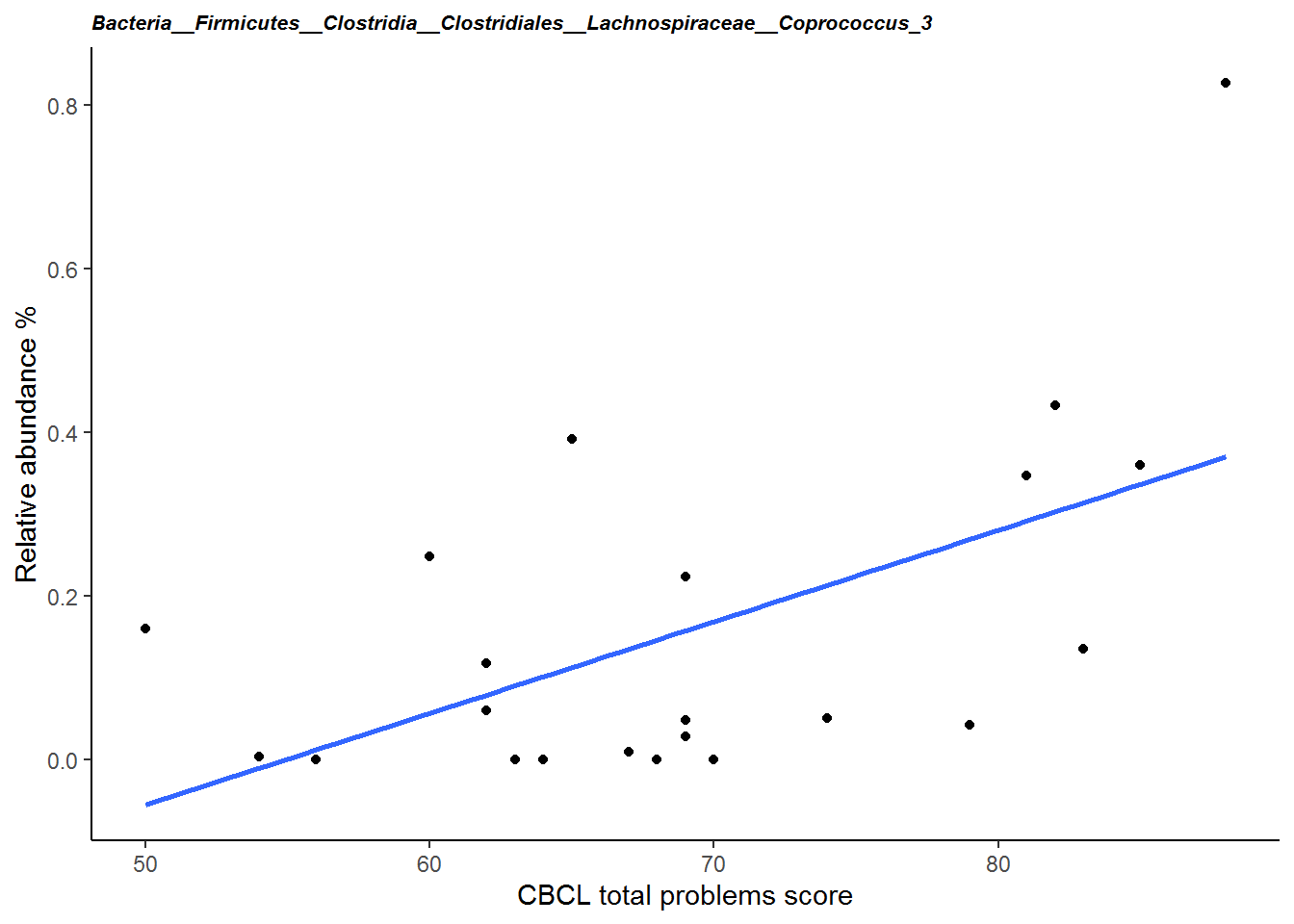

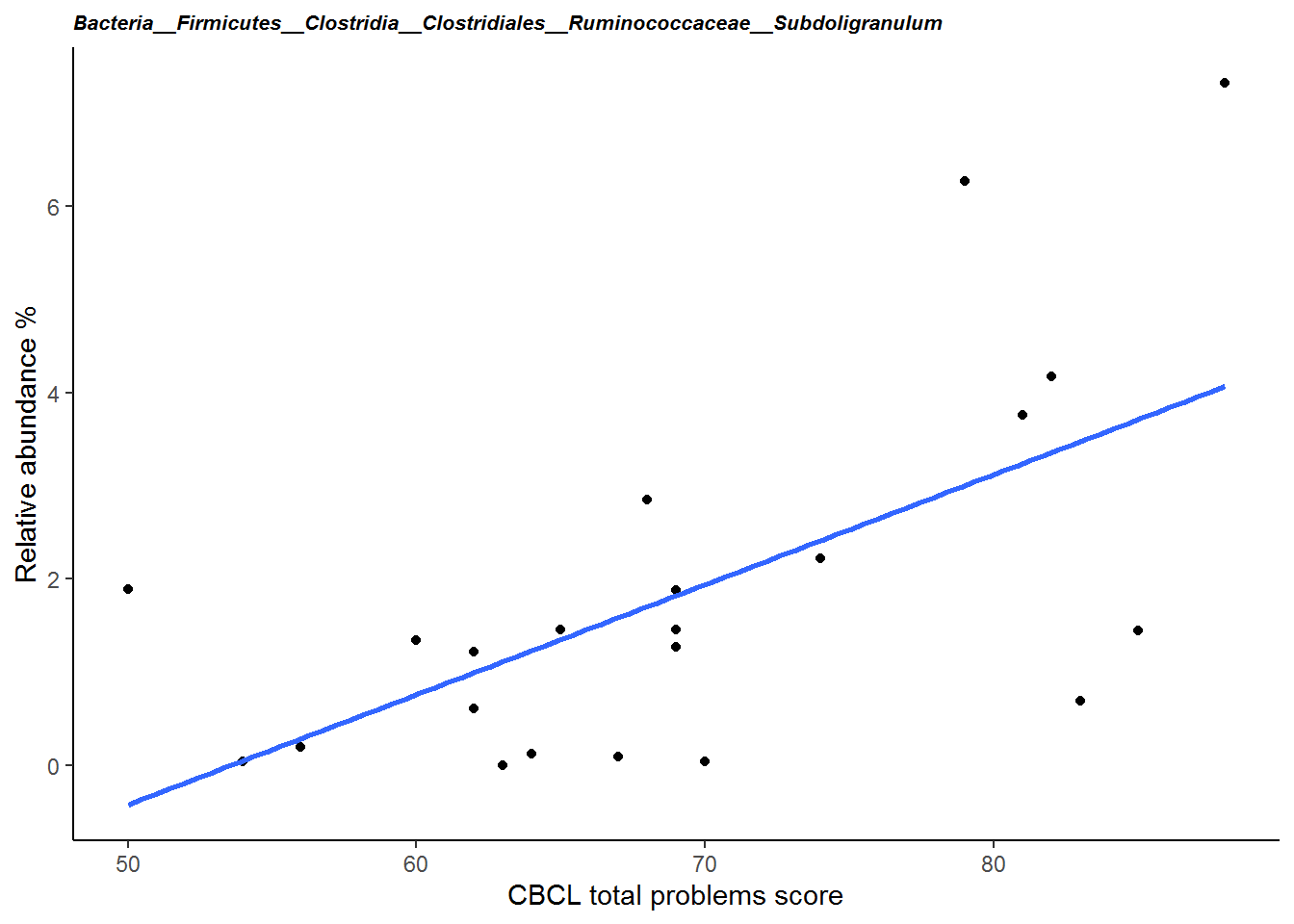

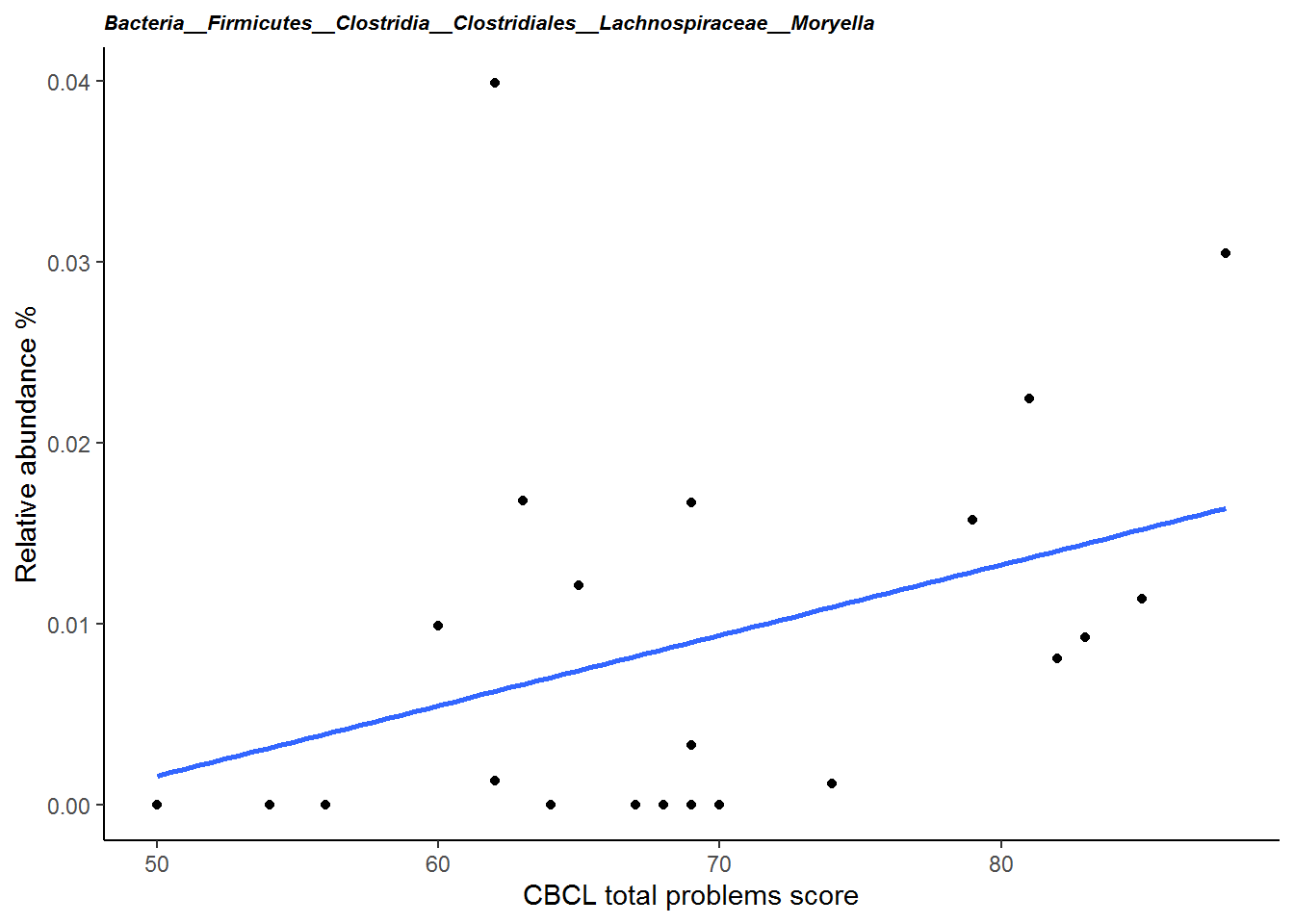

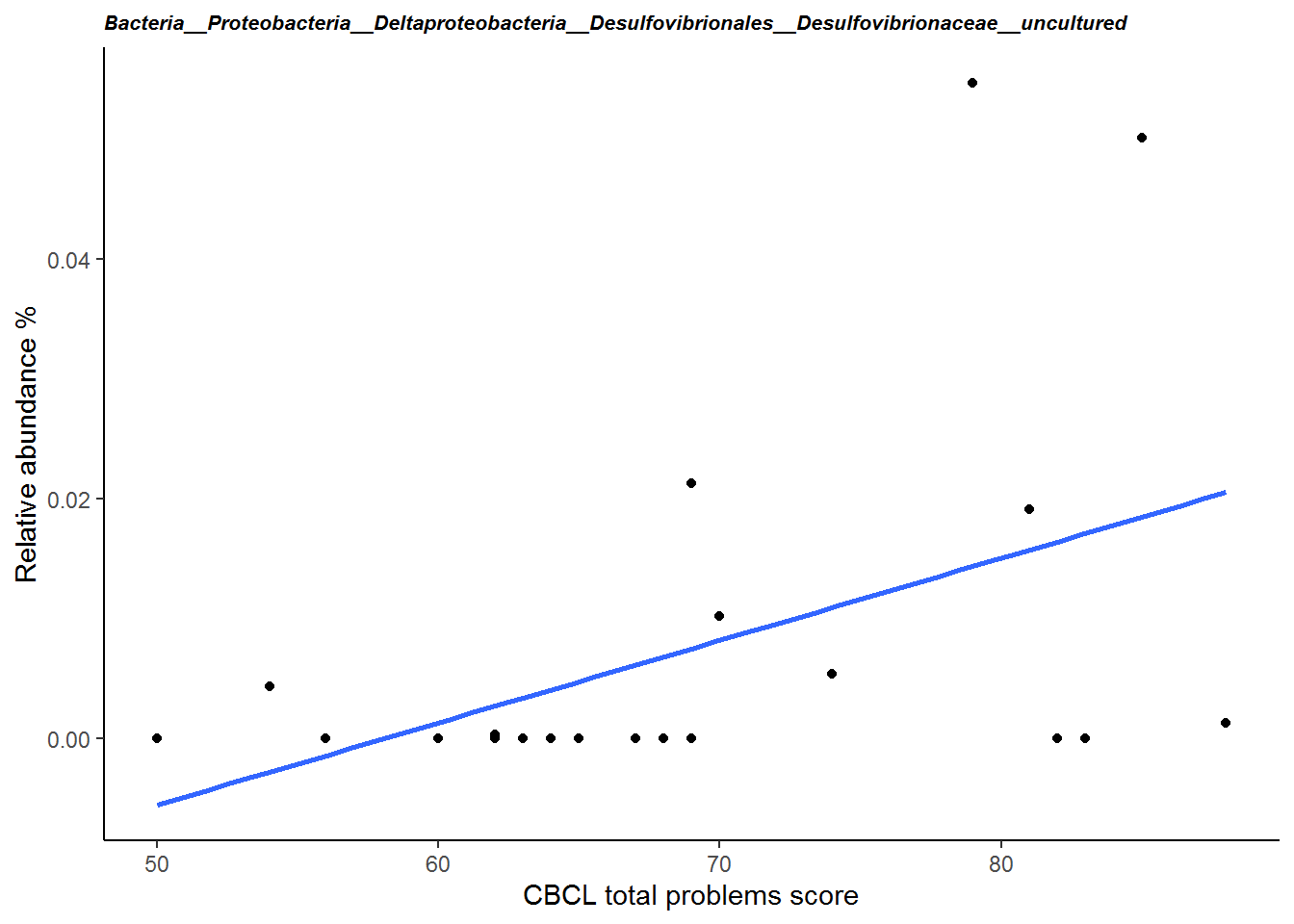

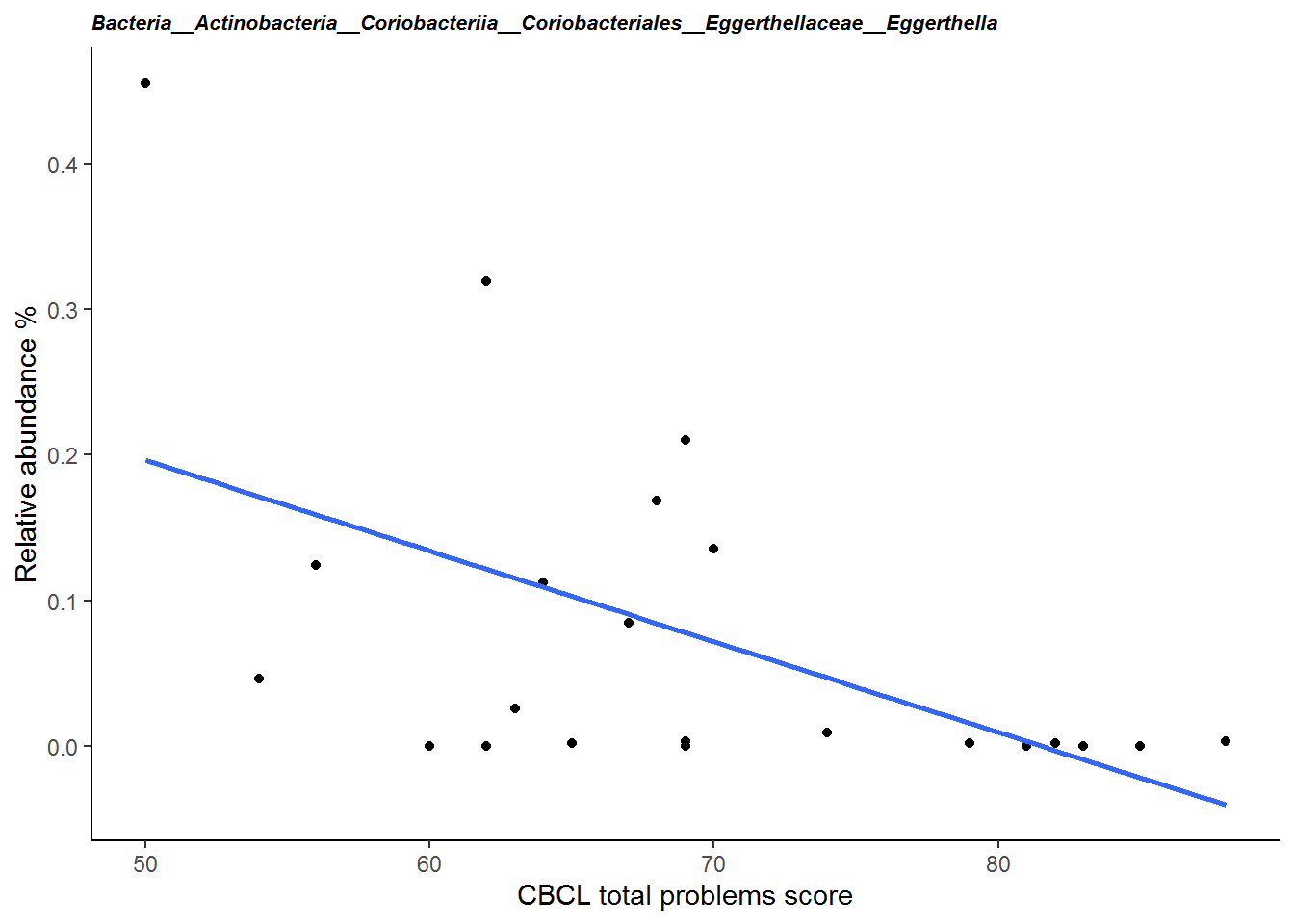

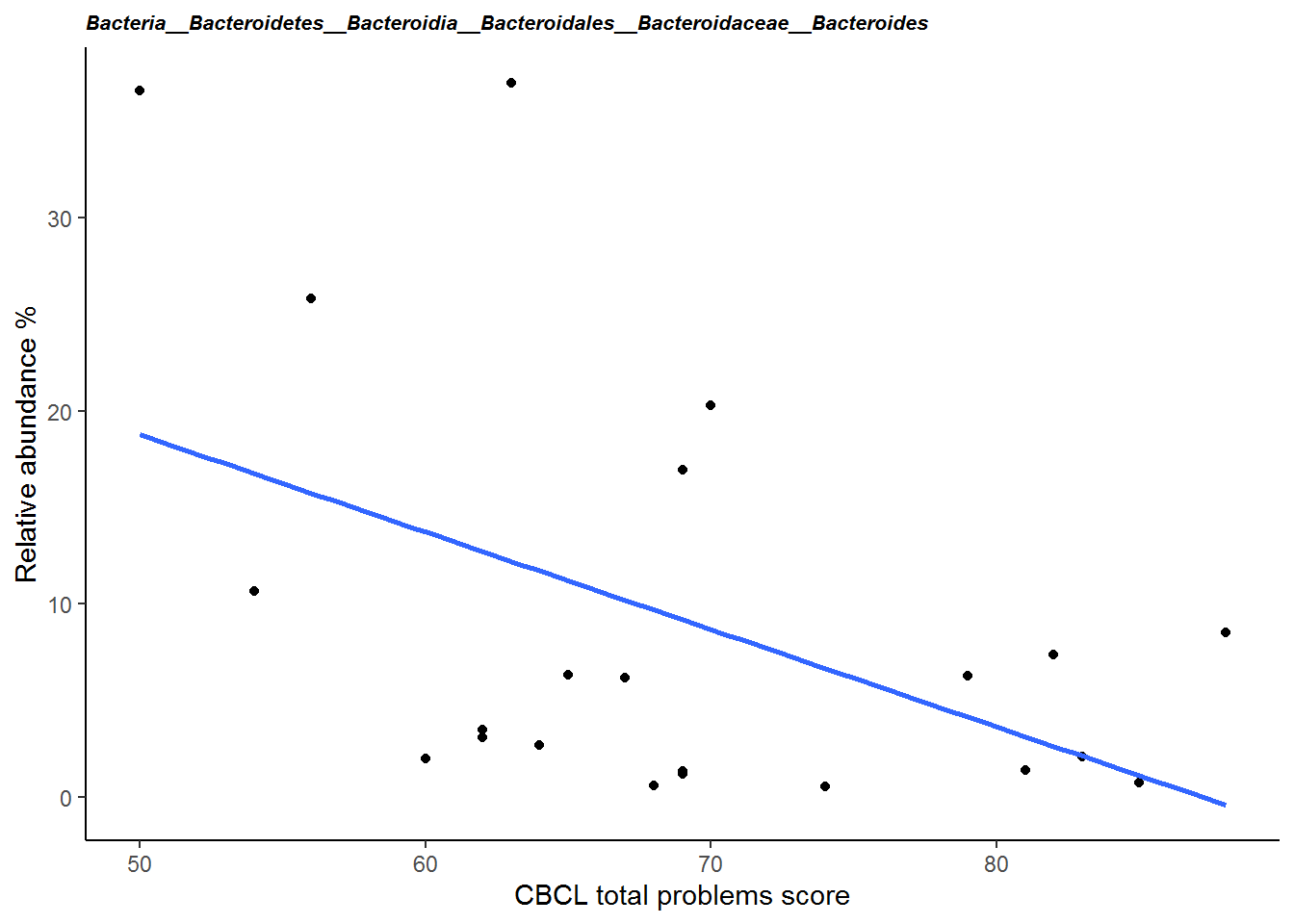

*ADOS comparative score*

Four genera, *Gemella, Atopobiaceae – uncultured, Butyricicoccus* and *Veillonella*, were found to be nominally significant associated with ADOS comparative score, see **Table S4** for statistical results. Genera *Gemella, Butyricicoccus* and *Veillonella* showed a decrease in relative abundance with an increase in symptom severity. This indicates a higher ADOS comparative score. Only one genus *Atopobiaceae – uncultured* showed an increase in patients with more severe symptoms, see **Figure S7**.

**Figure S7. Four scatterplots showing the main effects of ADOS comparative score on the relative abundance of the nominal significant genera from the compositional analysis of the patients only datasets (n=22).** The blue lines represent a linear regression showing the direction of the effect.

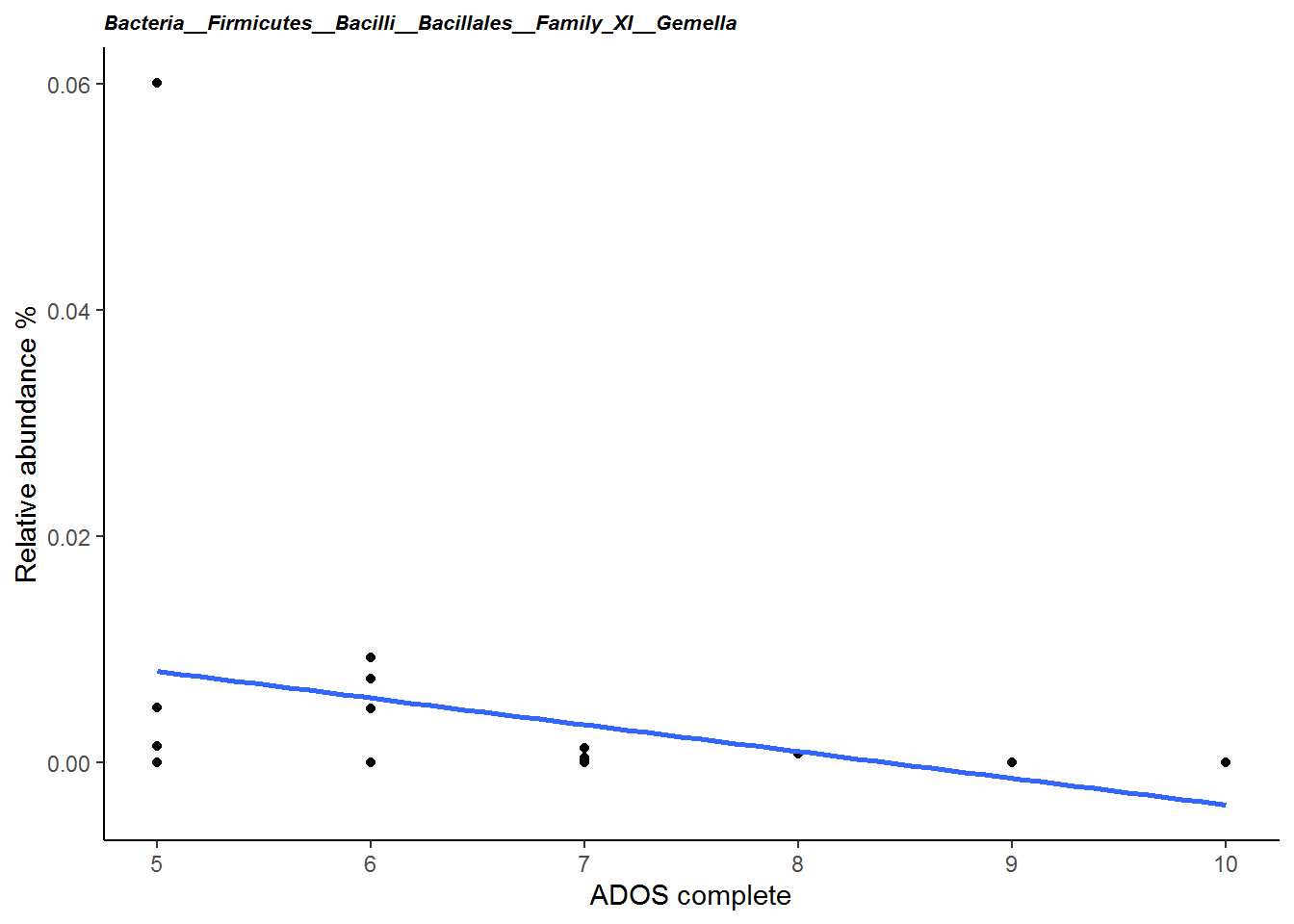

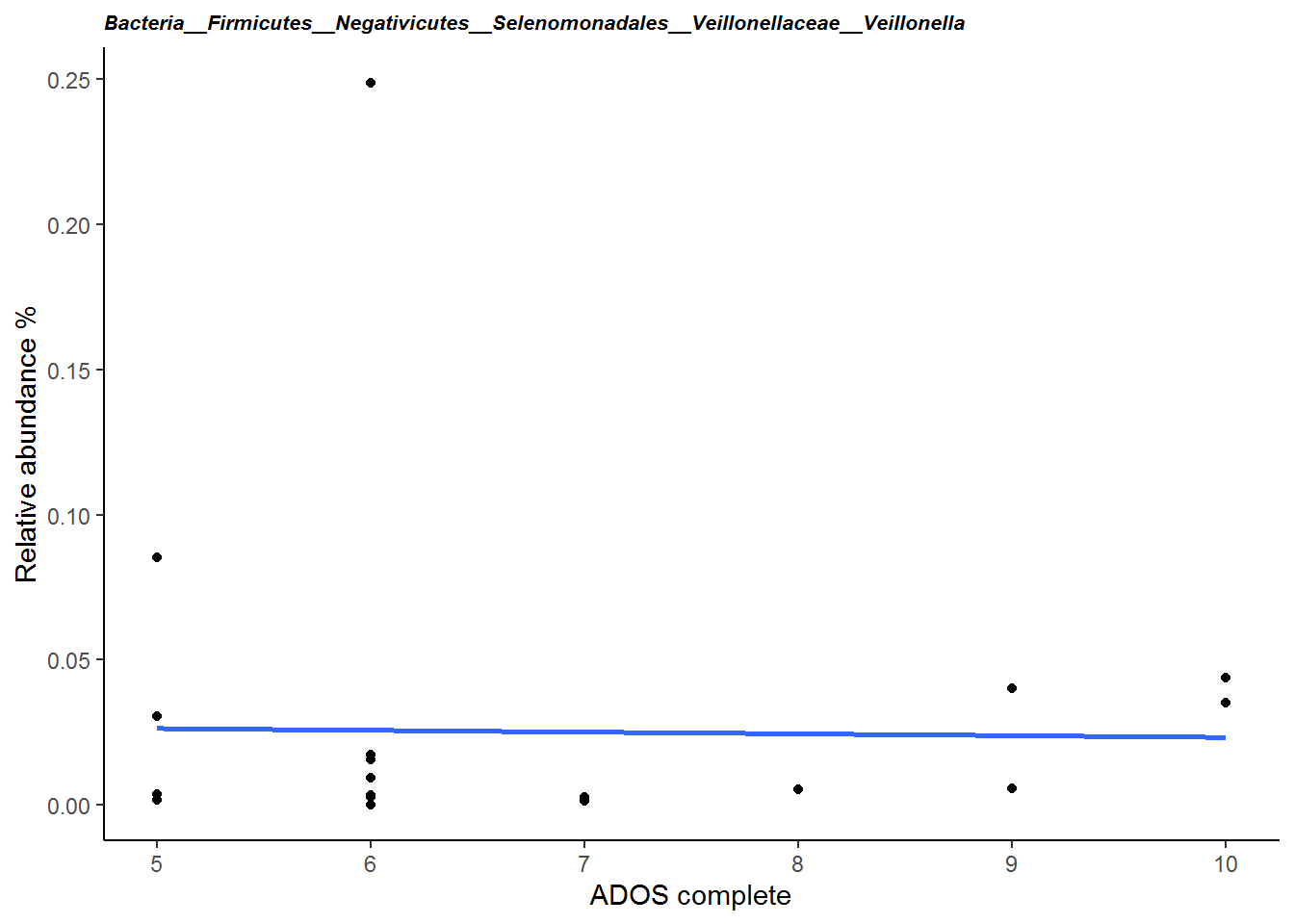

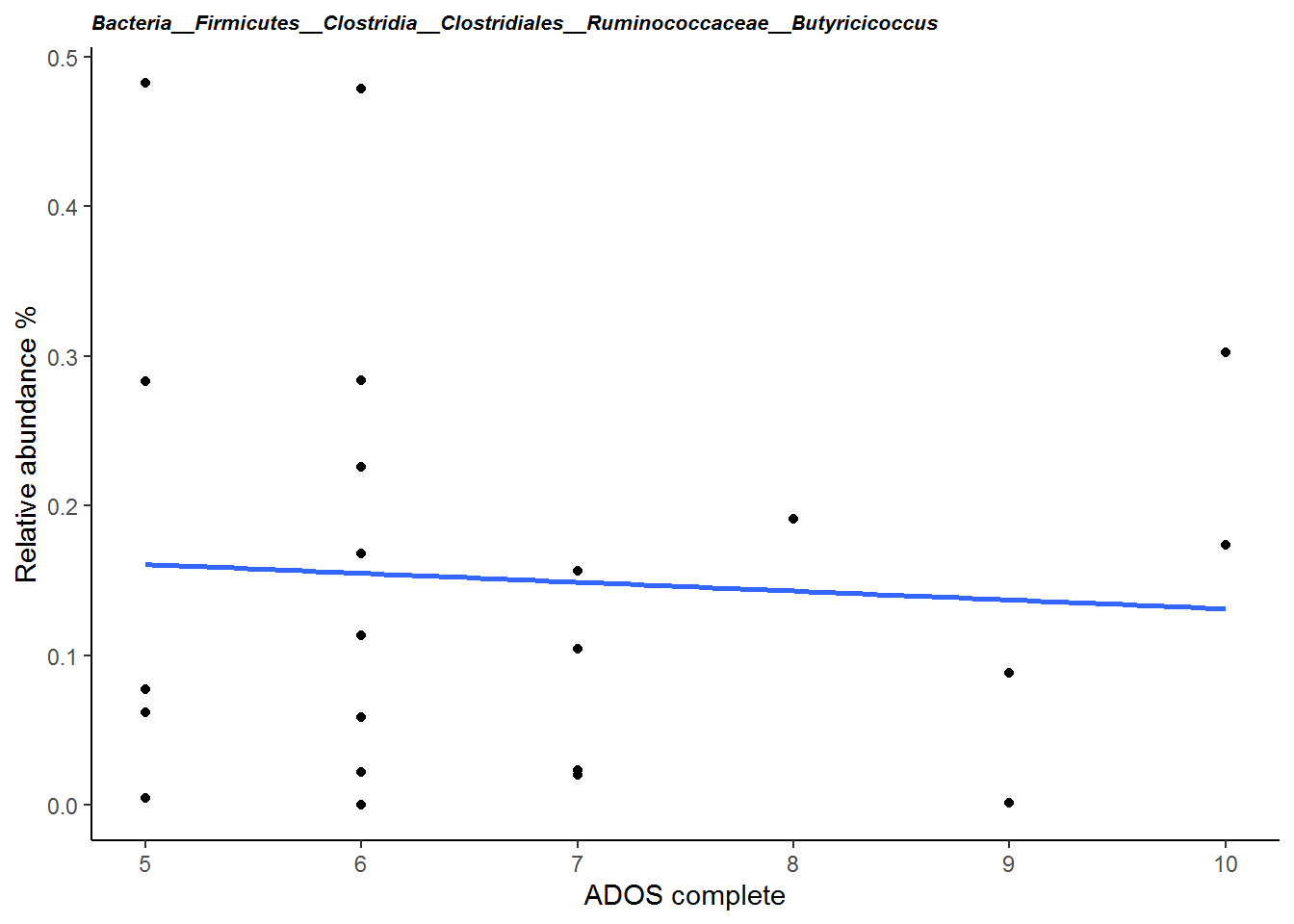

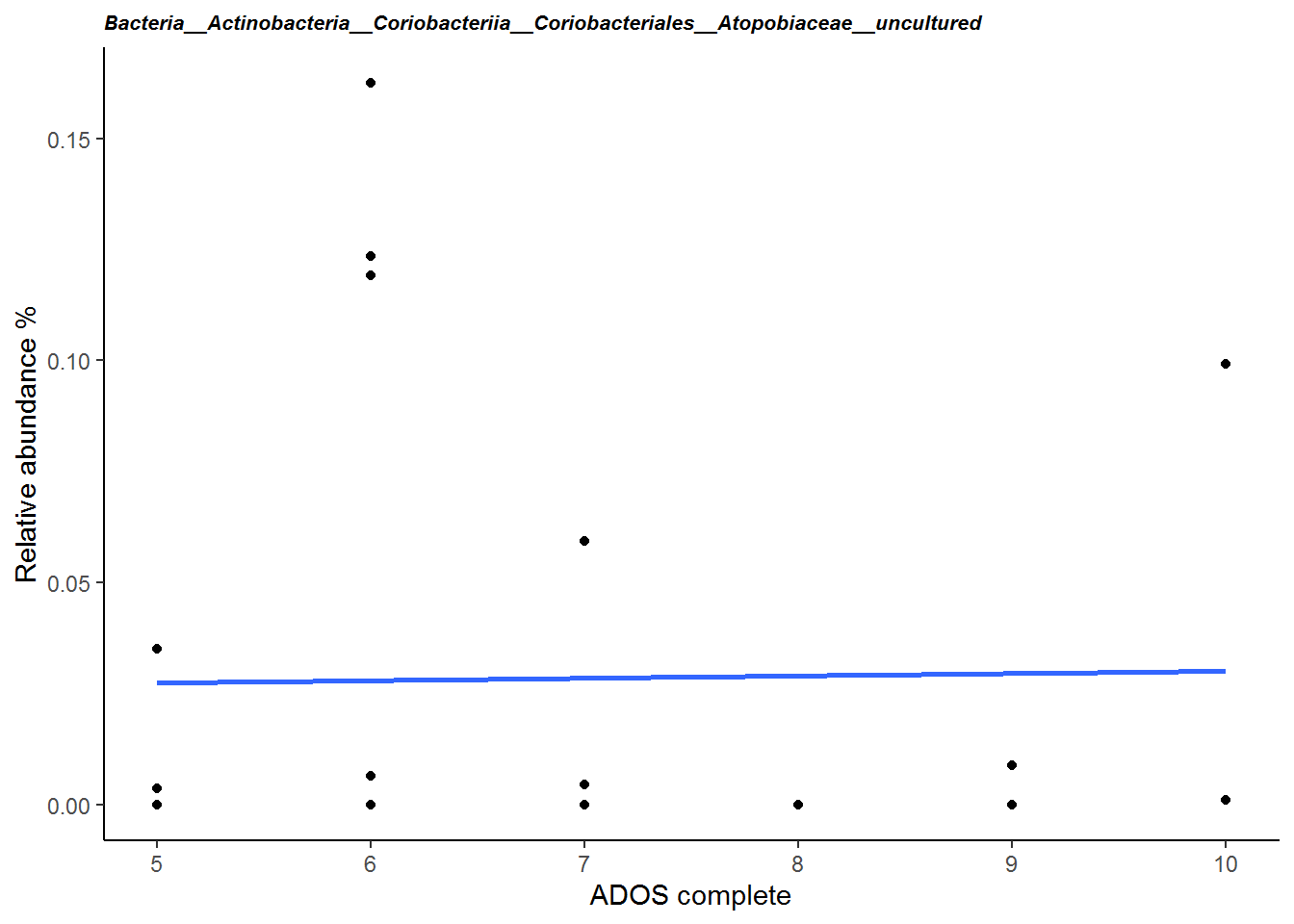

*Relation between disease-related symptom severity and genetic variants*

**Figure S8. Boxplot of CBCL total problems score grouped by intestinal complaints in the patients (yes/no).** The boxplot indicates the median (black horizontal line) and 25^th^ and 75^th^ quartiles as the outside of the box. The lines represent the largest and smallest values within the 1.5 interquartile range and the dots represents the outliers which is > 1.5 times and < 3 times the interquartile range.

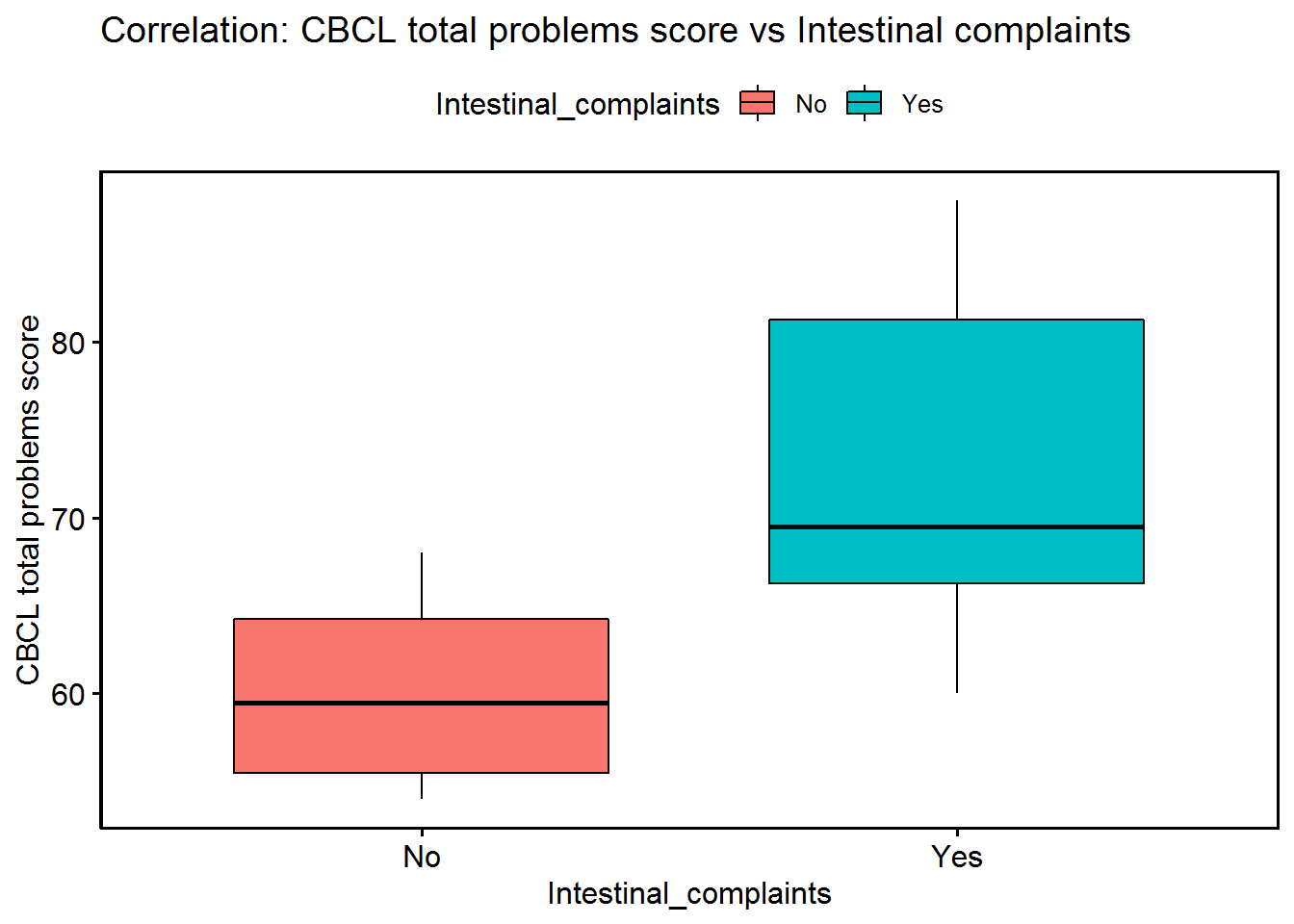

*Results overview*

**Table S4**. Statistical results from the separate models testing for effects of disease status, disease status*intestinal complaints, CBCL total score, ADOS comparative score and genetic variants.

| **Genus** | **Relative abundance %** | | **Disease status** | |  | **Disease status * Intestinal complaints** | | **CBCL total problems score** | | | **ADOS comparative score** | | | | **Literature** | |
| --- | --- | --- | --- | --- | --- | --- | --- | --- | --- | --- | --- | --- | --- | --- | --- | --- |
|  | Patient | Family | P-value | F-value | Direction | P-value | F-value | P-value | F-value | Direction | P-value | F-value | | Direction |  |  |
| *Phylum Actinobacteria* |  |  |  |  |  |  |  |  |  |  |  |  |  | |  |  |
| ***Family Atopobiaceae*** |  |  |  |  |  |  |  |  |  |  |  |  |  | |  |  |
| **uncultured** | 0.027 | 0.029 |  |  |  |  |  | **0.0003** | **21.141** | **+** | 0.0216 | 6.570 | + | |  |  |
| *Family Eggerthellaceae* |  |  |  |  |  |  |  |  |  |  |  |  |  | |  |  |
| Eggerthella | 0.074 | 0.069 |  |  |  |  |  | 0.0039 | 11.568 | - |  |  |  | | + ADHD  - ASD | Aarts et al., 2017,  Bungaard-Nielsen et al., 2020 |
| Gordonibacter | 0.045 | 0.021 |  |  |  | 0.0051 | 8.625 |  |  |  |  |  |  | |  |  |
| *Phylum Bacteroidetes* |  |  |  |  |  |  |  |  |  |  |  |  |  | |  |  |
| *Family Bacteroidaceae* |  |  |  |  |  |  |  |  |  |  |  |  |  | |  |  |
| Bacteroides | 8.791 | 8.506 |  |  |  | 0.0080 | 7.680 | 0.0328 | 5.531 | - |  |  |  | | - ADHD  + ASD  - Rett | Wang er al., 2020, Strati et al., 2016 |
| *Family Rikenellacaea* |  |  |  |  |  |  |  |  |  |  |  |  |  | |  |  |
| Alistipes | 0.601 | 0.871 | 0.0443 | 4.396 | - | 0.0181 | 5.993 |  |  |  |  |  |  | | + ADHD  - Rett | Wang et al., 2020, Strati et al., 2016 |
| *Phylum Firmicutes* |  |  |  |  |  |  |  |  |  |  |  |  |  | |  |  |
| *Family Clostridiaceae 1* |  |  |  |  |  |  |  |  |  |  |  |  |  | |  |  |
| Clostridium sensu stricto  1 | 1.442 | 0.95 |  |  |  | 0.0229 | 5.534 |  |  |  |  |  |  | |  |  |
| *Family Clostridiales vadinBB60 group* |  |  |  |  |  |  |  |  |  |  |  |  |  | |  |  |
| gut metagenome | 0.012 | 0.027 | 0.0333 | 4.963 | - |  |  |  |  |  |  |  |  | |  |  |
| uncultured organism | 0.117 | 0.178 | 0.0348 | 4.876 | - |  |  |  |  |  |  |  |  | |  |  |
| *Family Erysipelotrichaceae* |  |  |  |  |  |  |  |  |  |  |  |  |  | |  |  |
| Clostridium innocuum  Group | 0.005 | 0.007 |  |  |  | 0.0017 | 11.028 |  |  |  |  |  |  | |  |  |
| Holdemania | 0.005 | 0.006 |  |  |  | 0.0074 | 7.847 |  |  |  |  |  |  | |  |  |
| **Merdibacter** | 0.01 | 0.01 |  |  |  | **0.0003** | **15.313** |  |  |  |  |  |  | |  |  |
| Turicibacter | 0.404 | 0.077 |  |  |  | 0.0273 | 5.192 |  |  |  |  |  |  | | - ASD | Bundgaard-Nielsen et al., 2020 |
| *Family Eubacteriaceae* |  |  |  |  |  |  |  |  |  |  |  |  |  | |  |  |
| Anaerofustis | 0.0029 | 0.0032 | 0.0456 | 4.339 | - | 0.0230 | 5.526 |  |  |  |  |  |  | |  |  |
| *Family Family XI* |  |  |  |  |  |  |  |  |  |  |  |  |  | |  |  |
| Gemella | 0.004 | 0.002 |  |  |  |  |  |  |  |  | 0.0189 | 6.928 | - | |  |  |
| *Family Lachnospiraceae* |  |  |  |  |  |  |  |  |  |  |  |  |  | |  |  |
| Blautia | 3.683 | 6.304 | 0.0199 | 6.023 | - |  |  |  |  |  |  |  |  | | + Rett | Strati et al., 2016 |
| **Coprococcus 3** | 0.157 | 0.292 | **0.0003** | **16.303** | **-** |  |  | 0.0441 | 4.829 | + |  |  |  | | (Coprococcus - lower in ASD) | |
| Dorea | 0.67 | 0.948 | 0.0376 | 4.718 | - |  |  |  |  |  |  |  |  | | + ASD | Bundgaard-Nielsen et al., 2020 |
| Eisenbergiella | 0.045 | 0.014 |  |  |  | 0.0180 | 6.008 |  |  |  |  |  |  | |  |  |
| Eubacterium fissicatena  Group | 0.005 | 0.004 |  |  |  | 0.0091 | 7.414 |  |  |  |  |  |  | | (Eubacterium - lower in ASD) | |
| Eubacterium  xylanophilum group | 0.034 | 0.106 | 0.0095 | 7.643 | - |  |  |  |  |  |  |  |  | |  |  |
| Lachnospiraceae FCS020  Group | 0.064 | 0.131 | 0.0057 | 8.820 | - |  |  |  |  |  |  |  |  | |  |  |
| Lachnospiraceae  ND3007 group | 0.232 | 0.584 | 0.0023 | 11.035 | - |  |  |  |  |  |  |  |  | |  |  |
| Marvinbryantia | 0.09 | 0.159 |  |  |  |  |  | 0.0252 | 6.183 | + |  |  |  | |  |  |
| Moryella | 0.009 | 0.016 | 0.0153 | 6.594 | - |  |  | 0.0497 | 4.558 | + |  |  |  | | Murine ASD model | de Theije et al., 2014 |
| uncultured | 0.061 | 0.114 | 0.0382 | 4.685 | - |  |  |  |  |  |  |  |  | |  |  |
| *Family Peptococcaceae* |  |  |  |  |  |  |  |  |  |  |  |  |  | |  |  |
| Peptococcus | 0.056 | 0.018 | 0.0264 | 5.435 | + |  |  |  |  |  |  |  |  | |  |  |
| Romboutsia | 1.47 | 0.554 |  |  |  | 0.0081 | 7.656 |  |  |  |  |  |  | | - ASD | Bundgaard-Nielsen et al., 2020 |
| *Family Ruminococcaceae* |  |  |  |  |  |  |  |  |  |  |  |  |  | |  |  |
| Acetanaerobacterium | 0.002 | 0.002 |  |  |  | 0.0481 | 4.117 |  |  |  |  |  |  | | + ASD | Bundgaard-Nielsen et al., 2020 |
| Butyricicoccus | 0.148 | 0.187 | 0.0103 | 7.467 | - |  |  |  |  |  | 0.0302 | 5.727 | - | | - Rett | Strati et al., 2016 |
| Ruminococcaceae  UCG003 | 0.156 | 0.144 |  |  |  | 0.0085 | 7.555 |  |  |  |  |  |  | |  |  |
| Ruminococcus 1 | 0.752 | 1.179 | 0.0108 | 7.361 | - | 0.0343 | 4.755 |  |  |  |  |  |  | | (Ruminococcus - higher and lower in ASD, lower in Rett) | Strati et al., 2016 |
| Subdoligranulum | 1.826 | 4.682 |  |  |  |  |  | 0.0016 | 14.755 | + |  |  |  | | - ASD | Bundgaard-Nielsen et al. 2020 |
| UBA1819 | 0.063 | 0.039 |  |  |  | 0.0313 | 4.927 |  |  |  |  |  |  | |  |  |
| uncultured | 0.189 | 0.453 | 0.0273 | 5.369 | - |  |  |  |  |  |  |  |  | |  |  |
| *Family Veillonellaceae* |  |  |  |  |  |  |  |  |  |  |  |  |  | |  |  |
| Dialister | 1.047 | 0.708 |  |  |  | 0.0239 | 5.448 |  |  |  |  |  |  | | - ADHD  - ASD  - Rett | Jiang et al., 2018, Strati et al. 2016 |
| Veillonella | 0.024 | 0.031 |  |  |  |  |  |  |  |  | 0.0439 | 4.841 | - | | + ADHD  - ASD - Down Syndrome | Akram, 2017, Wang et al. 2020, Biagi et al., 2014 |
| *Phylum Proteobacteria* |  |  |  |  |  |  |  |  |  |  |  |  |  | |  |  |
| *Family Desulfovibrionaceae* |  |  |  |  |  |  |  |  |  |  |  |  |  | |  |  |
| uncultured | 0.011 | 0.014 |  |  |  |  |  | 0.0500 | 4.543 | + |  |  |  | |  |  |
| *Family Pseudomonadaceae* |  |  |  |  |  |  |  |  |  |  |  |  |  | |  |  |
| Pseudomonas | 0.157 | 0.095 |  |  |  | 0.0362 | 4.649 |  |  |  |  |  |  | | + ASD | Bundgaard-Nielsen et al. 2020 |

**Disease status:** - or + coding indicates lower or higher relative abundance in patients compared to their family members. **Associations within the patient group:** + indicates increasing relative abundance with more severe behavioral symptoms, - marks decreasing relative abundance of the patients as a function of increasing behavioral symptom. For the interaction between disease status and intestinal complaints no color coding was used, see **Figure S5** for the direction of the effects. **Literature:** per genus, relevant literature on NDDs or IDDs is listed with (where possible) the direction of the effect of this genus in these studies. A result in **bold** indicates a significant effect after correction for multiple testing (FDR).

**Supplementary Discussion**

*Compositional differences in Kleefstra syndrome – FDR-uncorrected significant*

Across the models we tested in this study, some genera overlap between these models and thereby, despite not being FDR-corrected significant, show their potential relevance in KS. Moreover, some of our FDR-uncorrected findings interestingly overlap with other relevant NDDs, see **Table S4**, column Literature. We feel it is relevant to share these as this is the first study into gut microbiome in KS. These uncorrected results may serve as selection short-list for future studies in KS.
For example, abundance of genus *Bacteroides* is lower in patients compared to family members when reporting intestinal complaints and at the same time, lower abundance within patients relates to more severe behavioral problems (on the CBCL). This complementary pattern indicates *Bacteroides* being a relevant bacterium in the KS pathology. Moreover, this genus is often related with neurodevelopmental and mental disorders; increased in abundance in ASD in several studies (Bundgaard-Nielsen et al., 2020), increased in RS (Borghi & Vignoli, 2019) and in patients in a first psychotic episode (Schwarz et al., 2018), as well as gastrointestinal syndromes (Tamana et al., 2021; Wang et al., 2020; Yang et al., 2020). This may hence be a relevant target for potential interventions aiming at improved MGBA functioning. A recent study showed such effect; *Bacteroides uniformis* levels were increased after fecal microbiota transfer in patients with metabolic syndrome, interestingly positively associating with dopamine transporter binding in the brain (Hartstra et al., 2020).
Several other genera observed in this study have been associated with NDDs: *Moryella* was associated with serotonin levels in a murine model for autism (de Theije et al., 2014). In the current study this genus showed lower abundance in patients compared to family members and associated positively with CBCL (i.e. higher abundance associated with more severe behavioral problem scores).
Genus *Alistipes* was found numerically higher in ADHD patients compared to controls (Wang et al., 2020) and higher in non-responders of ketogenic diet in children with epilepsy (Zhang et al., 2018). KS patients showed lower abundance *Alistipes* versus family members, but only in those subjects reporting intestinal complaints.
*Lachnospiraceae (group)* genera and *Ruminococcaceae,* in the current study found lower in KS patients, are also observed lower in patients in their first psychotic episode and in Rett syndrome patients (Borghi et al., 2017; Schwarz et al., 2018).  *Eggerthella* was increased in Rett syndrome patients (Borghi et al., 2017) and found relating negatively with CBCL scores in our study.
In a small sample of patients with Down syndrome (n=17 patients versus n=16 family members), decreased *Veillonella* abundance was found (Biagi et al., 2014). In our study *Veillonella* abundance negatively associated with ADOS scores.

*ADOS versus CBCL associations with gut microbiota*

In the associations with ADOS, none were FDR-significant and correlation plots with relative abundance do not show convincing patterns, possibly partly due to the lower sample size available for this analysis. This difference compared to CBCL is a relevant point of attention for future studies, as ADOS is a specific measure for autistic symptoms assessed by a professional in a 30-60 minutes during observation. It is hence a more independent measure focused on autistic behaviours compared to CBCL, in which the occurrence of a wider range of behavioural symptoms (cognitive, emotional, physical and social) during the past 6 months is rated by a representative of the patient. Consequently, ADOS and CBCL measure different aspects of behaviour and daily functioning, which is reflected in the absence of correlation between these symptoms scores. This also seems to be reflected in the non-overlapping relations with the gut microbiota of KS patients. Possibly, despite the fact many patients meet the diagnostic criteria of ASD, in the context of the gut microbiota, the wider range of behavioural symptoms assessed by the CBCL is a more relevant measure.

*Associations between gut microbiota and behavioural symptoms versus intestinal complaints*
Both behavioral symptoms and intestinal complaints are frequent in KS. Moreover, behavioral symptoms were higher in patients reporting intestinal complaints in the current dataset. Despite this phenotypic relation, the association between the gut microbiota was more consistently related to behavioral symptom severity (associations both with community and composition) compared to intestinal complaints (associations only in composition) even though intestinal complaints are intuitively more closely linked to the gut microbiota. A study by Kang et al. (2013) showed a similar effect, where autistic symptoms were, but gastrointestinal symptoms were not related with less diverse microbiota. A complicating factor here is that due to limited verbal abilities in NDDs patients may express intestinal complaints non-verbally through problematic behaviors. These associations between the microbiota and disease-associated symptom severity (such as intestinal complaints and behavioral symptoms) suggest gut microbial and MGBA alterations are part of the KS phenotype.

*Blasting results*

Blasting of the most frequent observed sequence classified into *Coprococcus 3* resulted in 100 mostly unclassified 100% identical hits. Of these, a Coprococcus sp. (GenBank: LC333659.1, Otu060) was found in an unpublished sample of preterm neonates, in the context of delayed gut microbial colonization preventing necrotizing enterocolitis (NEC) via epigenetic regulation of immune-metabolic pathways. The Uncultured Coprococcus sp. clone R102.P1.UK_96645 was found in a study assessing effects of the gut microbiota on the pharmacodynamics of the diabetes medication metformin (Kim et al., 2021).
Blasting of the most frequent observed sequence classified into Atopobiaceae – uncultured resulted in two hits which were 100% identical and had the lowest E-value (4e-122). Of these, an Uncultured Bacterium clone e5ca (GenBank: MT54672.1) was found in a study assessing the role of gut microbiota in the symptom severity in Schizophrenia where the bacteria in phylum Actinobacteria were increased in patients with Schizophrenia (Li et al., 2020). The Uncultured Bacterium clone dnOTU_387 (GenBank: MN211939.1) was found as an direct submission sample.
Blasting of the most frequent observed sequence classified into Merdibacter resulted in 8 hits which were 100% identical and had the lowest E-value (1e-121). Of these, Clostridiales bacterium 10-3b (GenBank: HQ452860.1) is described as butyrate-producing bacteria in the caecum of chickens (Eeckhaut et al., 2011). An Uncultured bacterium clone RL246_aai76e06 (GenBank: DQ793739.1) was found in a study assessing gut microbiota in obesity where obese participants had more Firmicutes bacteria compared to lean control. After dieting the abundance of Firmicutes decreased (Ley, Turnbaugh, Klein, & Gordon, 2006). An Uncultured bacterium clone YO00080F06 (GenBank: EU197712.1) was found in a study assessing gut microbiome in recurrent Clostridium difficile induced antibiotic-associated diarrhea where patients had a decreased overall microbiome diversity and an altered distribution where the majority of bacteria were not part of the phyla Bacteriodetes and Firmicutes (Chang et al., 2008). Another Uncultured bacterium clone H08UC9050 (GenBank: HM807201.1) was found in a twin study assessing the interaction between the microbiota and mucosa of patients with ulcerative colitis where Firmicutes was visually less represented in patients with UC (Lepage et al., 2011).
